# Lipid-induced Caveolin1-Lipid droplet trafficking is associated with lipid droplet growth

**DOI:** 10.64898/2025.12.10.693432

**Authors:** Esther Ocket, Tamara Pietrucik, Luis Wong-Dilworth, Michaela Rath, Daniel Aik, Dmytro Puchkov, Max Ruwolt, Martin Lehmann, Claudia Matthaeus

## Abstract

Caveolae are 50-100 nm sized plasma membrane invaginations involved in cellular lipid uptake and lipid accumulation. Several caveolin1 and cavin1 mutations are associated to human lipodystrophy. Interestingly, previous research identified caveolae proteins at lipid droplets although their specific cellular function at lipid droplets and in lipid trafficking is not understood. Here, we show that extracellular dietary lipids like oleic acid and cholesterol shift caveolae at the plasma membrane to highly spherical invaginations. Within 3 hours of lipid treatment caveolin1 accumulates specifically to lipid droplets. Correlative fluorescence FIB-SEM and split-APEX proteomics revealed no caveolar vesicles at the lipid droplet but caveolin1 alone. Mechanistically, caveolin1-lipid droplet trafficking is regulated by EHD2 at the plasma membrane level, followed by caveolin1 localization to EEA1- and Rab5-positive endosomes within 1 hour of oleic acid treatment, and subsequently lipid droplet accumulation. Caveolin1-lipid droplet trafficking does not depend on lysosomal activity. Surprisingly, the deletion of the central caveolin1 β-barrel (Cav1-F160X), a mutation found in lipodystrophy patients, abolished caveolin1-lipid droplet trafficking and lipid droplet growth. Taken together, our results show that dietary lipids induce caveolae uptake, caveolin1 accumulation to lipid droplets and a caveolin1-dependent lipid droplet growth.

## INTRODUCTION

The uptake of nutrients into eukaryotic cells is a highly regulated process that uses multiple entry routes. Electron microscopy (EM) reveals a strikingly large number of 50-100 nm sized invaginations that can populate the entire cytosolic leaflet of the plasma membrane in fat, muscle, or endothelial cells^1,2^. These membrane invaginations are called caveolae and while these structures are known to be involved in endocytosis^2–4^, previous findings also clearly demonstrated that caveolae are essential for cellular lipid uptake^5–7^. Structurally, caveolae are formed by caveolin oligomers (caveolin1-3) that insert into the cytoplasmic side of the phospholipid bilayer and cavin proteins (cavin1-4) that form the striped coat around the invagination typically seen in EM images^8–10^. Previous research showed that caveolin1 and cavin1 are essential for correct caveolae formation^3^. Additionally, the membrane binding protein EHD2 supports the stabilization of the caveolae invaginations at the plasma membrane, and pacsin2 (also called syndapin2) forms the caveolar neck^8,9,11^.

Caveolae have a unique lipid sorting function that leads to the accumulation of specific lipid species and membrane curvatures^12–18^. Therefore, caveolae form a specialized plasma membrane environment that is essential for localization of various plasma membrane channels or receptors such as the insulin receptor^19–22^. Additionally, caveolae are involved in physiological processes such as regulation of membrane tension, endothelial signaling and blood vessel relaxation (via eNOS^23,24^) and to a smaller extend also endocytosis of receptors, plasma membrane proteins or viruses^2,4,25,26^. However, one major function of caveolae includes the lipid uptake in various cell and tissue types^3,27–29^. Caveolae dependent cellular lipid uptake was previously shown in adipocytes, muscle, endothelial cells and fibroblasts^5–7,30–33^. Importantly, loss of caveolae at the plasma membrane reduces lipid uptake and decreases the intracellular accumulation of lipids in various cell and tissue types^5,6,31,34–36^. Mice lacking caveolae are resistant to diet induced obesity, depict a lipodystrophy phenotype and show insulin resistance when placed on a high fat diet^5–7^. Conversely, increased numbers of caveolae result in enhanced lipid uptake and enlarged adipose tissue depots *in vivo*^29,37^. In our previous work, we observed that an increased caveolae uptake rate results in increased cellular lipid uptake *in vivo*, which led to characteristic lipid accumulation in caveolae-containing tissues^30^. Taken together, these data illustrate that caveolae are important regulators of cellular lipid metabolism. In line with the latter, several caveolin1 and cavin1 mutations were found in patients suffering from lipodystrophy^38–41^.

Currently it is not known how caveolae facilitate the lipid uptake at the plasma membrane, how they are involved in intracellular lipid trafficking and metabolism, and what the underlying timescale of these processes may be. Proteomic analysis of the lipid droplet coat revealed the localization of caveolae proteins to lipid droplets^42,43^. Furthermore, a recent publication showed caveolin1 localization to lipid droplets in seipin knockout cells due to ceramide accumulation and associated impairment of the trans-Golgi network^44^. Additionally, it was reported previously that a dominant negative caveolin3 mutant associates to lipid droplets resulting in intracellular cholesterol imbalance^45^. Notably, after stimulation of fibroblasts or adipocytes with extracellular lipids, caveolae associated proteins such as caveolin1 and cavin1 were also detected at lipid droplets^46–52^. However, currently, it is not understood how extracellular lipids trigger caveolae endocytosis, how caveolar trafficking is facilitated and regulated on a molecular level, and why caveolae proteins may be needed at lipid droplets.

Here, we show that extracellular dietary lipids such as oleic acid and cholesterol induce a caveolae curvature shift at the plasma membrane to highly invaginated caveolae. Within 3 hours caveolin1 accumulates at lipid droplets, thereby forming a coat surrounding the lipid droplets. The caveolin1-lipid droplet trafficking depends on caveolae endocytosis, is regulated by EHD2 and early endosomes but not depended on lysosomal activity. Surprisingly, deletion of the central caveolin1 β-barrel or the specific mutation caveolin1-R171P impaired caveolin1-lipid droplet accumulation and lipid droplet growth. Our data suggest that caveolin1 lipid droplet localization is needed for correct lipid droplet growth during cellular lipid storage.

## RESULTS

### Extracellular dietary lipids shift caveolae to spherical membrane curvature

To analyze caveolae mediated lipid trafficking, we distinguished several cellular environments: plasma membrane, cytosol, and lipid droplets. At first, we asked how extracellular lipids modulate caveolae curvature at the plasma membrane. There, caveolae are largely immobile invaginations that can undergo a shift towards more mobile caveolae, which ultimately undergo endocytosis as small vesicles^53–58^. To facilitate caveolae endocytosis, caveolae form most likely an invaginated vesicle-like curvature with a small neck size. Analysis of caveolae curvature in platinum replica EM images is a precise method to investigate caveolae curvature in high resolution and in large quantities at the plasma membrane^1^. To test how extracellular lipids modulate caveolae at the plasma membrane, mouse embryonic fibroblasts (MEFs) were treated with oleic acid for 30min, 1, 3 or 6h, followed by removal of the cell body (unroofing) and platinum replica preparation (PREM^59^). In TEM images, caveolae were identified by their characteristic elongated striped coat and approximal size of 60-80 nm (Fig. 1A). Oleic acid treatment induced only a minor increase in the total caveolae number found at the plasma membrane after 1 and 6h (Fig. 1B). Segmentation of caveolae in PREM images in low, medium or highly curved caveolae (accordingly to previous analysis^1^) revealed a significant decrease of medium curved caveolae to 20% after 1h oleic acid treatment vs. 60% in control cells (Fig. 1C-E). The percentage of low curved caveolae did not change upon oleic acid treatment (Fig. 1C). Surprisingly, within 1h oleic acid treatment the amount of highly curved caveolae increased from 20% in control cells to 65% (Fig. 1E, deeply invaginated, spherical caveolae). Longer treatment of oleic acid resulted in a slight reduction of spherical caveolae, however still significantly more caveolae were highly curved in the presence of oleic acid (Fig. 1E). Next, we treated the cells with cholesterol to identify if other dietary lipids induce a similar caveolae curvature shift. Indeed, MEFs treated with cholesterol for 1h revealed a similar decrease of medium curved caveolae and a significant increase in highly curved, spherical caveolae (Fig. S1A-C). In summary, extracellular dietary lipid treatment resulted in a shift to highly invaginated, spherical caveolae at the plasma membrane.

**Figure 1:**
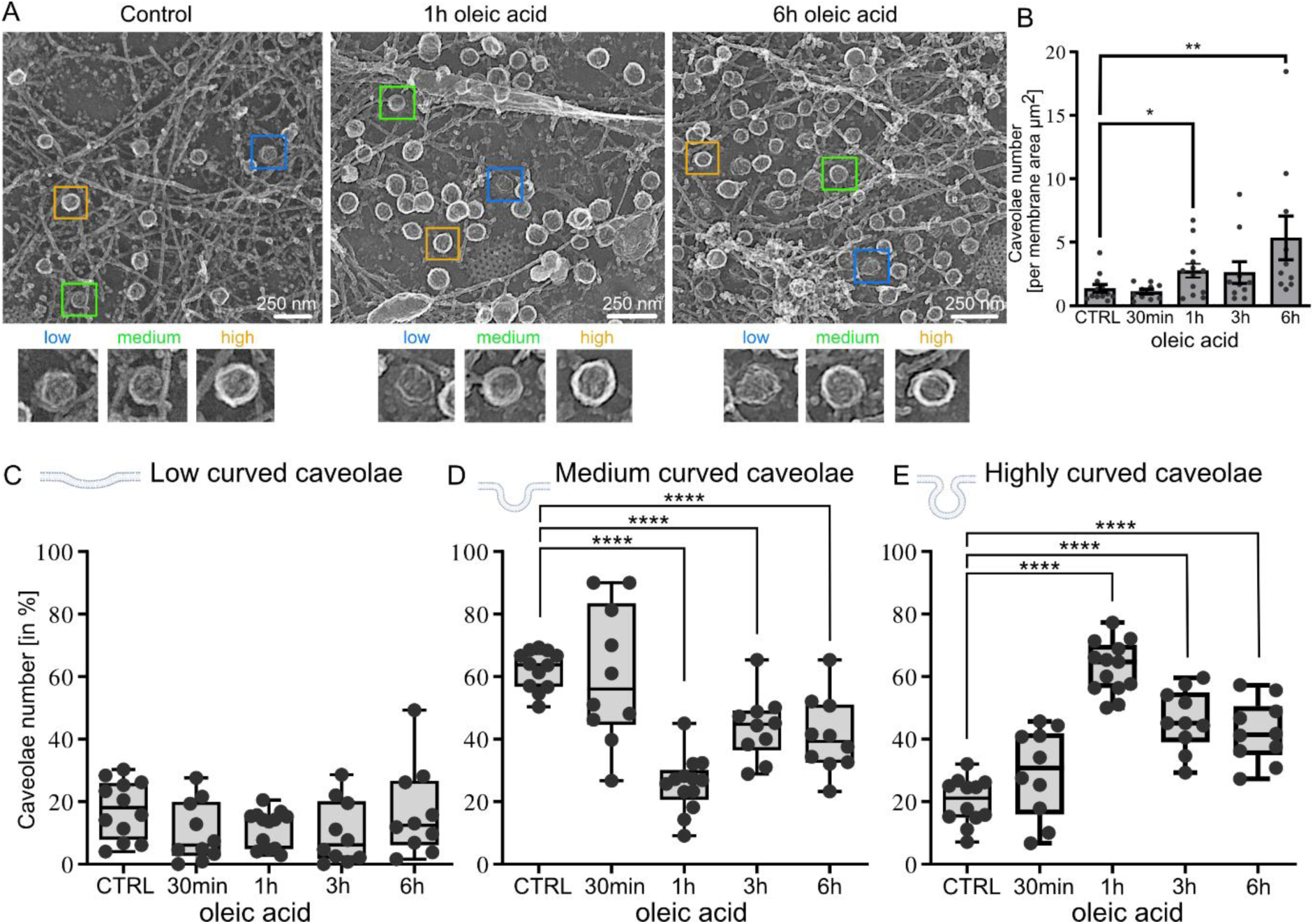
Oleic acid treatment shifts caveolae to spherical curvature at the plasma membrane. (A) MEF were treated with oleic acid for 30 min, 1, 3, or 6 h and were subsequently unroofed and a platinum replica was prepared. Caveolae in platinum replica EM images were segmented in low, medium, highly curved (spherical) groups (scale bar is 250 nm). (B) Caveolae number per plasma membrane area is depicted (bar plot illustrates mean ± SEM, each membrane sheet is depicted). (C-E) Box plots depict percentage of low curved (C), medium curved (D) or highly curved caveolae per cell (box shows min to max, line represents median, each membrane sheet is depicted). For all graphs: n(CTRL) = 12 membrane sheets/7 cells, n(30min) = 10 membrane sheets/5 cells, n(1h) = 13 membrane sheets/7 cells, n(3h) = 10 membrane sheets/6 cells, n(6h) = 10 membrane sheets/5 cells, 2 independent experiments, tested for significant differences with Man-Whitney test, *p≤0.05; **p≤0.01; ***p≤0.001 and ****p≤0.0001.

We next asked if a removal of plasma membrane lipid content shifts caveolae towards a less curved state. To test this, we treated MEFs with methyl-beta-cyclodextrin to remove cholesterol from the plasma membrane^60^. MEFs treated with methyl-beta-cyclodextrin showed a significant increase in low curved caveolae whereas medium curved caveolae were dramatically reduced (Fig. S1D-F). Notably, there was also an increase of highly curved caveolae after methyl-beta-cyclodextrin treatment to 35% vs. 20% in control cells (Fig. S1F), but not as dramatic as observed when MEFs were treated with oleic acid or cholesterol for 1-6h (up to 70%, Fig. 1E, Fig. S1C). Caveolae radius measurements revealed that either oleic acid, cholesterol or methyl-beta-cyclodextrin treatment resulted in significant increased sizes of low curved caveolae (up to 1h treatment, Fig. S2), whereas highly curved caveolae showed a significant smaller radius in oleic acid or cholesterol treated MEFs (Fig. S2). Lastly, we asked if extracellular lipid treatment may influence other endocytic membrane structures in a similar manner. Therefore, we focused on clathrin-mediated endocytosis to evaluate the effect of oleic acid treatment on flat, dome or spherical clathrin structures^61^. Surprisingly, 1h and 6h oleic acid treatment significantly increased the percentage of flat clathrin structures up to 78% (vs. 38% in untreated MEFs, Fig. S3) and significantly reduced dome-like clathrin coated invaginations (Fig. S3). In contrast to caveolar invaginations, the amount of spherical clathrin structures (pits) was not elevated after oleic acid treatment (Fig. S3D). In summary, the treatment of extracellular dietary lipids oleic acid or cholesterol results in a shift of caveolar curvature from medium curved (bulb-shaped) caveolae to highly curved (spherical) caveolae. Assuming that caveolae get endocytosed in this highly curved state, we next asked where caveolae migrate intracellularly.

### Extracellular lipids trigger caveolin1 trafficking to lipid droplets

To follow caveolae in the cytosol, we overexpressed EGFP-tagged caveolin1 in MEFs followed by oleic acid treatment for 1, 3 and 6h. In line with previous research^33,46^, we observed an accumulation of caveolin1 at the lipid droplets starting after 1h oleic acid treatment but becoming more prominent after 3 and 6h (Fig. 2A, B). As expected, lipid droplet sizes increased significantly after 3 and 6h of oleic acid treatment (Fig. 2C). Fig. 2A illustrates the caveolin1-EGFP accumulation around the lipid droplets (stained by Nile Red, in magenta) whereby a distinct coat is formed after 3 and 6h of oleic acid treatment. Line scan analysis of caveolin1 positive lipid droplets further indicate the coat formation of caveolin1 surrounding lipid droplets (Fig. 2A, inset a). Notably, only 55% of all cytosolic lipid droplets were coated with caveolin1 (Fig. 2D).

**Figure 2:**
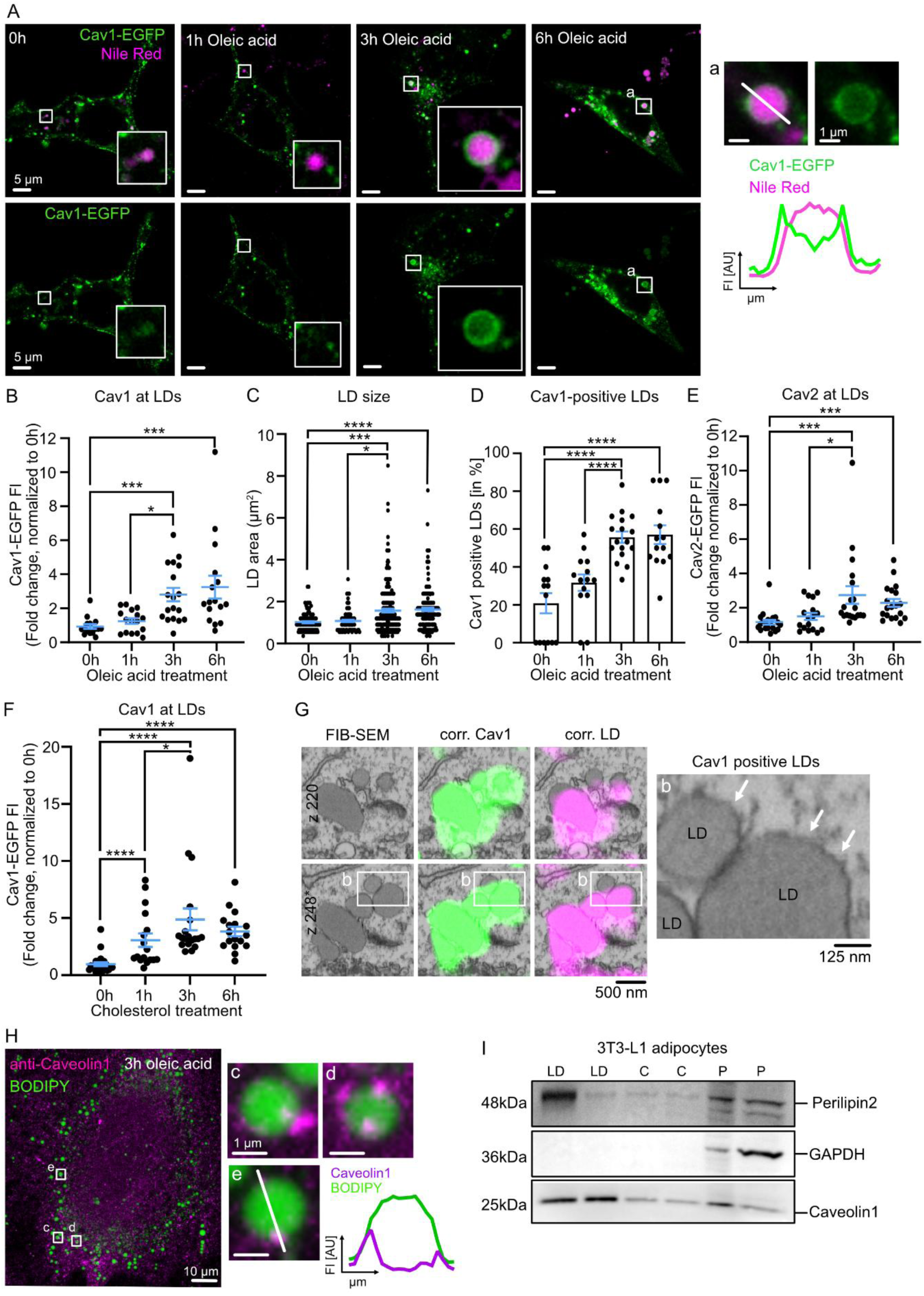
Oleic acid treatment triggers caveolin1 accumulation at lipid droplets. (A) Representative confocal microscopy images of MEFs transfected with caveolin1-EGFP (green), treated with oleic acid for 1, 3, or 6h. Lipid droplets were stained with Nile Red (magenta). Scale bar is 5 µm. Inset (a) after 6h oleic acid treatment, intensity plot profile of caveolin1-EGFP (green) and Nile Red (magenta) across the diameter of a lipid droplet (white line in inset a), scale bar 1 µm. (B) Normalized caveolin1-EGFP fluorescence intensity (FI) measured at lipid droplets, each spot represents averaged fluorescence intensities per cell (n(0h) = 15, n(1h) = 16, n(3h) = 18, n(6h) = 16). (C) Lipid droplet (LD) area in µm^2^, each spot represents a single LD (n(0h) = 88, n(1h) = 89, n(3h) = 186, n(6h) = 177). (D) Percentage of LDs positive for caveolin1 relative to the total amount of LDs per cell (n(0h) = 15, n(1h) = 16, n(3h) = 18, n(6h) = 16). (E) Plot depicts normalized caveolin2-EGFP fluorescence intensity (FI) measured at lipid droplets, each spot represents averaged fluorescence intensities per cell (n(0h) = 20, n(1h) = 17, n(3h) = 19, n(6h) = 19). (F) Plot illustrates normalized caveolin1-EGFP fluorescence intensity (FI) measured at lipid droplets after cholesterol treatment, averaged fluorescence intensities per cell (n(0h) = 23, n(1h) = 18, n(3h) = 17, n(6h) = 15). (G) Correlative confocal fluorescence microscopy and FIB-SEM illustrate caveolin1-positive lipid droplets (caveolin1 in green, lipid droplets (LD) in magenta, 2 individual FIB-SEM slices are shown, scale bar is 500 nm). Inset b depict lipid droplets, arrow heads indicate lipid droplet coat (scale bar is 125 nm). (H) Representative confocal microscopy image of 3h oleic acid treated MEF immuno-labelled against caveolin1 (magenta) and lipid droplets stained with BODIPY (green, scale bar is 5 µm), insets c-e illustrate caveolin1 foci at lipid droplets, intensity plot profile of caveolin1 (magenta) and BODIPY (green) across the diameter of a lipid droplet in (e, white line), scale bar is 1 µm. (I) Western Blot depicts isolated lipid droplet (LD), cytosolic (C), and membrane and cell debris pellet (P) fractions of 3T3-L1 adipocytes, immuno-labelled against Perilipin2, GAPDH and caveolin1 (2 independent experiments are shown). All graphs illustrate mean ± SEM, minimum 5 cells/experiment, 3 independent experiments, tested for significant differences with multiple comparison Kruskal-Wallis test, *p≤0.05; **p≤0.01; ***p≤0.001 and ****p≤0.0001.

Next, we tested if caveolin2 can also migrate to lipid droplets after oleic acid treatment. We observed a similar accumulation of caveolin2-EGFP and increase in LD size after 1, 3 and 6h of oleic acid treatment (Fig. 2E, Fig. S4A-C). Similar to our previous results for caveolin1, caveolin2 also targeted only 60% of all lipid droplets (Fig. S4D). Additionally, we tested if other dietary lipids trigger caveolin1 accumulation at the lipid droplets. Indeed, caveolin1-EGFP also accumulated at lipid droplets after 1, 3 and 6h cholesterol treatment whereby the most prominent caveolin1-lipid droplet coat was found after 3h (Fig. 2F, Fig. S5). Next, we applied triolein to MEFs overexpressing caveolin1-EGFP for 3h in different concentrations (diluted in serum-free DMEM) to test if neutral lipids can also induce caveolin1 migration to lipid droplets. Triolein is a triglyceride containing three oleic acid esters bound to glycerol (glyceryl trioleate). However, no caveolin1-EGFP accumulation at the lipid droplets was observed after 3h in presence of triolein (Fig. S6A, B). Furthermore, lipid droplet sizes did not increase with triolein treatment, but rather decreased compared to untreated cells (Fig. S6C). This indicates that neutral lipids cannot trigger caveolin1 lipid droplet accumulation, whereby the observed reduction in lipid droplet sizes may be induced due to 3h incubation in serum-free medium.

Next, we wondered if the observed caveolin1 accumulation to lipid droplets results from caveolar vesicles or caveolin1 alone. Therefore, we investigated caveolin1 positive lipid droplets by a correlative fluorescence and FIB-SEM approach (Fig. 2G, Fig. S7). As expected, in untreated MEFs caveolae were found at the plasma membrane showing specifically caveolin1 fluorescence (Fig. S7A-C, Video-S1). Detailed inspection of caveolin1 positive lipid droplets in 3D FIB-SEM volume of MEFs expressing caveolin1-EGFP treated with oleic acid revealed no caveolar vesicles or extensive endoplasmic reticulum (ER) accumulation in close proximity (Fig. 2G, inlet b with arrows, Fig. S7D-G, Video-S2-4). This suggests that caveolin1 alone, and not caveolae vesicles, accumulates at lipid droplets when stimulated with oleic acid.

We next investigated endogenous caveolin1 levels at lipid droplets by immuno-fluorescence microscopy against caveolin1 to verify that caveolin1 also accumulated to lipid droplet when not overexpressed (as also shown previously^33,42,49,62^). Again, after 3h of oleic acid treatment we observed caveolin1 accumulation at the lipid droplets (Fig. 2H, Fig. S8A, B). However, instead of a complete caveolin1 coat we detected several caveolin1 foci at the lipid droplets (Fig. 2H, insets c-e, Fig. S8A). Line scan analysis revealed the precise localization of the caveolin1 foci at the lipid droplet (Fig. 2H, inset e). Additionally, we tested endogenous caveolin1 accumulation in differentiated 3T3-L1 adipocytes containing large lipid droplets (Fig. 2I, Fig. S8C-E). Similar to MEFs treated with oleic acid, caveolin1 foci were detected at the lipid droplets in 3T3-L1 adipocytes (Fig. S8E), suggesting that caveolin1 migrates to lipid droplets. The observed difference in caveolin1 accumulation (foci vs. coat formation) to lipid droplets in endogenous vs. over-expressing caveolin1 cells most-likely results from different caveolin1 levels in these cells. Moreover, the accessibility of caveolin1-antibody to the caveolin1 at the lipid droplet may be reduced due to different steric orientation. In addition, we isolated the lipid droplet fractions of 3T3-L1 adipocytes and confirmed caveolin1 localization by Western Blot (Fig. 2I). In summary, we could show specific caveolin accumulation to lipid droplets within 3h after dietary lipid treatment in endogenous and caveolin1 or 2-overexpressing MEFs and 3T3-L1 adipocytes.

To verify that we detect active caveolin1 migration and localization to lipid droplets only when triggered with oleic acid, we applied a splitAPEX2 system that allows to identify organelle/protein contacts^63,64^. Thereby, caveolin1 is tagged with one APEX2-fragment (EX) and perilipin1, as lipid droplet marker, with the other APEX2 fragment (AP) (Fig. S9A). Biotinylation of surrounding proteins can only be detected when both APEX2 fragments are in close proximity^64^ due to migration of caveolin1 to lipid droplets (Fig. S9B). MEFs co-expressing caveolin1-EX and perilipin1-AP were treated with oleic acid for 3h, after which the biotinylation reaction was started and the cells were fixed. The biotinylation was observed by a streptavidin antibody conjugated to a fluorescent probe^64^. Indeed, lipid droplets (stained with BODIPY, green) were biotinylated specifically only in the presence of oleic acid (Fig. S9C). In untreated MEFs no biotinylation could be detected (Fig. S9D). Proteomic analysis of biotinylated proteins isolated from 3h oleic acid treated MEFs co-expressing caveolin1-EX and perilipin1-AP revealed characteristic lipid droplet proteins such as perilipin2 and -3, ACSL4 and -6, as well as fatty acid binding proteins Fabp4 and 5 (Fig. S10). In line with our previous FIB-SEM analysis showing no caveolar vesicles in close proximity to caveolin1 positive lipid droplets, we also did not detect an increased accumulation of caveolae proteins such as caveolin2, cavin1 or cavin2. Taken together, these experiments showed that oleic acid specifically induced caveolin1 accumulation to lipid droplets within 3h.

Previous studies showed that insulin induces phosphorylation of caveolin1 tyrosine 14, which is needed for caveolae endocytosis^20,65–67^. Therefore, we tested if caveolin1-lipid droplet trafficking depends on insulin. However, we did not detect any difference in caveolin1-EGFP accumulation to lipid droplets after oleic acid treatment in the absence or presence of insulin (Fig. S11). A recent publication showed that caveolin1 is mis-localized to lipid droplets when the intracellular ceramide balance is impaired due to long-term oleic or palmitic acid treatment^44^. Treatment with the serine palmitoyltransferase inhibitor (myriocin), which blocks sphingosine biosynthesis, rescued the caveolin1 mis-localization to lipid droplets^44^. However, when the similar sphingosine inhibitor (myriocin) was applied to MEFs expressing caveolin1-EGFP together with 3h oleic acid treatment we observed no impaired caveolin1 accumulation to lipid droplets (Fig. S12). This suggest that a short-time inhibition of *de novo* sphingolipid synthesis by myriocin does not influence oleic acid induced caveolin1 trafficking. The previously observed altered caveolin1 lipid droplet accumulation^44^ may also be caused by different lipid treatments (e.g. lipid species and duration). Taken together, we showed that the dietary lipids oleic acid and cholesterol trigger caveolin1 trafficking to lipid droplets within 3h independently of insulin treatment.

### Increase in membrane fluidity is not sufficient to induce caveolin1- lipid droplet trafficking

As the incorporation of oleic acid molecules into the lipid bilayer of the plasma membrane may increase membrane fluidity, we measured lipid diffusion of DiI-C18 before and after oleic acid treatment as a marker for membrane fluidity (Fig. 3A). As expected, lipid diffusion increased with duration of oleic acid treatment which indicates increasing membrane fluidity (Fig. 3B, C). To determine if a raised membrane fluidity alone triggers caveolae uptake and caveolin1 accumulation at lipid droplets, we artificially increased membrane fluidity by alcohol treatments (hepta-, octa-, nonanol)^68,69^. However, no caveolin1 accumulation at the lipid droplets was observed after 3h (Fig. 3D-E). As expected also the lipid droplet sizes did not increase after alcohol treatments (Fig. 3F). This indicates that an increase in membrane fluidity itself is not sufficient to induce caveolin1 trafficking to lipid droplets, but that the uptake and/or incorporation of lipids to caveolae is a necessary step for caveolae mediated lipid uptake. To further confirm this observation, we also treated caveolin1-EGFP expressing MEFs with oleic acid complexed to albumin (BSA). Oleic acid bound to BSA represents the physiological relevant fatty acid transport mechanism, allowing a better solubility of oleic acid in cell culture medium. Notably, for the oleic acid uptake into cells the fatty acids need to dissociate from albumin^70^. Again, we could detect caveolin1 accumulation at lipid droplets for various physiological relevant oleic-acid-BSA concentrations after 3h (Fig. 3G-H, Fig. S13). When comparing oleic acid and oleic acid-BSA treated MEFs both treatments showed similar caveolin1-EGFP accumulation to lipid droplets (Fig. S13E). MEFs treated with 100 µM oleic acid-BSA revealed a significantly higher caveolin1-EGFP fluorescence intensity at lipid droplets compared to oleic acid treated cells (Fig. S13E, 100 µM). In summary, these results show that an increase in membrane fluidity alone is not able to trigger caveolin1 trafficking, but rather plasma membrane-integrating lipids are needed. Next, we investigated how caveolin1 trafficking to lipid droplets is regulated intracellularly and which organelles may play a role in this trafficking route.

**Figure 3:**
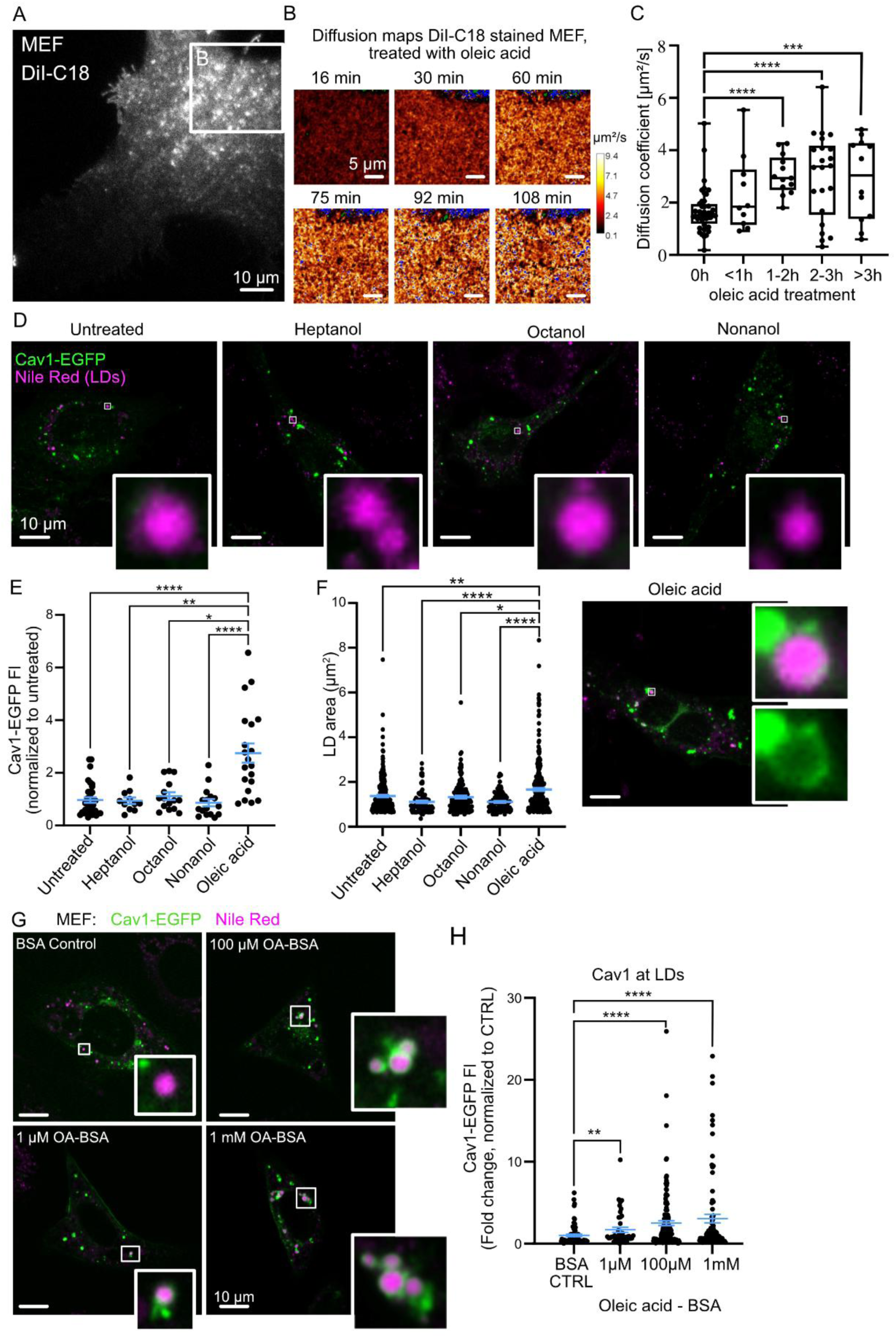
Increase in membrane fluidity is not inducing caveolin1 trafficking and lipid droplet accumulation. (A) Representative TIRF image of the plasma membrane of MEF cells treated with DiI-C18 (scale bar 10 µm). (B) Representative FCS DiI-C18 diffusion maps acquired from MEF plasma membrane area depicted in (A), after oleic acid treatment for 16 – 108 min (scale bar 5 µm, color legend illustrates diffusion coefficients in µm^2^/s). (C) Box plot shows averaged diffusion coefficient per cell (mean ± max/min, n(0h) = 44, n(<1h) = 10, n(1-2h) = 13, n(2-3h) = 22, n(>3h) = 12; 3 independent experiments). (D) Representative confocal images of MEFs transfected with Cav1-EGFP (green), treated with alcohols of different chain lengths (2.5 mM heptanol, 0.5 mM octanol or 0.5 mM nonanol) or oleic acid for 3h, lipid droplets depicted in magenta. Scale bars are 10 µm. (E) Plot depicts normalized caveolin1-EGFP fluorescence intensity (FI) measured at lipid droplets, each spot represents averaged fluorescence intensities per cell (n(untreated) = 35, n(heptanol) = 11, n(octanol) = 15, n(nonanol) = 16, n(oleic acid) = 20). (F) Lipid droplet (LD) area in µm^2^, each spot represents a single LD (n(untreated) = 305, n(heptanol) = 88, n(octanol) = 146, n(nonanol) = 108, n(oleic acid) = 323). (G-H) Representative confocal images of MEFs transfected with Caveolin1-EGFP (Cav1-EGFP, green), treated with BSA or oleic acid-BSA complexes (OA-BSA) for 3h. Lipid droplets are stained with Nile Red (magenta). Scale bars are 10 µm (G). Plot in (H) depicts normalized caveolin1-EGFP fluorescence intensity (FI) measured at lipid droplets, each spot represents a lipid droplet (n(BSA CTRL) = 72, n(1 µM OA-BSA) = 43, n(100 µM OA-BSA) = 133, n(1 mM OA-BSA) = 87). For all graphs E, F and H: mean ± SEM, minimum 5 cells/experiment, 2-3 independent experiments, tested for significant differences with multiple comparison Kruskal-Wallis test, *p≤0.05; **p≤0.01; ***p≤0.001 and ****p≤0.0001).

### Caveolin1 accumulation at lipid droplets is regulated by EHD2

Previous publications showed that caveolae endocytosis is regulated by EHD2^30,53,54,71^. Therefore, we investigated caveolin1 accumulation to lipid droplets in EHD2 knockout^30^ (KO) and EHD2 overexpressing MEFs (Fig. 4A-C). As shown previously, EHD2 stabilizes caveolae at the plasma membrane and loss of EHD2 results in faster caveolae uptake^30,53,71^. Therefore, one would expect an accelerated caveolin1 accumulation at lipid droplets in absence of EHD2. Indeed, when wildtype and EHD2 KO MEFs were treated with oleic acid, an increased caveolin1-EGFP accumulation was already observed after 1h in EHD2 lacking cells that did not increase further after 3h (Fig 4D, E). Contrary, in MEFs overexpressing EHD2-EGFP, caveolin1 accumulation at lipid droplets was completely abolished (Fig. 4F, caveolin1-mApple). In summary, these results illustrate that for caveolin1 accumulation at the lipid droplets, caveolae endocytosis is an essential step and that the migration of caveolin1 from the plasma membrane to lipid droplets is regulated by EHD2.

**Figure 4:**
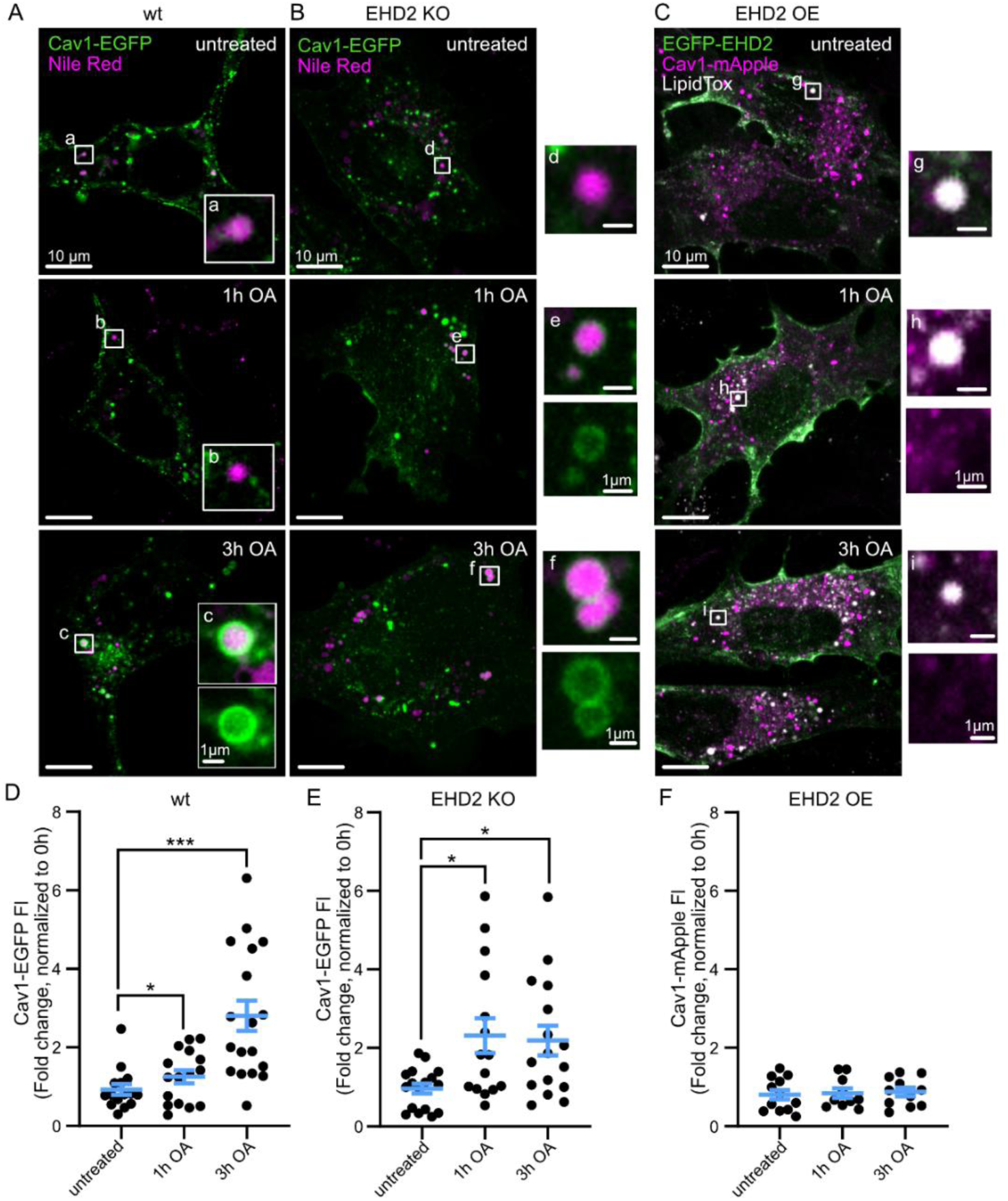
Caveolin1 lipid droplet accumulation is regulated by EHD2. (A-B) Representative confocal images of wildtype (A, wt) or EHD2 KO MEFs (B) expressing caveolin1-EGFP and treated with oleic acid for 3h. Subpanels a-f indicate caveolin1 accumulation to lipid droplets (Nile Red, magenta). Scale bars are 10 µm, subpanels 1 µm. (C) Representative EHD2 KO MEF overexpressing EGFP-EHD2 (green) and caveolin1-mApple (magenta), treated with oleic acid for 3h, lipid droplets were stained with LipidTox (white). Subpanels g-i illustrate lipid droplets (white) and caveolin1 (magenta). (D) Plot depicts normalized caveolin1-EGFP fluorescence intensity (FI) measured at lipid droplets in wt MEFs, each spot represents averaged fluorescence intensities per cell (n(untreated) = 15, n(1h OA) = 16, n(3h OA) = 18). (E) Plot depicts normalized caveolin1-EGFP fluorescence intensity (FI) measured at lipid droplets in EHD2 KO MEFs, each spot represents averaged fluorescence intensities per cell (n(untreated) = 17, n(1h OA) = 15, n(3h OA) = 16). (F) Plot depicts normalized caveolin1-mApple fluorescence intensity (FI) measured at lipid droplets in EHD2 OE MEFs, each spot represents averaged fluorescence intensities per cell (n(untreated) = 18, n(1h OA) = 16, n(3h OA) = 17). For all graphs: mean ± SEM, minimum 5 cells/experiment, 3 independent experiments, tested for significant differences with Kruskal-Wallis test (subpanels D and E) or one-way ANOVA (subpanel F), *p≤0.05; ***p≤0.001 and ****p≤0.0001

### Caveolin1 trafficking to lipid droplets depends on endosomal maturation but not lysosome activity

Next, we asked if caveolin1-lipid droplet trafficking depends on the classical endo-lysosomal pathway. To test this, we inhibited lysosomal activity by treating MEFs expressing caveolin1-EGFP with Bafilomycin A1 (V-ATPase inhibitor^72^), thereby preventing the acidification of the lysosome (Fig. 5A). Efficient Bafilomycin A1 treatment was evaluated by Lysotracker staining of live cells (Fig. S14). Caveolae endocytosis was induced by oleic acid treatment and caveolin1-EGFP fluorescence intensity at the lipid droplets was measured (Fig. 5A, B). Surprisingly, lysosome inhibition did not impair caveolin1 accumulation at lipid droplets. Compared to DMSO treated MEFs caveolin1-EGFP fluorescence intensity was slightly decreased, however similar amounts of caveolin1 positive lipid droplets were detected in Bafilomycin A1 compared to DMSO treated MEFs (Fig. 5B). Next, we blocked lipid export from the lysosome by treating MEFs with NPC1 inhibitor U18666a^73,74^. Following 3h oleic acid treatment, caveolin1-EGFP accumulation at the lipid droplets was analyzed (Fig. 5C). Again, we did not detect any differences between U18666a treated MEFs and control cells (Fig. 5D). Next, we investigated the involvement of the endosome in caveolin1-lipid droplet trafficking. Therefore, we inhibited at first early-to-late-endosomal maturation by using vacuolin-1^75,76^ (Fig. 5E). Surprisingly, when analyzing caveolin1-EGFP at the lipid droplets, the vacuolin-1 treated MEFs showed a significant decrease in caveolin1 accumulation compared to DMSO treated cells (Fig. 5F). In line with this, the amount of caveolin1 positive lipid droplets was also strongly reduced in vacuolin-1 treated MEFs (Fig. 5F). To verify this observation, we next inhibited Vps34, a class III phosphoinositide 3-kinase that regulates endosome maturation and sorting^77^. MEFs expressing caveolin1-EGFP were treated with Vps34 inhibitors Vps34-IN1 or SAR405^78^ and oleic acid for 3h followed by lipid droplet staining (Fig. 5G). Similar to the vancuolin-1 experiments, both Vps34 inhibitors impaired caveolin1-EGFP accumulation at lipid droplets. Compared to DMSO treated MEFs inhibition of Vps34 resulted in a significant reduced caveolin1-EGFP fluorescence intensity at lipid droplets (Fig. 5H). In line with the latter, also the number of caveolin1 positive lipid droplets per cell were significantly reduced after Vps34 inhibition (Fig. 5H). Taken together, oleic acid induced caveolin1 accumulation to lipid droplets depends on endosome maturation but not on lysosome activity. This suggests that after lipid induced caveolae endocytosis caveolae may migrate to the endosome, fuse with the endosomal membrane after which caveolin1 can accumulate to lipid droplets. To test this hypothesis, we investigated caveolin1 localization within the endosome prior to its lipid droplet accumulation.

**Figure 5:**
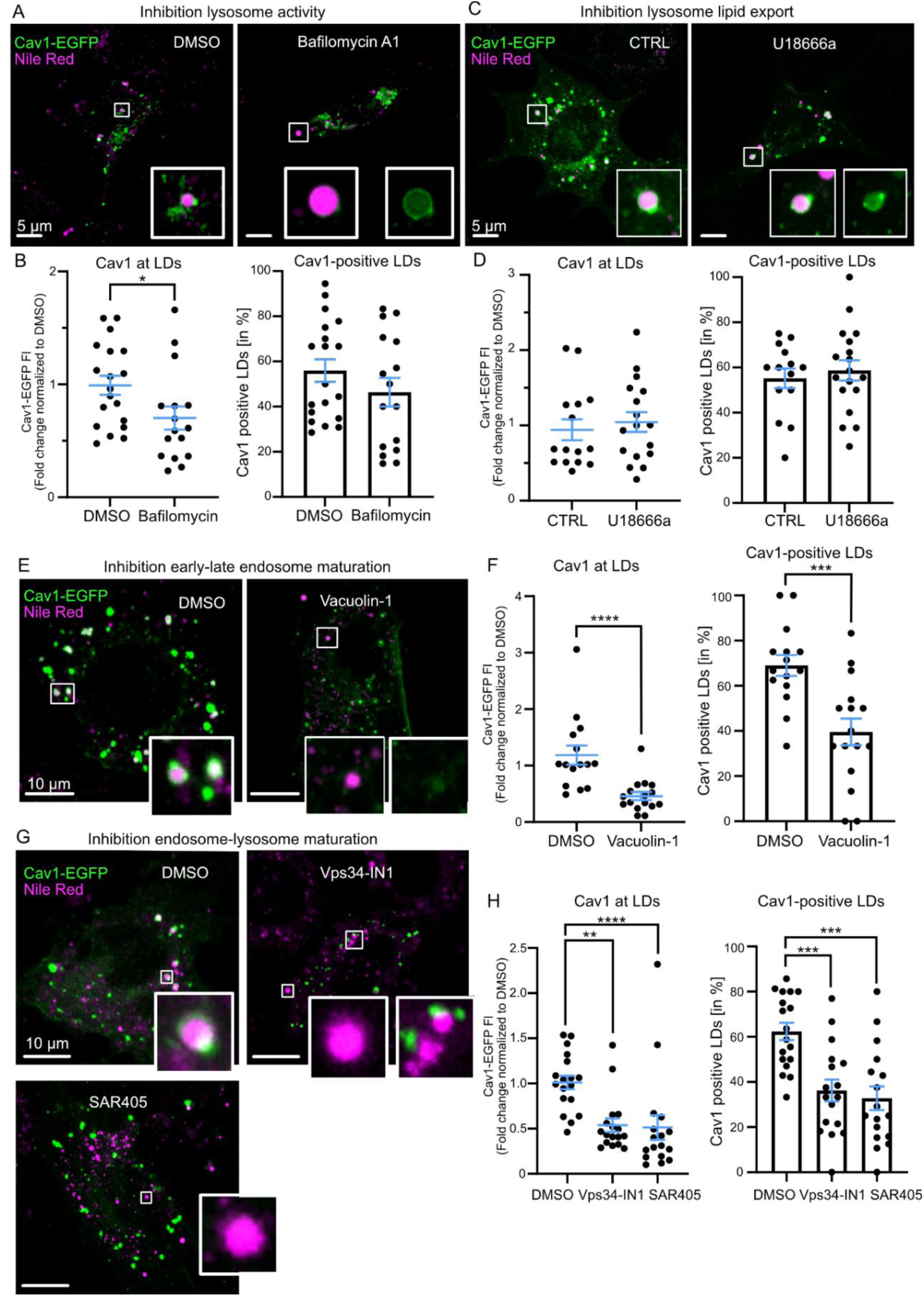
Caveolin1-lipid droplet accumulation depends on endosome maturation but not lysosomal activity. (A) Representative confocal images of MEFs transfected with caveolin1-EGFP (green), treated with 100 nM Bafilomycin A1 or DMSO control (0.1%) and oleic acid for 3h. Cells were stained with Nile Red to visualize lipid droplets (magenta), scale bars are 5 µm. (B) Plot depicts normalized caveolin1-EGFP fluorescence intensity (FI) measured at lipid droplets (LDs, each spot represents averaged fluorescence intensities per cell) or percentage of lipid droplets positive for caveolin1 relative to the total number of lipid droplets/cell (n(DMSO)= 19, n(Bafilomycin) = 16). (C) Representative confocal images of MEFs transfected with caveolin1-EGFP (green) and treated for 3h with oleic acid and 2.5 µM U18666a or H2O control. (D) Similar plots as described in (B), (n(CTRL)= 15, n(U18666a) = 17-18). (E) Representative confocal images of MEF cells transfected with caveolin1-EGFP (green), treated for 3h with OA and 1 µM vacuolin-1 or DMSO (0.05%). Scale bars are 10 µm. (F) Similar plots as described in (B), (n(DMSO)= 15, n(Vacuolin-1) = 16). (G) Representative confocal images of MEFs transfected with caveolin1-EGFP (green) and treated for 3h with oleic acid and 1 µM Vps34-IN1, 1 µM SAR405 or DMSO (0.05%). (H) Similar plots as described in (B), (n(DMSO)= 18, n(Vps34-IN1) =17, n(SAR405) =17).For all graphs: mean ± SEM, minimum 5 cells/experiment, 3 independent experiments, tested for significant differences with Mann-Whitney test (panel D (cav1 at the LDs) and panel F (cav1 at the LDs)) or Welch’s t test (panel B (both graphs)), panel D (cav1 positive LDs) and panel F (cav1 positive LDs)), *p≤0.05; ***p≤0.001 and ****p≤0.0001.

Endosomal compartments can be identified by specific marker proteins such as EEA1 and Rab5 for early endosomes or Rab7 for late endosomes^79^. To investigate if oleic acid treatment triggers caveolin1 migration to endosomes, MEFs expressing caveolin1-EGFP were treated for 30min, 1h and 3h with oleic acid and EEA1, Rab5 or Rab7 positive endosomes were identified by immuno-fluorescence staining with specific antibodies against these marker proteins (Fig. 6A-C). Indeed, within 30min of oleic acid treatment EEA1- and Rab5-positive endosomes showed caveolin1 accumulation (Fig. 6A, subpanels a and c). Rab7 endosomal compartments as well as lipid droplets did not show prominent caveolin1 co-localization after 30 min oleic acid treatment (Fig. 6A, subpanels b, d-f). Prolonged oleic acid treatment resulted in further caveolin1 localization to EEA1 and Rab5 positive endosomes (Fig. 6B, subpanels a-d), whereby after 3h oleic acid treatment caveolin1-EGFP accumulated most prominently to lipid droplets (Fig. 6C, subpanels a-d). When we blocked early-to-late endosome maturation by vacuolin-1 (Fig. S15), we observed caveolin1 accumulation to large EEA1 and Rab5 positive compartments (Fig. S15, subpanels d-e) but not to lipid droplets (Fig. S15, subpanels a, c, d) further indicating that oleic acid treatment triggers caveolin1 trafficking to endosomes prior to lipid droplet accumulation. Additionally, correlative caveolin1-EGFP and FIB-SEM of MEF treated with oleic acid revealed close contact between caveolin1-positive lipid droplets and endosomes (Fig. 6D). In summary, our data showed that oleic acid triggers caveolin1 localization to endosomes within 30min and 1h prior to caveolin1 lipid droplet accumulation after 3h.

**Figure 6:**
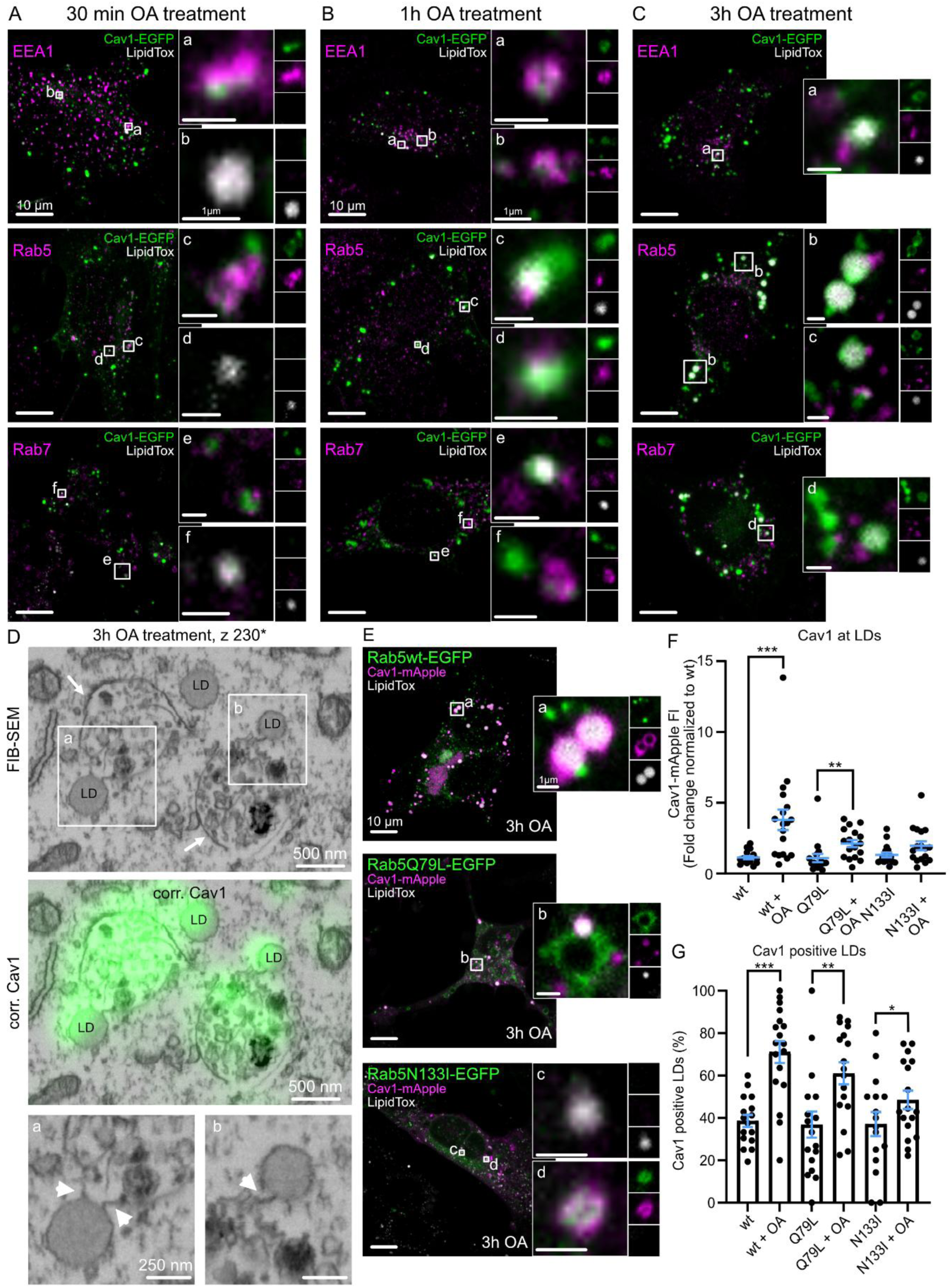
Oleic acid triggers caveolin1 migration to endosomes prior to its lipid droplet localization. (A-C) Representative confocal images of MEFs transfected with caveolin1-EGFP (green), treated with oleic acid for up to 3h, followed by immuno-fluorescence staining against EEA1, Rab5 or Rab 7 (magenta) and LipidTox staining (white, lipid droplets, scale bars are 10 µm). Insets represent single confocal planes of acquired z-stack (scale bars are 1 µm). (D) Correlative confocal fluorescence microscopy and FIB-SEM illustrate caveolin1-positive lipid droplets (LD) and endosomal compartments (indicated by arrows), caveolin1 in green, 1 FIB-SEM slice is shown, scale bar is 500 nm. Insets depict lipid droplet-endosome contacts (arrow heads, scale bar is 250 nm). (E) Representative confocal images of MEFs expressing caveolin1-mApple (magenta) and Rab5-EGFP, Rab5-Q79L-EGFP, or Rab5-N133I-EGFP (green). Cells were treated for 3h with oleic acid (OA) and lipid droplets were stained with LipidTox (white). Scale bars are 10 µm, insets scale bars are 1 µm. (F) Plot depicts normalized caveolin1-mApple fluorescence intensity (FI) measured at lipid droplets, each spot represents averaged fluorescence intensities per cell (n(wt) = 16, n(wt+OA) = 18, n(Q79L) = 17, n(Q79L+OA) = 17, n(N133I) = 16, n(N133I+OA) = 17). (G) Number of caveolin1-positive lipid droplets per cell (in %, n(wt) = 16, n(wt+OA) = 18, n(Q79L) = 17, n(Q79L+OA) = 17, n(N133I) = 16, n(N133I+OA) = 17). For all graphs: mean ± SEM, minimum 5 cells/experiment, 3 independent experiments, tested for significant differences with Kruskal-Wallis test (panel E) or one-way ANOVA (panel F), **p≤0.01; ***p≤0.001 and ****p≤0.0001.

To quantitatively validate if oleic acid induced caveolin1-endsome trafficking is needed for correct caveolin1 localization to lipid droplets, we expressed in MEFs constitutively active Rab5 (Rab5-Q79L-EGFP^80^) or dominant-negative Rab5 mutants (Rab5-N133I-EGFP^81^) together with caveolin1-mApple (Fig. 6E). This allowed us to modulate early-to-late endosome maturation mechanistically on the Rab5 level and independent of inhibitor treatments. As expected, overexpression of Rab-Q79L-EGFP resulted in enlarged Rab5 endosomes (Fig. 6E, inset b), however caveolin1 accumulation to lipid droplets was not impaired (Fig. 6F, G) although regularly caveolin1-positive lipid droplets and Rab5-Q79L co-localized. Similar to Rab5-wt-EGFP expressing MEFs, oleic acid treatment in Rab5-Q79L MEFs triggered a significant increased caveolin1 accumulation to lipid droplets (Fig. 6F). Contrary, when MEFs expressed Rab5-N133I-EGFP, oleic acid induced caveolin1 accumulation to lipid droplets was strongly impaired (Fig. 6E inset c, Fig. 6F, G). Strikingly, caveolin1 was stuck in Rab5-N133I positive endosomes (Fig. 6E inset d). In summary, our inhibitor, endosome-caveolin1 colocalization immuno-fluorescence microscopy and Rab5-mutant experiments revealed that oleic acid treatment triggers caveolin1 migration to endosomes within 30min and 1h, followed by caveolin1 accumulation to lipid droplets after 3h independent of lysosome involvement.

### Caveolin1 β-barrel deletion abolishes caveolin1 trafficking and lipid droplet growth

To further understand why caveolin1 is needed at lipid droplets we generated caveolin1 mutations and investigated their trafficking efficiency to lipid droplets. Specifically, we thereby focused on the central β-barrel of the caveolin1 8S complex^82^ which resembles a lipophilic tunnel that connects the inner plasma membrane phospholipid bilayer area with the cytosol^8,82^. It was suggested previously that fatty acids (and other lipid molecules) may bind within the β-barrel^8,16,82^. At first, we mutated the asparagine residue at the entry of the β-barrel to a proline residue (Cav1-R171P, Fig. 7A), thereby increasing steric hinderance. MEFs expressing Cav1-R171P showed correct plasma membrane localization under basal condition (Fig. S16) but also increased Cav1-R17P-EGFP accumulation to lipid droplets in untreated cells (Fig. 7A, inset b). When oleic acid was added to these cells, a slight increase in caveolin1-lipid droplet accumulation was observed, but that was neither significantly different to untreated MEFs expressing Cav1-R171P-EGFP nor MEFs expressing wt caveolin1 treated with oleic acid (Fig. 7B). Next, we mutated the exit residue of the caveolin1 8S β-barrel from glutamic acid to valine (Cav1-E177V, Fig. 7A). We observed correct plasma membrane localization of Cav1-E177V-EGFP in untreated MEFs (Fig. S16), and oleic acid treatment triggered its accumulation to lipid droplets similar to MEFs expressing caveolin1 wt (Fig. 7C). We therefore conclude that the mutation to valine at the exit side of the caveolin1 8S β-barrel has no effect on caveolin1 trafficking and accumulation to lipid droplets. However, increasing steric hinderance at the entry side by introducing a proline residue resulted in increased lipid droplet accumulation under basal condition.

**Figure 7:**
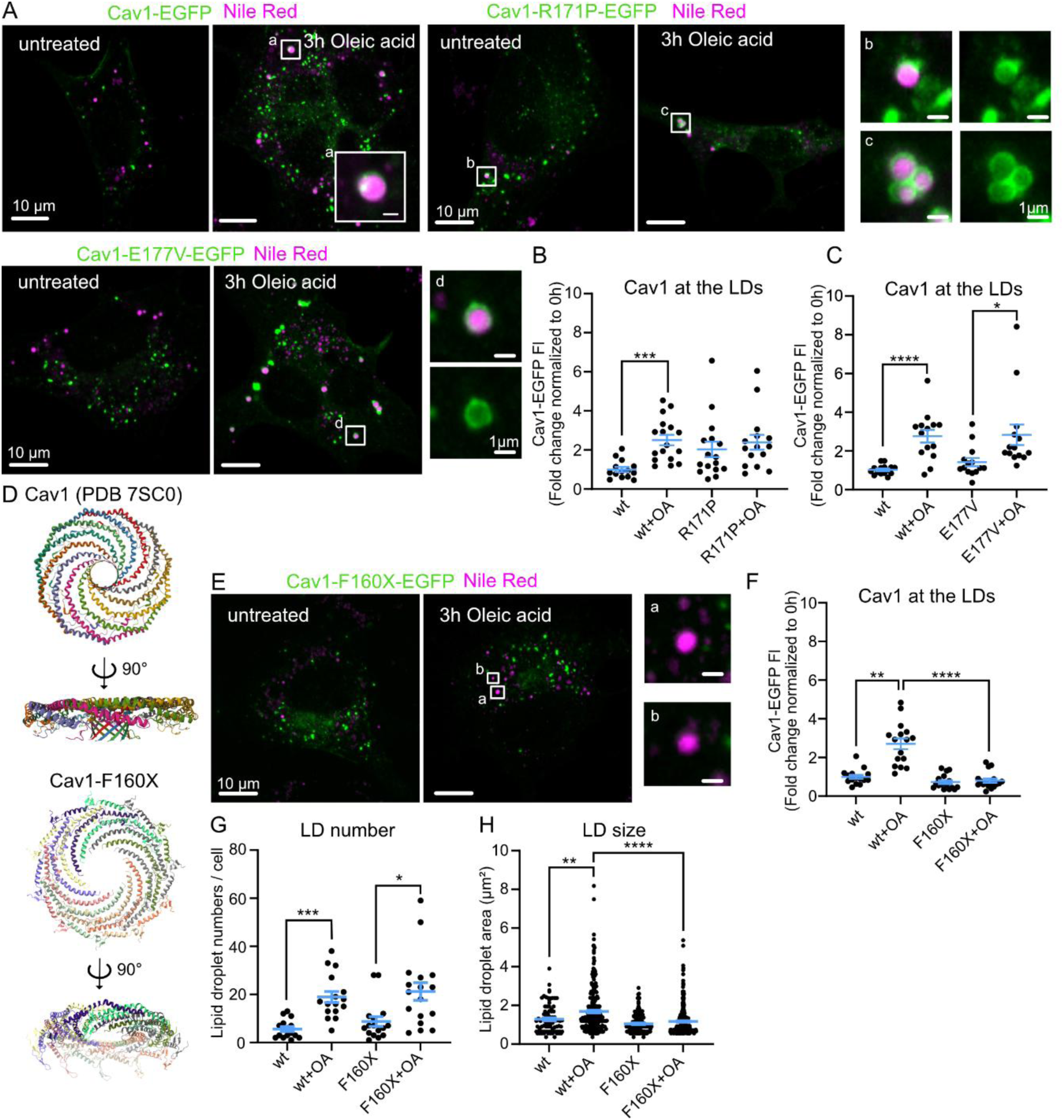
Deletion of caveolin1 β-barrel diminished caveolin1 lipid droplet trafficking and lipid droplet growth. (A) Representative confocal images of MEFs transfected with caveolin1-EGFP, Cav1-R171P-EGFP or Cav1-E177V-EGFP (green) treated with oleic acid for 3h. Cells were stained with Nile Red to visualize lipid droplets (magenta, scale bars are 10 µm), insets a-d show coated lipid droplets (scale bars are 1µm). (B-C) Plot depicts normalized Cav1-R171P-EGFP fluorescence intensity (B) or Cav1-E177V-EGFP fluorescence intensity (C) measured at lipid droplets (LDs, each spot represents averaged fluorescence intensities per cell, for (B): n(wt)= 14, n(wt+OA)= 17, n(R171P)= 16, n(R171P+OA)= 15, for (C): n(wt)= 15, n(wt+OA)= 14, n(E177V)= 14, n(E177V+OA)= 14). (D) Caveolin1 8S protein structure (PDB 7SC0^82^), and the predicted Alphafold (v3^113^) structure of caveolin1-F160X mutant. (E) Representative confocal images of MEFs transfected with Cav1-F160X-EGFP (green) treated with oleic acid for 3h. Cells were stained with Nile Red to visualize lipid droplets (magenta, scale bars are 10 µm), insets (a-b) show enlarged lipid droplets (scale bars are 1 µm). (F) Plot depicts normalized Cav1-F160X-EGFP fluorescence intensity measured at lipid droplets (LDs, each spot represents averaged fluorescence intensities per cell, n(wt) =15; n(wt+OA) = 16, n(F160X) =16, n(F160X+OA) = 17). (G) Plot illustrates total number of lipid droplets per cell (n(wt)= 15, n(wt+OA) = 16, n(F160X) = 16, n(F160X+OA) = 17). (H) Lipid droplet (LD) area in µm^2^, each spot represents a single LD (n(wt) = 84, n(wt+OA) =303, n(F160X) = 140, n(160X+OA) = 361). For all graphs: mean ± SEM, minimum 4 cells/experiment, 3 independent experiments, tested for significant differences with Kruskal-Wallis test, *p≤0.05; ***p≤0.001 and ****p≤0.0001

Several caveolin1 mutations were previously associated to generalized or partial lipodystrophy in human patients^38,83,84^. However, currently it is not understood why these caveolin1 mutations are causing such severe impaired lipid accumulation. The caveolin1 F160X mutation^39–41^ is an interesting candidate as this frameshift mutation causes a complete deletion of the β-barrel within the caveolin1 8S complex (Fig. 7D). To test the effect of this deletion on caveolin1 trafficking and lipid accumulation, we generated a caveolin1-F160X-EGFP construct and expressed it in MEFs (Fig. 7E). When inspecting the plasma membrane localization of this construct we did not observe a different localization compared to caveolin1 wt in untreated MFEs (Fig. S16). Next, we treated MEFs expressing Cav1-F160X-EGFP with oleic acid to trigger caveolin1 trafficking to lipid droplets. However, compared to caveolin1 wt expressing MEFs the measured fluorescence intensity of Cav1-F160X-EGFP at lipid droplets was not increased after 3h of oleic acid treatment (Fig. 7E-F, inset a and b). Lipid droplet numbers per cell were not changed in Cav1-F160X-EGFP expressing MEFs indicating that new lipid droplets were formed after oleic acid treatment (Fig. 7G). Surprisingly, compared to caveolin1 wt MEFs the lipid droplet sizes were significantly reduced in Cav1-F160X-EGFP cells, even showing no detectable size difference between control and oleic acid treatment (Fig. 7H). Based on this observation we next investigated if caveolin1-lipid droplet accumulation may influence lipid droplet growth.

### Caveolin1 accumulation to lipid droplets supports lipid droplet growth

First, we asked if caveolin1 positive lipid droplets are significantly larger in size compared to lipid droplets not targeted by caveolin1. Therefore, we re-inspected our initial experiments in MEFs expressing caveolin1-EGFP (Fig. 2) and analyzed the lipid droplet sizes (Fig. 8A, B). In cells overexpressing caveolin1-EGFP lipid droplets with a detectable caveolin1 signal were larger on average than caveolin1 negative lipid droplets (Fig. 8A), as also observed previously^49,85^. Independently of the duration of oleic acid treatment (0, 1, 3 or 6h) caveolin1 positive lipid droplets were always significantly enlarged compared to caveolin1 negative lipid droplets in the same cell, whereby the averaged lipid droplet sizes increased with duration of the oleic acid treatment (Fig. 8B). Notably, when only comparing caveolin1 negative lipid droplets, the average size did not increase from 0h to 6h treatment (Fig. 8B, no Cav1). This suggest that caveolin1 accumulation is associated with lipid droplet growth. To further test this hypothesis, we investigated lipid droplet sizes in MEFs overexpressing EHD2 resulting in complete lack of caveolin1 accumulation to lipid droplets (Fig. 4F). As illustrated in fig. 8C lipid droplet sizes did not increase during oleic acid treatment in these cells. After 1h oleic acid treatment lipid droplets in MEFs overexpressing EHD2 even showed a slight reduction in their size (Fig. 8C). In line with this, lipid droplet sizes in MEFs overexpressing the dominant-negative Rab5-N133I-EGFP variant, that also impaired caveolin1 lipid droplet accumulation (Fig. 6F), were significantly smaller compared to wt Rab5 (Fig. 8D). Surprisingly, when inspecting caveolin1-R171P we observed many caveolin1 positive LDs (Fig. 7A-C), however the average lipid droplet sizes were not increased after oleic acid treatment although new lipid droplets were formed (Fig. 8E). Furthermore, detailed comparison of caveolin1 positive and negative lipid droplets in MEFs expressing caveolin1-R171P revealed no size increase for caveolin1 positive droplets in contrast to caveolin1 positive lipid droplets in MEFs expressing caveolin1 wt (Fig. 8F). This suggest that the mutated entry side of the caveolin1 β-barrel impairs lipid transfer necessary for lipid droplet growth. In contrast, caveolin1-E177V did not show any difference to caveolin1 wt, whereas the β-barrel deletion in caveolin1-F160X resulted in significantly reduced trafficking and decreased lipid droplets sizes (Fig. 7H). In summary, these results indicate that the lipid induced caveolin1 trafficking to lipid droplets supports lipid accumulation and lipid droplet growth, and that specifically the caveolin1 β-barrel is necessary for caveolin1-lipid droplet localization and lipid droplet growth (Fig. 8G).

**Figure 8:**
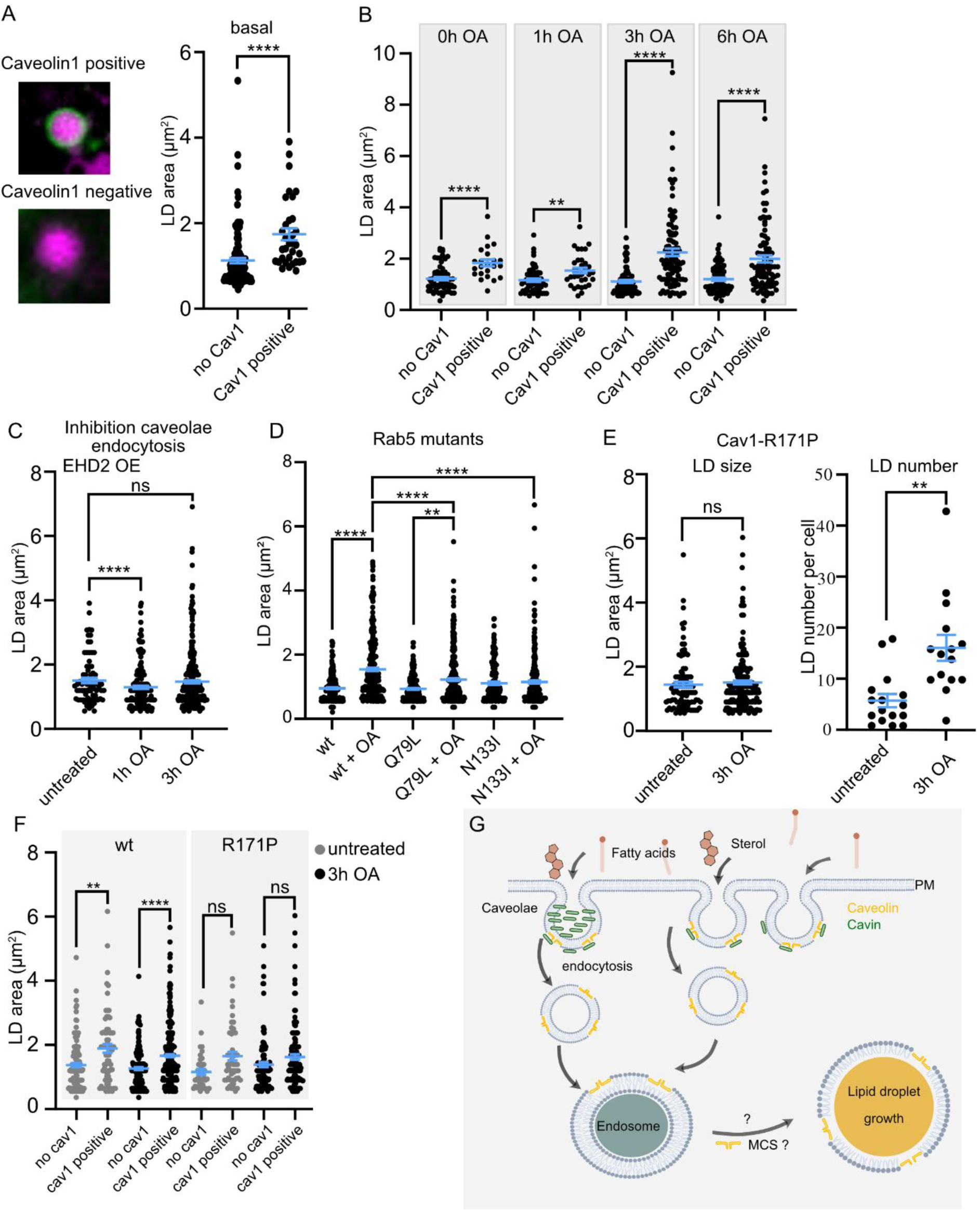
Caveolin1 accumulation at lipid droplets associates with lipid droplet growth. (A-B) Caveolin1 positive and negative lipid droplet sizes were analyzed in MEFs expressing caveolin1-EGFP before (A) and after oleic acid treatment (B). Plots depict lipid droplet (LD) area in µm^2^, each spot represents a single LD (n(0h, no cav1) = 62 / 15 cells, n(0h, cav1 positive)= 23 /15 cells, n(1h, no cav1) = 57/ 16 cells , n(1h, cav1 positive)= 33 /16 cells, n(3h, no cav1)= 80/ 18 cells, n(3h, cav1 positive) = 102/ 18 cells, n(6h, no cav1)= 85/ 16 cells, n(6h, cav1 positive) = 90/ 16 cells). (C) Lipid droplet sizes in MEFs overexpressing EGFP-EHD2 and caveolin1-mApple. Plot depicts lipid droplet (LD) area in µm^2^, each spot represents a single LD (n(untreated) = 76 / 18 cells, n(1h) = 202 / 16 cells, n(3h) = 364 / 17 cells). (D) Plot shows lipid droplets sizes in MEFs co-expressing Rab5 mutants (EGFP-tagged) and caveolin1-mApple; each spots represents a lipid droplet. (n(wt) = 177 / 16 cells, n(wt+OA) = 370 / 18 cells, n(Q79L) = 217 / 17 cells, n(Q79L+OA) = 319 / 17 cells, n(N133I) = 140 / 16 cells, n(N133I+OA) = 246 / 17 cells). (E) Lipid droplet sizes and number of LDs per cell in MEFs overexpressing Cav1-R171P-EGFP in untreated cells vs. cells treated with oleic acid for 3h. Plots depict lipid droplet (LD) area in µm^2^ (left) or number of LDs per cell (right), each spot represents a single LD (left) or a single cell (right) with (n(untreated) = 95 / 16 cells, n(3h OA) = 244 / 15 cells)). (F) LD sizes of caveolin1 positive LDs vs. LDs without caveolin1 in cells expressing wt caveolin1-EGFP or caveolin1-R171P-EGFP in untreated or 3h OA treated conditions. Plot depicts LD sizes in µm^2^, each spot is a single lipid droplet (n(wt, untreated, no cav1)=143 / 28 cells, n(wt, untreated, cav1 positive)=68 / 28 cells, n(wt, 3h OA, no cav1)=219 / 31 cells, n(wt, 3h OA, cav1 positive)=446 / 31 cells, n(R171P, untreated, no cav1)=39 / 16 cells, n(R171P, untreated, cav1 positive)=55 / 16 cells, n(R171P, 3h OA, no cav1)= 109 / 15 cells, n(R171P, 3h OA, cav1 positive)= 134 / 15 cells). (G) Model scheme illustrating caveolin1 – lipid droplet trafficking: Extracellular dietary lipids shift caveolae to highly invaginated curvature, followed by endocytosis and endosome fusion. Afterwards, caveolin1 accumulates at lipid droplets leading to lipid droplet growth. For all graphs: mean ± SEM, minimum 5 cells/experiment, 3 independent experiments, tested for significant differences with Kruskal-Wallis test, **p≤0.01 and ****p≤0.0001

## DISCUSSION

Various studies previously reported the localization of caveolae proteins such as caveolin1 to lipid droplets^42,43,46,48,49,52,62^, even though these proteins are in general found at the plasma membrane where they form caveolae. Currently, the specific cellular function of caveolin1 at lipid droplets is not known, however, caveolin1 knockout mouse models^3,86^ and reported caveolin1 mutations in lipodystrophy patients indicated an important role in cellular lipid accumulation^38^. Here, we now reveal molecular details about caveolin1 trafficking from the plasma membrane to lipid droplets and show that caveolin1-lipid droplet localization is associated with lipid droplet growth. By using high resolution Pt replica EM, we revealed that dietary lipids like oleic acid and cholesterol shift caveolae to spherical curvature at the plasma membrane within 1h of lipid treatment (Fig. 1). When we followed caveolae within the cell, we observed the accumulation of caveolin1 to lipid droplets within 3h whereby no caveolar vesicles and other caveolae related proteins (e.g. cavin1-3) where detected, but rather caveolin1 alone localized to lipid droplets (Fig. 2). Mechanistically, lipid induced caveolin1 trafficking is regulated by EHD2 at the plasma membrane, thereby indicating that caveolae endocytosis might be required for this process (Fig. 4). Lastly, we revealed that caveolin1 accumulation to lipid droplets does not depend on lysosomal activity, but rather on endosome maturation, suggesting a novel trafficking route via endosomes to lipid droplets (Fig. 8G).

Caveolae endocytosis was reported in various cell types^26^ and although these events are rare, and the percentage of plasma membrane proteins internalized by caveolae is around 15%^56^, caveolae endocytosis is essential for the cellular lipid uptake ^25^. To undergo internalization caveolae change their membrane curvature into highly spherical invaginations including a small neck connecting the caveolar bulb/sphere with the plasma membrane^1,15^. Indeed, we observed a shift to highly curved caveolae after oleic acid or cholesterol treatment suggesting that the lipid accumulation primes caveolae for endocytosis. In line with the latter, Hubert et al. previously showed that cholesterol or glycosphingolipids reduce caveolar neck diameter and support caveolae uptake^15^. It can be speculated that the increase in membrane fluidity due to the lipid insertion into the plasma membrane triggers caveolae endocytosis, however, an increase in membrane fluidity alone, is not sufficient to trigger caveolin1-lipid droplet accumulation (Fig. 3). Notably, we did not directly quantify caveolae curvature after alcohol treatment (by PREM), thus conclusions about caveolae curvature under alcohol induced membrane fluidization cannot be made. How caveolae membrane scission is mechanistically facilitated is currently under debate^25,87^, nevertheless, previous studies clearly showed that caveolae can internalize as small vesicles and that their endocytosis is regulated by EHD2^15,30,44,54,71^. In line with the latter, we showed that caveolin1-lipid droplet accumulation is regulated by EHD2 (Fig.4): In the absence of EHD2, caveolae endocytosis is significantly increased^30,53,54^, and this is also reflected in our experiments where an accelerated caveolin1-lipid droplet accumulation was detected. Already after 1h of oleic acid treatment caveolin1 formed a clearly visible coat around the targeted lipid droplets (Fig. 4B) in EHD2 KO MEFs contrary to wt MEFs where the caveolin1 coat is only detectable after 3h. Stabilizing caveolae at the plasma membrane and blocking caveolae endocytosis by overexpression of EHD2^30,71,88^ completely abolished the caveolin1 coat formation at lipid droplets, further indicating that caveolae endocytosis is an essential first step for the lipid induced caveolin1-lipid droplet trafficking. However, we did not directly quantify caveolae endocytosis in live cells within this manuscript, thus alternative uptake routes cannot be fully excluded. Notably, we cannot exclude the possibility that extracellular lipids not only induce caveolae endocytosis but may also trigger caveolae disassembly, a process generally induced by increased membrane tension^4,89,90^. However, our PREM analysis revealed a strong increase in highly curved caveolae indicating that membrane tension is not increased^1,90^ and therefore caveolae disassembly most likely is not triggered. In line, total caveolae numbers at the plasma membrane were also not reduced after lipid treatment (Fig. 1). Recently it was reported that oxidative stress and lipid peroxidation drive caveolae disassembly from the plasma membrane^91,92^. However, increased lipid peroxidation occurs more likely in poly-unsaturated fatty acids (PUFAs) and not mono-unsaturated fatty acids such as oleic acid^93^. Additionally, in all our experiments we observed lipid droplet formation during oleic acid treatment indicating that the internalized oleic acid molecules are converted in neutral triglycerides stored within lipid droplets resistant to lipid peroxidation.

Where caveolae migrate after internalization? We tracked caveolae by using EGFP-tagged caveolin1 as a marker protein for caveolae. As shown previously, caveolin1 localized to lipid droplets whereby we could detect an increase in caveolin1-EGFP at the lipid droplet coat within 1h, which finalizes in a caveolin1-coat surrounding lipid droplets after 3h. Importantly, we also detected endogenous caveolin1 levels at lipid droplets, however there was no caveolin1 coat surrounding lipid droplets but rather several caveolin1 foci consistent with an expression-dependent difference in apparent coat formation. Presumably, depending on the cell type and the endogenous caveolin1 expression levels specific cell types may show more or less caveolin1 foci. In mature white adipocytes containing large lipid droplets and a very reduced cytosolic area, caveolin1-lipid droplet accumulation may occur near the plasma membrane^94^ making it challenging to discriminate exact localization of caveolin1. Future studies in primary adipocytes are needed to elucidate this possibility.

We speculated that we would be able to detect caveolae vesicles at caveolin1 positive lipid droplets, however, correlative FIB-SEM imaging revealed no vesicles in close proximity. In line with the latter, we also did not detect any increased accumulation of other caveolae coat proteins such as cavin1 in proteomic analysis of caveolin1 positive lipid droplets (split-APEX). Therefore, we assume the caveolin1-lipid droplet accumulation is independent of caveolar vesicles at the lipid droplets. So, how can caveolin1 reach lipid droplets? It was shown previously that caveolae endocytosis results in caveolin1 localization to the endosome^58,95–98^. Indeed, we detected caveolin1 localization in endosomal compartments after 30 min – 1h of oleic acid treatment, whereby mainly early endosomal marker EEA1 and Rab5 co-localized with caveolin1. Strikingly, inhibition of early-to-late endosome maturation by vacuolin-1, Vps34 inhibition or over-expression of dominant negative Rab5 mutant (Rab5-N133I) impaired the caveolin1-lipid droplet accumulation tremendously (Fig. 5 and 6). In classical endocytosis, cargos are transferred to early endosomes and sorted in late endosomes (or multi-vesicular bodies for plasma membrane recycling), followed by lysosomal formation and degradation^79^. However, lysosomal inhibition did not impair caveolin1 localization to lipid droplets. Therefore, we concluded that oleic acid induced caveolae endocytosis results in fusion of caveolin1 vesicles with endosomal compartments, followed by caveolin1 localization to lipid droplets (Fig. 8G). In line with the latter, it was shown previously that endosomes and lipid droplets can be in close contact^99–102^. Additionally, lipid and protein trafficking between endosomes and lipid droplets was recently reported^99,102,103^. Future studies are needed to reveal if lipid droplets and endosomal compartments form membrane contact sites similar to lipid droplet – mitochondria or ER contact sites^100^.

To dissect the cellular function of caveolin1 at the lipid droplets we investigated the effect of several caveolin1 mutants on caveolin1-lipid droplet accumulation and lipid droplet growth. Of particular interest is a clinical reported caveolin1 frame-shift mutation resulting in a truncated caveolin1 protein (caveolin1-F160X) and generalized lipodystrophy^38,41,83^. Based on the caveolin1 8S structure^82^ we used Alphafold to model the mutant structure (Fig. 7D).

Interestingly, the central β-barrel, which forms a lipophilic tunnel from the caveolin1 8S membrane-facing side to the cytosolic side of the oligomer^82^, is completely removed in caveolin1-F160X. It was previously discussed that the β-barrel may contain and/or transfer lipid molecules^8,82^. Here, we now show that the loss of the caveolin1 8S β-barrel results in a complete loss of lipid induced caveolin1 trafficking to lipid droplets which results in a significantly impaired lipid droplet growth although new lipid droplets are formed. To further dissect this mechanistically, we also tested a caveolin1 mutation in which the entry of the β-barrel from the membrane-facing side is sterically hindered (caveolin1-R171P). Surprisingly, trafficking to lipid droplets was enhanced (even in absence of oleic acid) but lipid droplets were not increasing in size (Fig. 7, Fig.8) suggesting that lipid transfer to lipid droplets may be impaired. All mutants localized to the plasma membrane prior to oleic acid treatment (Fig. S16) suggesting correct trafficking to the plasma membrane. However, we did not assess mutant oligomerization state or correct caveolae formation by EM, therefore we cannot exclude entirely that assembly defects may contribute to the observed phenotypes. Taken together, this suggests that caveolin1, and more specifically its β-barrel, is needed for the lipid trafficking to lipid droplets and that blocking the entry of the caveolin1 β-barrel results in decreased lipid transfer by the caveolin1 complex to lipid droplets. This reveals for the first time the function of caveolin1 at lipid droplets and why caveolin1 gene mutations may result in such severe lipodystrophy phenotypes. Notably, we did not directly quantify fatty acid uptake or triglyceride synthesis, our conclusions are based on increased lipid droplet growth (and numbers) which only indirectly measure lipid accumulation. However, previous studies showed that lipid droplet sizes correlate with lipid accumulation^104^ and that caveolin1 expression regulates lipid uptake as shown extensively in the past^3^.

Future research is needed to elucidate how fatty acids and cholesterol (and maybe other lipid species) may be transferred through the caveolin1 β-barrel, which other caveolin1 amino acid residues are involved and how these lipids are then processed at the lipid droplet. Here, first evidence can be obtained from the split-APEX proteomics data set of caveolin1-positive lipid droplets (Fig. S10) that showed an enhanced accumulation of long-chain-fatty-acid CoA ligases (ACSL4 and 6) in close proximity to caveolin1. ACSL converts free fatty acids (such as oleic acid) to fatty acid-CoA esters, which are precursor molecules for triglycerides stored in lipid droplets. This suggests that fatty acids may be transferred via the caveolin1 β-barrel to the lipid droplet coat where ACSL activates the fatty acids for further processing resulting ultimately in formation of neutral lipids and lipid droplet growth. In addition, a current open question is to elucidate where the lipids that are needed for lipid droplet growth are coming from: are these lipids already bound to caveolin1 within the caveolin1 β-barrel prior to trafficking to the lipid droplets, or does the caveolin1 β-barrel serve as a tunnel to fuel lipids from endosomes to lipid droplets? In unilocular white adipocytes we also can envision a caveolin1 mediated direct lipid transfer from the plasma membrane to lipid droplets. Future studies are needed to understand this process in its entirety.

## METHODS

### Cell culture

Wildtype or EHD2 KO mouse embryonic fibroblast (MEF) cells (previously reported^30^) were cultured according to standard cell culture practice. In short, cells were cultured in standard T75 flask in DMEM complete medium containing: DMEM (Gibco, 11504496), 10% FBS (Bio&Sell FBS.S.0615), 1% pen/strep (Gibco, 11548876). Cells were split at 80-90% confluency by aspirating medium and rinsing once with 6 mL DPBS. Then 3 mL prewarmed trypsin-EDTA 0.3% (Gibco, 11560626) was added, and cells were incubated with trypsin for 2 minutes at 37 °C. 7 mL prewarmed DMEM complete medium was added, the flask was rinsed, and cell suspension was transferred to a 15 mL falcon tube. Cells were pelleted by centrifugation (1000xg, 4 minutes, room temperature). The medium was aspirated, and the pellet was dissolved in fresh DMEM complete medium. 10 µL of the cell suspension was mixed with 10 µL tryphan blue (Gibco, 15-250-061) and cells were counted using a DeNovix CellDrop BF Cell Counter. 8.000 – 12.000 cells were seeded in µ-Slide 8 Well High (Ibidi, STEM-M-0204-LC) pre-coated with fibronectin (1:1000 fibronectin/DPBS, Sigma F1141, overnight, 4 °C). Cells were incubated at 37 °C under 5% CO2 and humidified atmosphere until further experiments.

3T3-L1 preadipocytes cells were cultivated in T75 flasks in complete DMEM medium (Gibco, 11504496) containing 10% FBS (Bio&Sell FBS.S.0615) and 1% pen/strep (Gibco, 11548876) at 37°C in a humidified atmosphere containing 5% CO2, for at least 3 days prior to induction. Differentiation was induced at 70% confluency, by replacing the complete medium with differentiation medium consisting of complete Medium supplemented with 10 µg/ml Insulin (Sigma, I9278), 500 µM IBMX (Sigma, I7018), 2,5 mM dexamethasone (Sigma, D2915) and 25 mM rosiglitazone (Sigma, R2408) (as previously described^30^) for 3 days with daily medium changes. On day 4 of differentiation, medium was changed to complete DMEM medium supplemented with 1 µg/ml Insulin (Sigma-Aldrich, I9278). For experiments, cells were either expanded in T150 flasks or seeded into fibronectin- coated 8-Well ibidi µ-slides and cultured until day 10 of differentiation, when they were processed for lipid droplet isolation or imaging.

### Plasmid transfection

Transfection was carried out using lipofectamine3000 (Invitrogen, L3000008) according to standard manufacturers protocol. In short, for transfection of 1 well in an µ-Slide 8 well ibidi, 1 µL lipofectamine 3000 reagent, 0.5 µg plasmid DNA and 1 µL P3000 reagent were complexed in opti-MEM medium and added dropwise to the cell, which were incubated 24-48h at 37°C, 5% CO2 before further experiments. The following plasmids were used: Caveolin1-eGFP^1^, Caveolin1-mApple (kind gift from Justin Taraska, NIH), Caveolin2-eGFP^1^, pEGFP-EHD2^1,30^, Rab5-EGFP, Rab5Q79L-EGFP, Rab5N133I (kind gift from Volker Haucke), Cav1R171P-EGFP, Cav1E177V-EGFP and Cav1F160X-EGFP mutants were cloned via PCR (primers: Cav1R171P-EGFP forward: 5’-gtttctttctgcatgttgatggGgatattgctgaatattttgc-3’, reverse: 5’-gcaaaatattcagcaatatccCcatcaacatgcagaaagaaac-3’. Cav1E177V-EGFP forward: 5’-gggcccgcggtgttActttctgcatgttg-3’, reverse 5’-caacatgcagaaagTaacaccgcgggccc-3’, Cav1F160X-EGFP primer forward: 5’-ACCCGTTCCCGCGGGCCCGGGATCCA-3’, primer reverse: 5’-CCCGCGGGAACGGGTCACAGAAGGTGT-3’). PCR cloning was carried out using the Platinum Superfi II DNA polymerase (Invitrogen, 12361010) according to the manufacturers’ protocol.

### Lipid treatment and preparation of OA-BSA complex

Oleic acid solution was prepared by mixing 1 mL prewarmed serum-free DMEM (Gibco, 11504496) with 1µg/ml insulin (Sigma, I9278) and 100 µM - 2.4 mM oleic acid (Sigma, O1383). This solution was very shortly vortexed. Cells were washed once with DMEM, after which the oleic acid dilution was added to the cells. Cells were incubated from 30 minutes up to 6 h at 37 °C, with oleic acid solution. 0.5, 1 or 2.4 µM Glyceryl trioleate (Sigma, T7140) or 10 µM cholesterol (Sigma, C8667) solutions were prepared as described for oleic acid, followed by 1-6h treatment on MEFs. Oleic acid-BSA complex was prepared by first dissolving 0.99 g fatty acid free BSA (Sigma, 126609) in 5 ml PBS (Carl Roth, 0890.1) to obtain a 3 mM BSA stock solution. Aliquots of 1320 µl from the 20 % BSA stock were incubated at 37°C. For the 10 mM oleate stock, 5 ml PBS was supplemented with 50 µl 1 N NaOH (Carl Roth, K021.1) and heated to 70°C. Oleic acid (14.3 mg; Sigma, O1383) was added and incubated for 30 min at 70°C with intermittent vortexing. Subsequently, 20 µl NaOH was added and incubation continued until a clear solution was obtained. The oleate/BSA complex (2:1 molar ratio) was prepared by mixing 800 µl of oleate solution (prewarmed to 70°C) with 1320 µl of 20% BSA (prewarmed to 37°C) in a glass vial (Carl Roth, XC41.1), resulting in a final concentration of 3.8 mM OA and 1.89 mM BSA. To obtain a 1 mM oleate-BSA solution, 1989 µl of the OA/BSA complex was diluted with 5511 µl of prewarmed serum-free DMEM (Gibco, 11504496). BSA control solution was prepared analogously, replacing oleate with PBS during complexation. Oleate-BSA solutions were prepared and stored at 4 °C for a maximum of 2 weeks. Before oleate-BSA was added to the cells, they were washed once with serum-free DMEM, after which the oleate-BSA solution was added, and cells were incubated for 3h at 37 °C.

### Inhibitor experiments

Stock solutions were prepared for the following inhibitors: 100 µM bafilomycin A1 (Sigma 196000, LOT 3919549) in DMSO (ITW reagents, A3672), 5 mM u18666a (Sigma, 662015, LOT 4191126) in dH2O (Carl Roth, T143), 2 mM vacuolin-1 (Sigma, 673000, LOT 4196531) in DMSO, 2 mM Vps34-IN-1 (Biozol, MCE-HY-12795, LOT614101) in DMSO, 2 mM SAR405 (Sigma, 5330630001, LOT 4157269) in DMSO, 5 mM Myriocin (Sigma, M1177) in methanol (Carl Roth, KK39.2) and stored according to manufacturer’s instructions. Final inhibitor solution was prepared by mixing 1 mL prewarmed DMEM (Gibco 11504496) with 1 µL insulin (Sigma, I9278) and 2.4 mM oleic acid (Sigma, O1383), and inhibitors were added with final concentrations: 100 nM bafilomycin A1, 2.5 µM u18666a, 1 µM vacuolin-1, 1 µM Vps34-IN-1, 1 µM SAR405, 2.5 µM myriocin (representative controls: DMSO 0.05% or 0.1%, Methanol 0.05% or dH2O). Cells were washed once with DMEM and incubated with the inhibitor solution for 3h at 37 °C.

### Alcohol treatments

Stock solutions of the alcohols were prepared in DMEM (Gibco 11504496): 8.6 mM heptanol (Sigma, 51778), 2.3 mM octanol (Sigma, 297887) and 0.9 mM nonanol (Sigma, 74280). These solutions were vortexed until fully mixed and stored at 4 °C for maximum 2 months. Treatment solutions were prepared by diluting stock concentrations in DMEM to get final concentrations of 2.5 µM heptanol, 0.5 µM octanol or 0.5 µM nonanol. The solutions were shortly vortexed. Cells were washed once with DMEM, after which the alcohol dilution was added to the cells and incubated for 3h at 37 °C.

### Immunofluorescence staining

Cells were seeded, transfected and treated with oleic acid for 30 min, 1h or 3h as described before. Cells were washed 1x with PBS and fixed for 10 min at RT with 4% PFA. Cells were washed 3x with PBS and incubated with a permeabilization and blocking solution of 5% goat serum (Sigma, G9023) and 0.3% Trition-X100 (Carl Roth, 3051.3) in PBS for 1h at room temperature. The blocking solution was removed and primary antibody solution of 1:100 in 5% goat serum/PBS was added to the cells without washing and incubated for 1h at RT. Cells were washed 3x with PBS before adding secondary antibody solution of Goat anti-Rabbit IgG (H+L) Cross-Adsorbed Secondary Antibody-AlexaFluor™594 (Invitrogen, A-11012) or Goat anti-Rabbit IgG (H+L) Cross-Adsorbed Secondary Antibody-AlexaFluor™568 (Invitrogen, A-11011) 1:500 in 5% goat serum/PBS. Cells were incubated at RT for 1h in the dark. Cells were washed 3x with DPBS, lipid droplets were stained as described below. After washing with DPBS the plate was stored at 4 °C in the dark until imaging (fluorescence microscopy was done in PBS). The following primary antibodies were used: anti-caveolin1-rabbit (Abcam, 2916), anti-cavin1-rabbit (Abcam, 76919), anti-Rab5-rabbit (Cell signaling, 3547T), anti-Rab7-rabbit (Cell signaling, D95F2), anti-EEA1-rabbit (Cell signaling, C45B10).

### Lipid droplet staining

Cells were washed 1x with DPBS and fixed for 10 min at RT with 4% PFA (Carl Roth). Next, cells were washed 3x with DPBS and incubated at RT for 20-30 minutes with Nile Red (Sigma, 72485, 1:1000), BODIPY (Invitrogen, D3922, 1:1000) or LipidTox (Invitrogen, H34477 1:500) in DPBS. Cells were washed 3x with DPBS and then DPBS was added to the wells and the plate was stored at 4 °C in the dark until imaging. Imaging was done in PBS at room temperature.

### Preparation of MEF plasma membrane sheets and platinum replica EM

Pt replica of MEF membrane sheets were prepared as described before^1,59^. Briefly, MEFs were seeded on 22 mm coverslips (1.5H, Marienfeld-Superior #0117640, precoated with fibronectin, 1:100), after 24-48h cells were treated with oleic acid or cholesterol as mentioned above, cholesterol depletion was done with 10 mM methyl-beta-cyclodextrin (15 min, at 37°C, diluted in HEPES, Sigma, M7439), followed by unroofing with a 12-gauge syringe with 2% PFA (EM Science) diluted in stabilization buffer (70mM KCl, 30mM HEPES buffer brought to pH 7.4 with KOH, 5mM MgCl2, 3mM EGTA). After fixation with 2% glutaraldehyde (EM Science) for 20 min membrane sheets were treated with tannic acid (EM Science) and 0.1% uranyl acetate (EM Science) for 20 min, followed by an ethanol row from 15% - 100% EtOH (diluted with H2O). Dehydrated membrane sheets were then dried by critical point drying with CO2 (Leica, CPD3000, manual) and a 3nm Pt coat followed by 5-6nm Carbon coat was added in Leica ACE600. The coverslip of the Pt coated membrane sheets was removed with 5% hydrofluoric acid, followed by extensive washing with H2O. Replicas were transferred to Formvar 75 mesh copper grids (Ted Pella #01802-F) grids and inspected by Joel 1400 microscope at 15,000 magnification (pixel size 1.23 nm).

### Correlative fluorescence and FIB-SEM

Glass coverslips etched with grid (Bellco Biotechnology #1916-91012) were washed with 100% ethanol, coated by adding 1 mL of a 1:100 fibronectin (Sigma, F1141) dilution in DPBS and incubating overnight at 4 °C. After removing the fibronectin solution, coverslips were washed once with DPBS and prewarmed DMEM medium was added the coverslips. 100000 cells were seeded, followed by caveolin1-EGFP expression by Lipofectamine3000 according to manufacturers’ instructions. After 48h MEFs were treated with oleic acid as described above, followed by 10 min 4% PFA fixation at room temperature (EM-grade, EM Science) and lipid droplet stain (Nile Red, Sigma, 72485). Confocal microscopy was performed in PBS using a Zeiss LSM880 Airyscan with 63x oil objective (Zeiss, NA) in airyscan super-resolution mode. MEFs expressing caveolin1-EGFP were selected, z-stacks (0.64 µm interval) were acquired of the whole cell volume. Immediately after imaging, coverslips were treated with 2% glutaraldehyde (EM grade, EM Science) for 1h and rinsed in 0,1M cacodylate buffer. Osmification and ultrathin embedding was done as described previously^105^. In short, cells were postfixed with 1% OsO4 and 1,5% K4Fe(CN)6 in 0,1M cacodylate buffer for 40 min on ice, washed in ddH2O, incubated with 0,1% thiocarbohydrazide in ddH2O for 3 min, washed with ddH2O, postfixed with 1% OsO4 in ddH2O for 20 min, rinsed with ddH2O, incubated in 1% aqueous uranyl acetate for 30 minutes and dehydrated in a graded series of acetone (30%, 50%, 70%, 90%, 100%) 10 minutes each, following infiltration in aceton/Durcupan mixes and pure resin incubation (all steps 1 hour), ultrathin embedding was performed. Coverslips were placed on glass slides (cells facing up), submerged into aceton atmosphere in 50 ml falcon for 10 minutes and centrifuged at 1000g. Afterwards coverslips (cells facing up) were placed for polymerization at 60°C for 48-72 hours; mounted onto SEM stabs with conductive epoxy, sputter carbon coated (30-50nm) and imaged at Helios 5CX FIBSEM. Overview and light microscopy stack projections were overlaid with topographic SEM image (ETD secondary electrons) of cells in MAPS software. Electron microscopy 3D stacks were produced using ASV 4.0 software with ion beam settings at 30kV/0,23nA and electron beam at 2kV/0,34nA, 5 µs dwelling time. Images were acquired using inColumn Detector (ICD) at 3,37x4,6 nm x,y resolution, z was set to 10 nm. Stack was aligned using Microscopy image browser and resized to 10nm isovoxel. For fluorescence microscopy and EM stacks alignment, BigWarp plugin from Fiji package was used.

### Lipid droplet isolation and Western Blot

Lipid droplets were isolated from differentiated 3T3-L1 adipocytes as previously described^106,107^. Cells were briefly washed twice with ice-cold DBPS with Ca²⁺/Mg²⁺ (Gibco, 14040174), scraped and pelleted. Cell pellets were resuspended in hypotonic lysis buffer (HLM buffer) containing 40mM Tris-HCL pH7.4, 2mM EDTA, 20mM NaF, cOmplete protease cocktail (Roche, 04693116001) and Halt phosphatase inhibitor cocktail (Thermo Fisher scientific, 78420) and homogenized on ice using a Potter-Elvehjem homogenizer. Post-nuclear supernatants were mixed with OptiPrep (Sigma, D1556) to a final concentration of 30% and overlaid with a discontinuous gradient from 25-0%. Following by ultracentrifugation, at 17200rpm, for 30min, (4°C, SW55Ti rotor, Beckman Coulter) the floating lipid droplet layer was collected, washed three times with HLM-buffer, and resuspended in 1 x HLM. For immunoblotting lipid droplets, cytosolic and pellet fractions were mixed with Laemmli sample buffer containing DTT, heated for 5min at 95°C and resolved on Bolt 8% Bis-Tris Plus gels (ThermoFisher, NW00080BOX). Proteins were transferred to PVDF membrane using methanol-free transfer buffer (Thermo Fisher scientific, 35045). Membranes were incubated with primary antibodies 1:1000 diluted in 5% w/v BSA overnight 4°C, followed by HRP-conjugated secondary antibodies. Signal detection was performed by chemiluminescence (Biorad, #1705061). Antibodies used: anti-Caveolin-1-rabbit (Proteintech 16447-1-AP (LOT 00071311)), anti-Perilipin-2-rabbit (Proteintech 15294-1-AP (LOT 00133249)), anti-GAPDH-mouse (Cell Signaling, D4C6R, 97166S (LOT 7)).

### Membrane fluidity measurement

Instrumentation: TIRF measurements were carried out using a wide-field fluorescent microscope (Zeiss, Axio Observer Z1 inverted fluorescence microscope) upgraded with TIRF module (Visitron). A 561-nm laser (Coherent OBIS, 561-100 LS) was brought to focus at the back focal plane of 100×α-Plan Apochromat NA 1.46 objective (Zeiss) after passing through a quad band excitation filter (Chroma, ZET 405/488/561/640 V2 361439). A quad band dichroic mirror (Chroma, TRF89901v2 ET Quad C203707) was used to reflect the excitation beam to the sample and allow emission to pass through. Total internal reflection is achieved in so-called point TIRF illumination by traversing the focus beam of a 561-nm beam towards the periphery of the back focal plane of the objective lens controlled by the Visitron Imaging Software (Visitron) for the excitation of DiI-C18 (Invitrogen, D3911). The fluorescence was collected through the same objective, passes the dichroic mirror and quad emission filter (Chroma, ZET 405/488/561/640 M-TRF V2 349420) onto an EMCCD camera with 16 μm pixel size (Andor, iXon EM+ DU-897).

Sample preparation: Live cells were incubated in 100-200 nM DiI-C18 0.1% DMSO (Invitrogen, D3911) in HBSS (Gibco, 14025092) for 20-30 min at 37°C. After incubation, cells were washed twice with HBSS, then medium was replaced with phenol-red-free DMEM supplemented with HEPES (Gibco, 21063029) and maintained at room temperature during image acquisition.

Data acquisition: sample was illuminated with excitation intensity of 30 W/cm2 at 561 nm. Image acquisition was performed using Imaging FCS Fiji-plugin to control the camera^108^. A stack of 40,000 frames (96 × 96 pixels after 2 × 2 in-camera pixel binning) was recorded at 250 fps with the following settings: pixel readout speed = 17 MHz, analog-to-digital gain = 3×, vertical clock voltage = 0, vertical shift speed = 0.5 μs/line, EM-gain = 300, baseline clamp = "on", cooling temperature = -80 °C. Image stack was saved as 16-bit TIFF file. Prior to acquisition, the sample was allowed to stabilize on-stage for at least 15 minutes to minimize focus drift. To control for xy-drift, cell edges were included in the field of view.

Analysis: Map of diffusion coefficients was obtained using Imaging FCS (Fiji plugin), in which autocorrelation function was computed with a multiple-τ correlator on every pixel and subsequently fitted with a single-component 2D free-diffusion model to determine per pixel average diffusion coefficients of DiI molecules^109^. Data processing was performed in batch mode on all TIFF stacks, generating intermediate Excel files for downstream analysis. Further processing was carried out using Python notebook. Briefly, the central 50% of the acquired ROI was considered, and an upper cutoff of 20 μm^2^/s was applied to exclude unrealistic fits (on the order of thousands of μm^2^/s) caused by accidental inclusion of background pixels in the selected region.

### SplitAPEX fluorescence labelling

SplitAPEX2 constructs were designed as described in Hung et al. (2016)^64^, whereby perilipin1 was tagged with first part of APEX2 (AP) and caveolin1 was tagged with EX-part of the enzyme (Fig. S6A). The immunofluorescence staining for the APEX experiments were done according to^63,64^. Briefly, cells were seeded in an 8-well plate as described above and transfected as described previously using lipofectamine3000 and 0.5 µg plasmid DNA of both caveolin1-EX and Perilipin1-AP. The cells were treated for 3h with 2.4 mM oleic acid and 1 µg/ml insulin in DMEM. The oleic acid solution was removed from the cells and they were treated with 200 µL of a 500 µM biotin phenol solution (Sigma, SML2135) in prewarmed DMEM complete (DMEM + 10% FBS + 1% Pen/strep) for 30 min at 37 °C. 2 µL of a 100 mM hydrogen peroxide (Sigma, 386790) solution was added to the wells (resulting in a concentration of 1 mM hydrogen peroxide in the well) and incubated for 90 seconds while gently shaking. The cells were immediately washed 3x with a freshly prepared quenching buffer containing 10 mM sodium ascorbate (Sigma, PHR1279), 10 mM sodium azide (Sigma, S2002) and 5 mM (±)-6-Hydroxy-2,5,7,8-tetramethylchromane-2-carboxylic acid (Trolox, Sigma, 238813) in DBPS. Quenching solution was removed, and cells were immediately fixed with 4% PFA for 10 min. For the control wells, one of these components in the reaction was omitted: oleic acid, biotin phenol or hydrogen peroxide. Cells were permeabilized for 20 min in blocking buffer containing 0.1% Tween-20. Cells were blocked for 30 min in a buffer containing 3% BSA (Carl Roth, 3737.3) in PBS. Blocking buffer was removed from the cells and Primary antibody dilution was added without washing. Primary antibody dilution consisted of 1:100 dilution of anti-perilipin1 antibody (Cell Signaling, 9349S) in blocking buffer and was incubated on the cells for 1h at RT. Cells were washed 3x with PBS and incubated with Secondary antibody solution consisting of 1:100 Streptavidin, Alexa Fluor™ 647 conjugate (Invitrogen, S21374) and 1:500 Goat anti-Rabbit IgG (H+L) Cross-Adsorbed Secondary Antibody, Alexa Fluor™ 594 (Invitrogen, A-11012) in blocking buffer. Cells were washed 3x with PBS and staining solution containing 1:1000 BODIPY and 1:1000 DAPI in PBS was added to the cells and incubated 15 min at RT in the dark. Lastly, cells were washed 3x with PBS and PBS was added to the wells. Samples were stored at 4 °C in the dark until imaging. Samples were imaged in PBS at room temperature.

### SplitAPEX Proteomics sample preparation

800 000 MEFs were seeded in a 10 cm well plate. Transfection was done using Dreamfect gold (OZ biosciences, OZB-DG81000) with 5 µg Cav1-EX and 5 µg Pln1-AP plasmid DNA and 30 µL Dreamfect gold reagent. After 6h the transfection medium was replaced with DMEM/10%FBS/1%P/S. After 48h, cells were treated with 2.4 mM oleic acid and 1µg/ml insulin in serum-free DMEM at 37 °C for 3h. Next, cells were treated for 30 min with 5 mL of a 500 µM biotin phenol (Sigma, SML2135) solution, in DMEM at 37 °C. 50 µL of a 100 mM hydrogen peroxide (Sigma, 386790) solution was added to the cells (resulting in a 1 mM hydrogen peroxide concentration) and this was incubated for 90 seconds while gently shaking. The reaction was immediately quenched by adding quenching buffer containing 10 mM sodium ascorbate (Sigma, PHR1279), 10 mM sodium azide (Sigma, S2002), and 5 mM Trolox (Sigma, 238813) in DPBS. This solution was removed and 200 µL APEX lysis buffer was added to the cells (1x RIPA buffer (Sigma, 20-188), 1x protease inhibitor cocktail (Roche, 11836153001), 10 mM sodium ascorbate, 10 mM sodium azide and 5 mM Trolox in DPBS). The plate was put on ice and cell lysates were scraped and transferred to a 1.5 mL tube. Control samples were made by omitting one of the following steps: transfection with cav1-ex and pln1-ap, oleic acid treatment, biotin phenol treatment or hydrogen peroxide treatment. Lysates were homogenized by pipetting and vortexing intermittently for at least 30 minutes while the samples incubated on ice. Lysates were cleared by centrifuging for 15 min at maximum speed and 4 °C. The supernatant was transferred to a new 1.5 mL tube and pellets were discarded. Next, 300 µL of streptavidin resin (Thermo Scientific, 20359) was added to a clean 1.5 mL tube (one per sample) and shortly centrifuged. The supernatant was removed, the resin was washed 2x with the APEX lysis buffer, shortly centrifuged and the supernatant was removed each time. The lysates were added to the resin and incubated overnight at 4 °C while rotating. The resin was washed in different steps: firstly, twice with APEX lysis buffer, then once with 1 M KCl, once with 0.1 M Na2CO3, once with 2M urea in 10 mM Tris-HCl buffer, pH 8 and lastly twice with APEX lysis buffer. After each washing step, samples were centrifuged briefly, and the supernatant was removed using gel loading tips. The proteins were eluted with a solution of 4x protein loading buffer (Thermo scientific J60015.AD), 20 mM DTT (Thermo scientific, A39255) and 10 mM biotin (Sigma, B4501) in miliQ water. This was boiled for 10 min at 95 °C while shaking. The samples were briefly vortexed and centrifuged for 30 min, 16000 RCF, 4 °C. The supernatant was collected and sent for mass spectrometry analysis.

### Mass spectrometry

Proteolytic digestion of SplitAPEX eluates using SP3: SplitAPEX experiments were carried out in five replicates and eluted using 4X SDS-PAGE Loading dye for 10 min at 95 °C and then digested using the SP3 protocol^110^. Proteins were reduced with 5 mM TCEP (Sigma) and alkylated with 40 mM CAA (Sigma) for 1 h at room temperature. Sera-Mag beads were added to the sample, followed by 50% Acetonitrile (ACN, Fisher scientific, MS grade). After a brief incubation the supernatants were removed, and the beads were washed twice with ethanol and once with ACN. Proteins were digested in 50 mM TEAB (pH 8.5, Sigma) with LysC (Fujifilm Wako) and Trypsin (Serva), both in enzyme:protein ratio 1:50 wt:wt, for 16 h at 37 °C. Beads were washed twice with an excess of ACN. Peptides were eluted with 5% DMSO and dried until further use.

Liquid chromatography and mass spectrometry: Peptides were resuspended in 1% ACN with 0.05% trifluoroacetic acid (TFA) and injected into a Thermo Scientific Vanquish Neo system coupled online to an Orbitrap Astral mass spectrometer (ThermoFisher, Tune version 1.1). Prior to separation, peptides were trapped on a PepMap C18 trap column (0.075 × 50 mm, 3 μm, 100 Å, Thermo Fisher). Peptide separation was performed using reverse-phase chromatography on an in-house packed C18 column (Poroshell 120 EC-C18, 2.7 μm, Agilent). Elution was carried out over a 40-minute linear gradient of increasing ACN concentration at a flow rate of 300 nL/min. Data were acquired in data-independent acquisition (DIA) mode. MS1 scans were performed in the Orbitrap at a resolution of 240,000 over a scan range of m/z 430–680, with a 40% RF lens setting, 500% AGC target, and a maximum injection time of 5 ms. MS2 scans were acquired in the Astral analyzer using 2 m/z isolation windows with no overlap, automatic window type, placement optimization enabled, normalized collision energy of 25%, scan range of m/z 145–2000, 40% RF lens, 500% AGC target, 5 ms maximum injection time, and a 0.6-second duty cycle.

Identification and quantification of proteins enriched with Streptavidin: All raw data were combined and analyzed using DIA-NN version 2.0^111^. Spectra were searched against an in silico spectral library generated from the murine proteome (one protein per gene) and common contaminants. DIA-NN was configured with the following parameters: MS1 tolerance of 10 ppm, MS2 tolerance of 20 ppm, peptide length range of 7 – 30, trypsin digestion allowing a maximum of two missed cleavage. Match-between-runs was enabled. Variable modifications included oxidation of methionine (+15.9949 Da) and protein N-terminal acetylation (+42.0106 Da), while carbamidomethylation of cysteine (+57.0215 Da) was set as a fixed modification. All other settings were left default. Quantitation analysis was performed in R. Proteins were retained if quantified in at least three out of four replicates in at least one condition (replicate three has been permanently removed before data processing). Intensity values were log2-transformed, normalized using median-centered normalization. Differential expression analysis was conducted using the limma package to compute log2 fold changes and p-values from an empirical Bayes-moderated t-test.

### Confocal microscopy

All confocal imaging was done using a Zeiss LSM880 confocal microscope equipped with a Plan-Apochromat 63x / 1.4 Oil DIC M27 objective and different lasers: Argon multiline laser 488 nm, HeNe laser 594nm, HeNe laser 633 nm and DPSS 561-10 nm laser. ZEN software (black edition) system 2.3 SP1 (Zeiss) was used for settings. In general, for all confocal imaging z-stacks were acquired spanning the entire cellular volume (from basal to apical plasma membrane, z-stack distance 0.64 µm, set by Zeiss ZEN settings).

### Image analysis

All fluorescence imaging data was inspected and analyzed in Fiji/ImageJ^112^. For lipid droplet analysis a custom Fiji macro was developed (code available in supplemental information). Here, at first maximum projections of z-stacks were generated, lipid droplets were selected using intensity thresholding of the lipid droplet channel. Next, ROIs were created and slightly enlarged for every lipid droplet. These ROIs were applied in the caveolin1 channel and size (in µm^2^) and fluorescence intensities (Integrated density, AU) were measured. If not otherwise indicated representative confocal images represent maximum projections to visualize all lipid droplets throughout the cell volume. Plasma membrane localization of caveolin1 mutants were analyzed on basal membrane plane of acquired z-stack, whereby fluorescence intensity was normalized to total cellular membrane area.

PREM images were inspected with Fiji, and segmentation and analysis were done as described before^1^. Briefly, all caveolae found in a TEM image were associated to either low, medium and highly curved group, followed by size measurements^1^ as depicted in Fig S2. Total caveolae number/membrane sheet area was counted and percentage of low, medium and highly curved caveolae was calculated. Similar analysis was applied to clathrin structures according to previously published analysis^61^.

### Molecular modeling

Modeling of caveolin1 mutant structures was done using Alphafold v3^113^ and visualization was done with UCSF ChimeraX 1.10^114^.

### Statistical analysis

Statistical analysis was done using GraphPad Prism 10. Data was tested for normal distribution (Gaussian). If data had a normal distribution, either Welsh’s t-test (for comparing 2 conditions) or ordinary ANOVA (for comparing multiple conditions) was performed to test for significant differences. If data was not distributed normally, Mann-Whitney test (for comparing 2 conditions) or Kruskal-Wallis multiple comparison test (for comparing multiple conditions) was used to test for significant differences. *p≤0.05; **p≤0.01; ***p≤0.001 and ****p≤0.0001.

### Data availability

All resource data are available in supplemental information. The mass spectrometry data have been deposited to the ProteomeXchange Consortium via the PRIDE^115^ partner repository with the identifier PXD069518.

### Code availability

Custom Fiji macros are found in supplemental information. Membrane fluidity python script is published here: https://github.com/danielaik/imfcs-output-handler

## ACKNOWLEDGMENT

We thank the Biology and Chemistry department of University Potsdam for support in light and electron microscopy. C.M. is supported by Chan Zuckerberg Initiative (#2023–331950), Deutsche Forschungsgemeinschaft (German Research Foundation, project grant: 531499831), and Else-Kröner-Fresenius-Stiftung (Project grant: 2023_EKEA.152).

## AUTHOR CONTRIBUTION

C.M., M.L., and E.O. designed and discussed the experiments. If not otherwise indicated E.O. performed and analyzed all experiments. D.A. performed and analyzed membrane fluidity measurements, T.P. performed oleic acid-BSA and insulin experiments. M.Rath supported cloning of plasmids, conducted all 3T3-L1 experiments and Western Blot. C.M. performed and analyzed PREM, M.Ruwolt performed mass spectrometry experiments and data analysis, C.M., L.W.D. and D.P. performed and analyzed correlative FIB-SEM. C.M. and E.O. wrote the paper with input from all authors.

## SUPPLEMENTAL FIGURES

**Figure S1:**
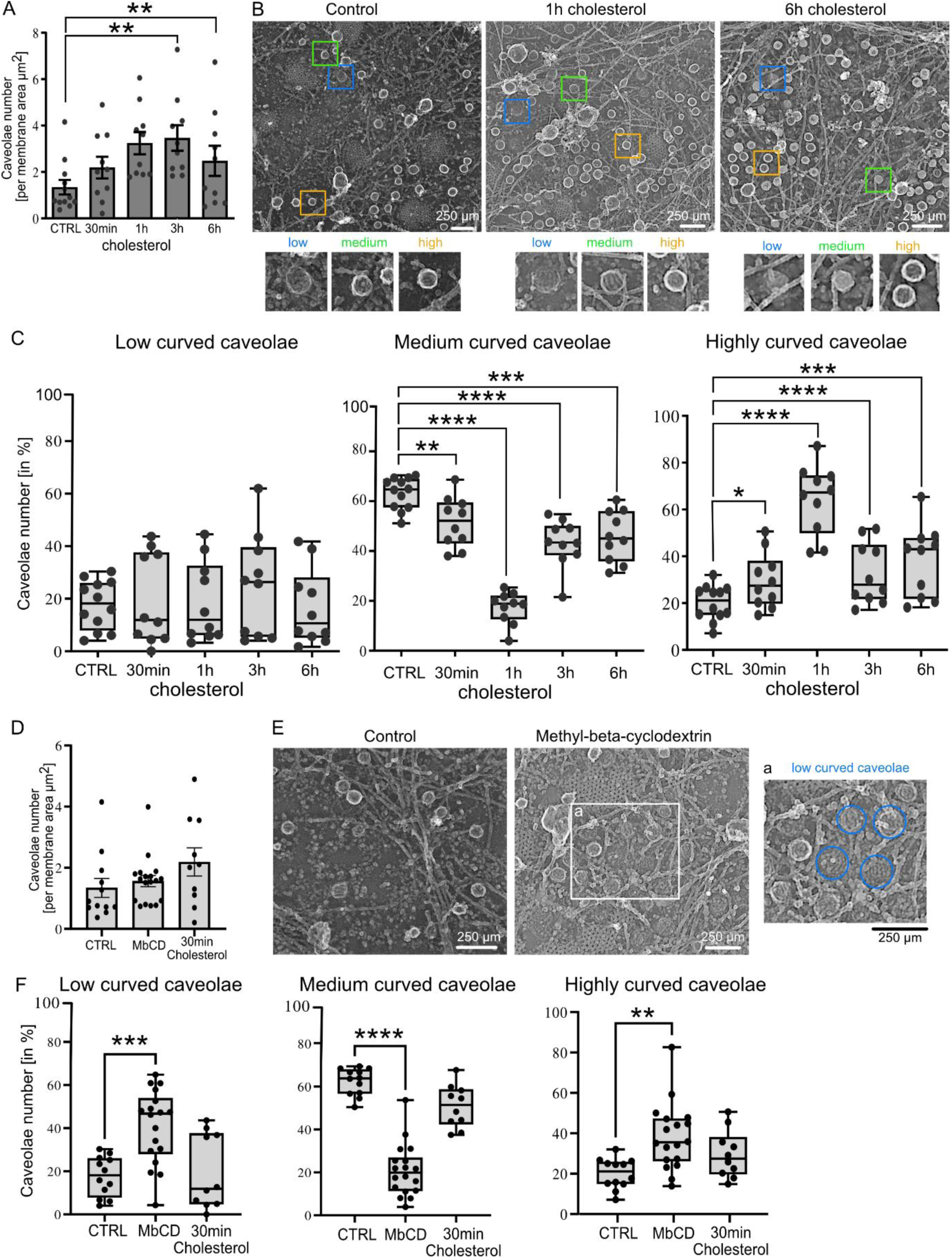
Extracellular cholesterol treatment shifts caveolae to spherical curvature. A-B) MEFs were treated with cholesterol for 30 min, 1, 3, or 6 h and were subsequently unroofed and platinum replica were prepared. Total caveolae number per plasma membrane area is shown (A, bar plot illustrates mean ± SEM, each membrane sheet is depicted). Caveolae in platinum replica EM images were segmented in low, medium, highly curved (spherical) groups (B, scale bar is 250 nm). C) Box plots depict percentage of low curved (C), medium curved (D) or highly curved caveolae per cell (box shows min to max, line represents median, each membrane sheet is depicted). D-E) MEFs were treated with 10 mM methyl-beta-cyclodextrin (MbCD) for 15 min and were subsequently unroofed and platinum replica were prepared. Caveolae number per plasma membrane area is depicted (D, bar plot illustrates mean ± SEM, each membrane sheet is depicted). Caveolae in platinum replica EM images were segmented in low, medium, highly curved (spherical) groups (E, scale bar is 250 nm). F) Box plots illustrate percentage of low curved (C), medium curved (D) or highly curved caveolae per cell (box shows min to max, line represents median, each membrane sheet is represented). For all graphs: n(CTRL) = 12 membrane sheets/ 7 cells, n(30min) = 10 membrane sheets/5 cells, n(1h) = 10 membrane sheets/6 cells, n(3h) = 10 membrane sheets/5 cells, n(6h) = 10 membrane sheets/5 cells, n(M-b-CD) = 10 membrane sheets/5 cells, 2 independent experiments, tested for significant differences with Man Whitney test, *p≤0.05; **p≤0.01; ***p≤0.001 and ****p≤0.0001.

**Figure S2:**
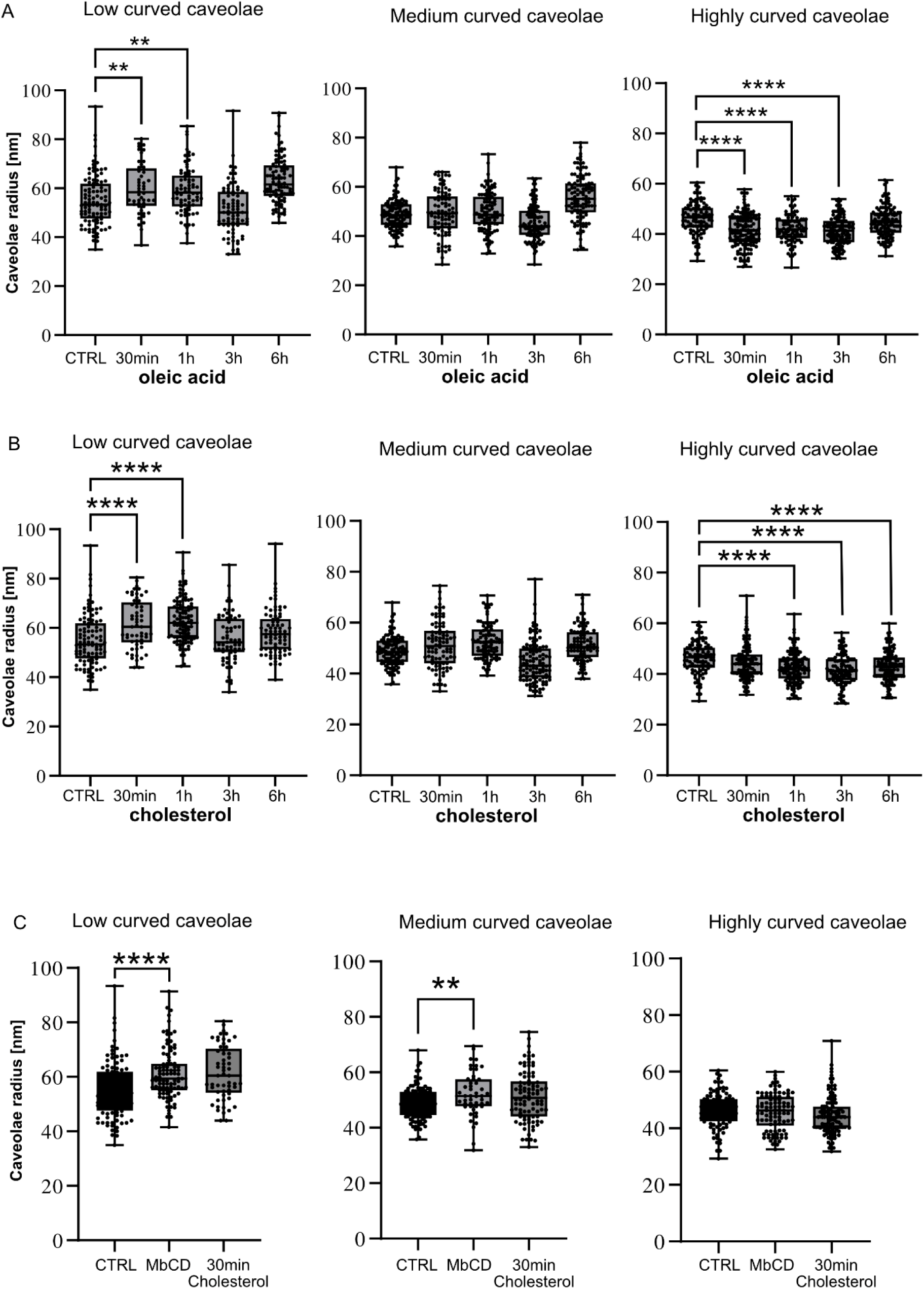
Overview about caveolae size changes after oleic acid, cholesterol, or methyl-beta-cyclodextrin treatment. A) Caveola radius was measured as previously described^1^. Box plots represent caveola radius for oleic acid treatment (box shows min to max, line represents median, each caveola is depicted, low curved: n(CTRL) = 100, n(30min) = 48, n(1h) = 74, n(3h) = 78, n(6h) = 96, medium curved: n(CTRL) = 113, n(30min) = 86, n(1h) = 103, n(3h) = 109, n(6h) = 107, highly curved: n(CTRL) = 96, n(30min) = 132, n(1h) = 91, n(3h) = 126, n(6h) = 120). B) Box plots represent caveola radius for cholesterol treatment (box shows min to max, line represents median, each caveola is depicted, low curved: n(CTRL) = 100, n(30min) = 57, n(1h) = 108, n(3h) = 65, n(6h) = 80, medium curved: n(CTRL) = 113, n(30min) = 92, n(1h) = 88, n(3h) = 109, n(6h) = 86, highly curved: n(CTRL) = 96, n(30min) = 115, n(1h) = 124, n(3h) = 106, n(6h) = 117). C) Box plots represent caveola radius for methyl-beta-cyclodextrin (MbCD) treatment (box shows min to max, line represents median, each caveola is depicted, low curved: n(CTRL) = 100, n(30min) = 57, n(MbCD) = 95, medium curved: n(CTRL) = 113, n(30min) = 92, n(MbCD) = 48, highly curved: n(CTRL) = 96, n(30min) = 115, n(MbCD) = 94). For all graphs: 2 independent experiments, tested for significant differences with Mann-Whitney test, **p≤0.01; and ****p≤0.0001.

**Figure S3:**
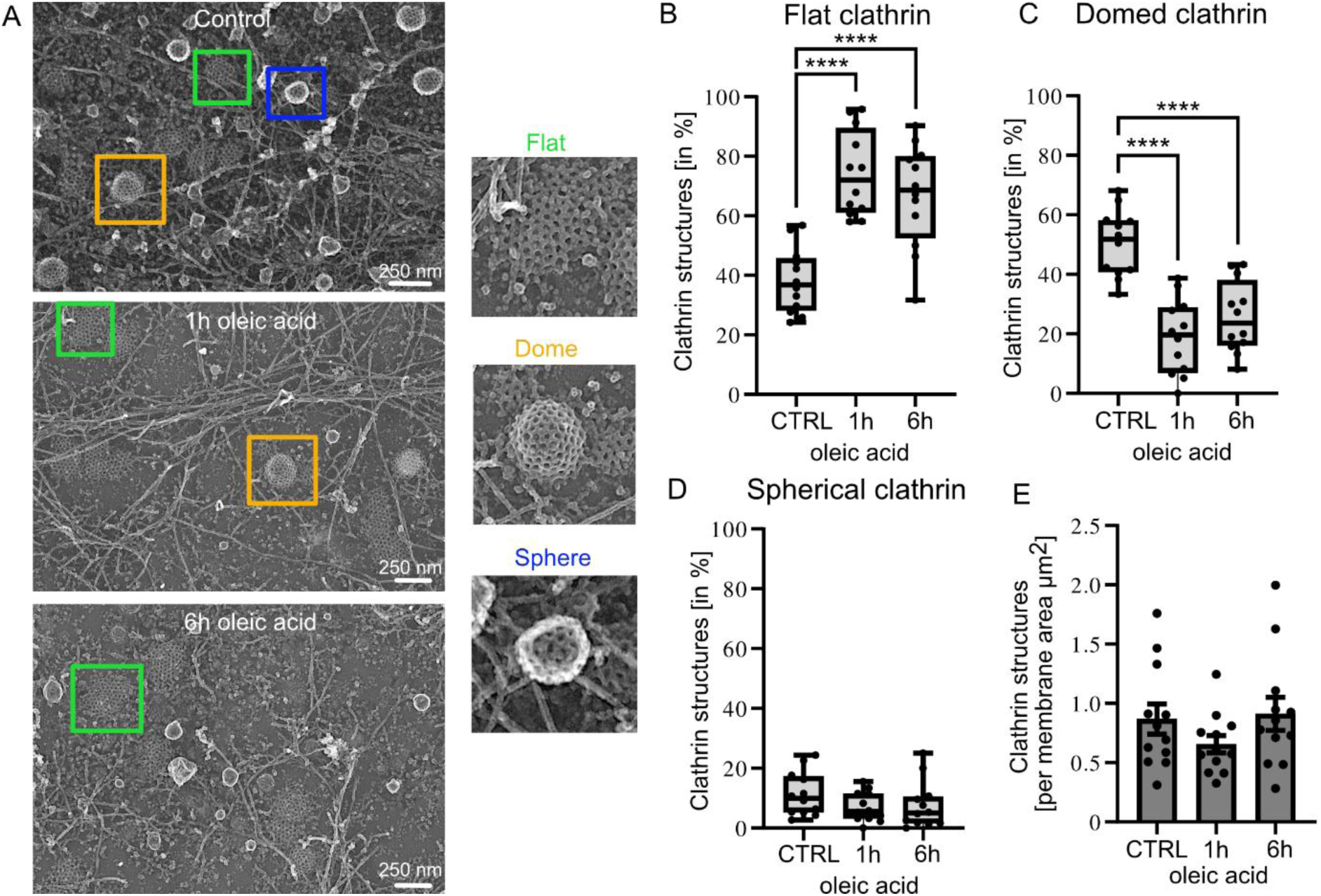
Oleic acid treatment significantly increased flat clathrin structures in MEFs. A) MEFs were treated with oleic acid for 1 or 6 h and were subsequently unroofed and platinum replica were prepared. Clathrin structures were segmented in platinum replica EM images as described before^61^ in flat, dome, or sphere (scale bar is 250 nm). B-D) Box plots depict percentage of flat clathrin structures (B), dome (C) or spheres (D) per cell (box shows min to max, line represents median, each membrane sheet is depicted). E) Clathrin structure number per plasma membrane area is represented (bar plot illustrates mean ± SEM, each membrane sheet is depicted). For all graphs: n(CTRL) = 12 membrane sheets/7 cells, n(1h) = 12 membrane sheets/8 cells, n(6h) = 12 membrane sheets/8 cells, 2 independent experiments, tested for significant differences with Man-Whitney test, ****p≤0.0001.

**Figure S4:**
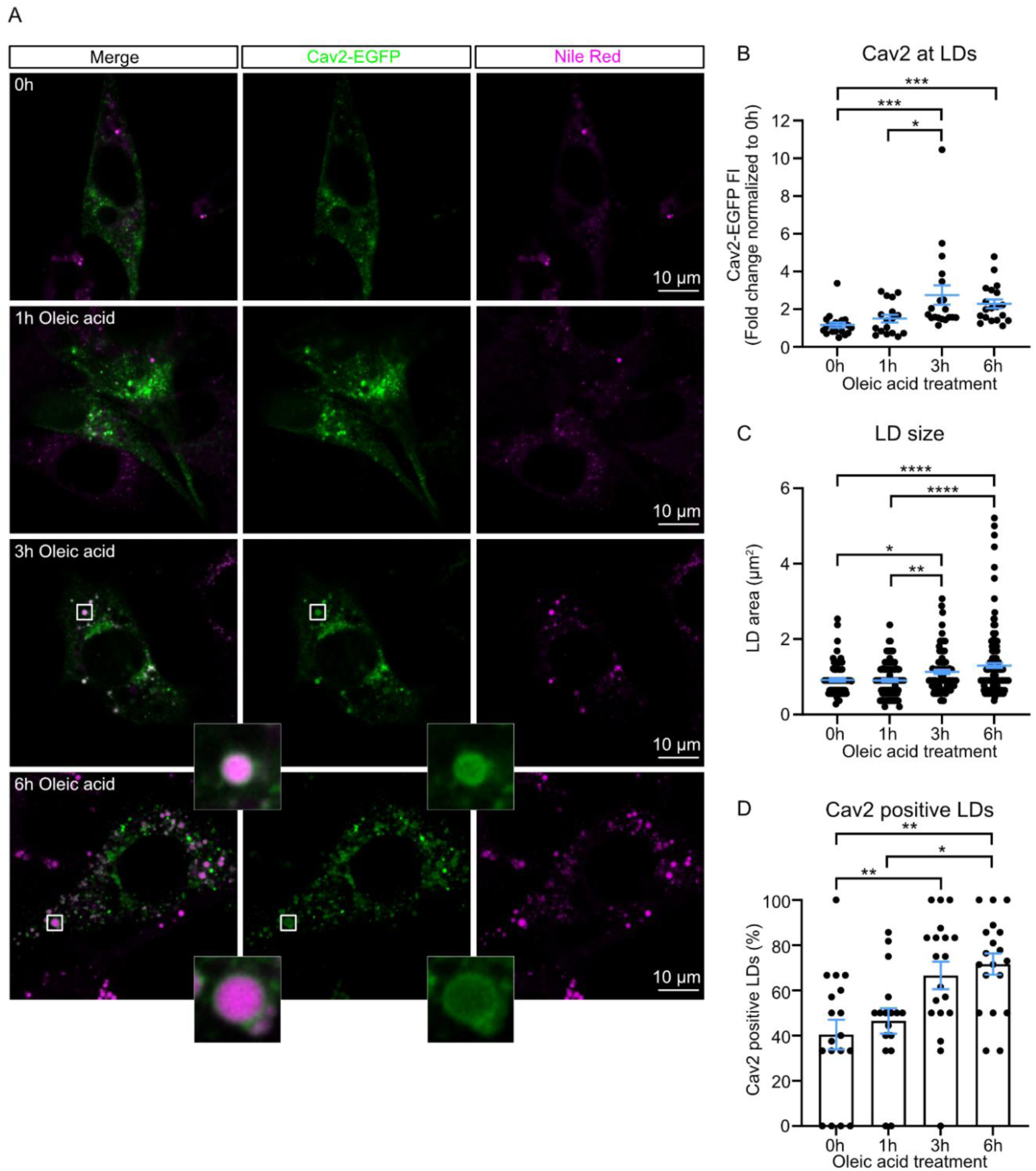
Oleic acid treatment triggers caveolin2 accumulation at lipid droplets. (A) Representative confocal microscopy images of MEFs transfected with caveolin2-EGFP (green), treated with oleic acid for 1, 3, or 6h. Lipid droplets were stained with Nile Red (magenta). Scale bar is 10 µm. (B) Plot depicts normalized caveolin2-EGFP fluorescence intensity (FI) measured at lipid droplets, each spot represents averaged fluorescence intensities per cell (n(0h) = 20, n(1h) = 17, n(3h) = 19, n(6h) = 19). (C) Lipid droplet (LD) area in µm^2^, each spot represents a single LD (mean ± SEM, (n(0h) = 85, n(1h) = 105, n(3h) = 101, n(6h) = 169). (D) Percentage of lipid droplets positive for caveolin2 relative to the total number of lipid droplets/cell (n(0h) = 18, n(1h) = 17, n(3h) = 19, n(6h) = 19). For all graphs mean ± SEM is plotted, minimum 5 cells/experiment, 3 independent experiments, tested for significant differences with multiple comparison Kruskal-Wallis test (panels B and C) or one-way ANOVA (panel D), *p≤0.05; **p≤0.01; ***p≤0.001 and ****p≤0.0001).

**Figure S5:**
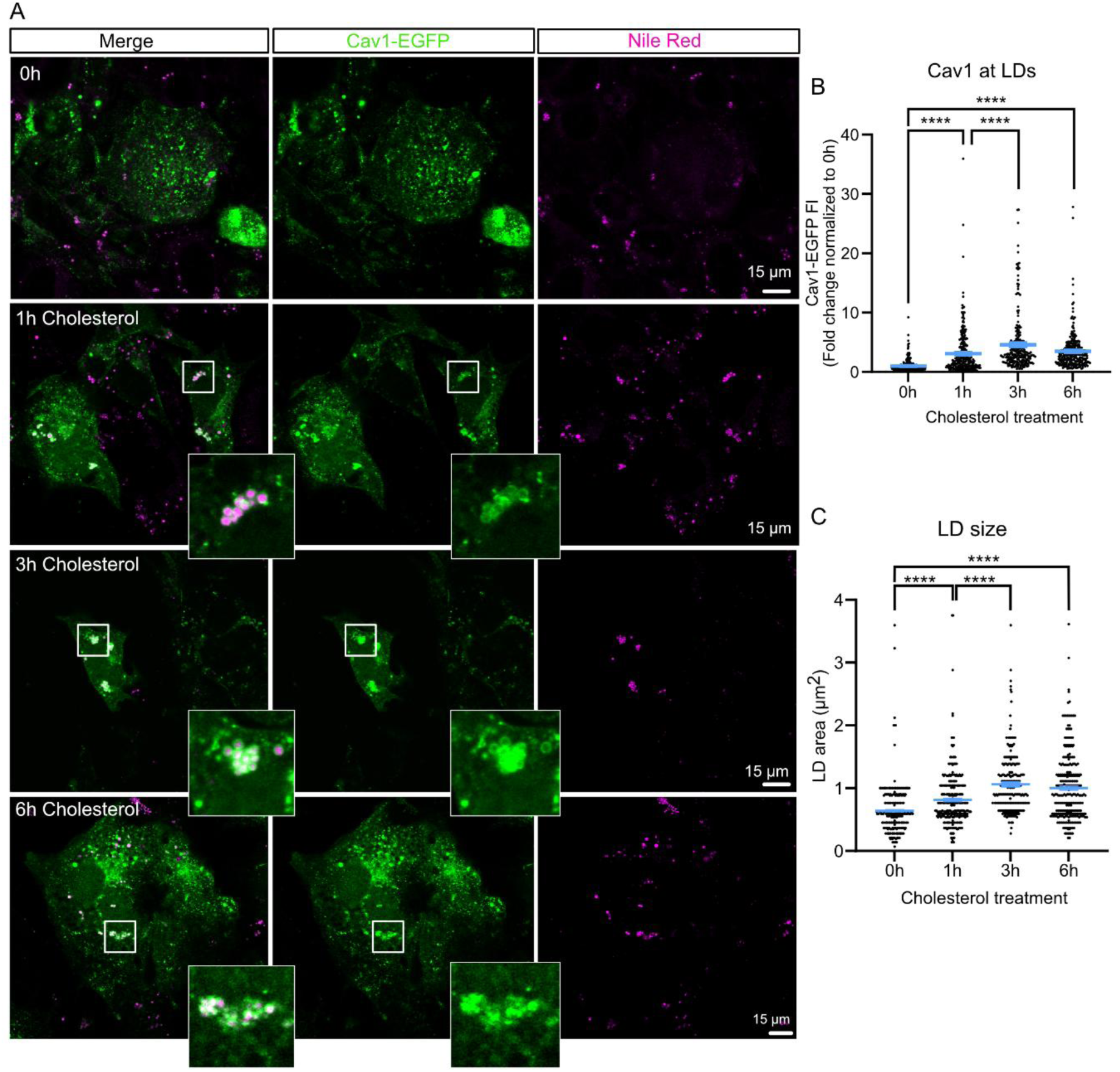
Cholesterol treatment triggers caveolin1 accumulation at lipid droplets. (A) Representative confocal microscopy images of MEFs transfected with caveolin1-EGFP (green), treated with cholesterol for 1, 3, or 6h. Lipid droplets were stained with Nile Red (magenta). Scale bar is 15 µm. (B) Plot depicts normalized caveolin1-EGFP fluorescence intensity (FI) measured at lipid droplets, each spot represents a single lipid droplet (n(0h) = 353/23 cells, n(1h) = 310/18 cells, n(3h) = 249/17 cells, n(6h) = 280/15 cells). (C) Lipid droplet (LD) area in µm^2^, each spot represents a single LD (n(0h) = 353, n(1h) = 310, n(3h) = 249, n(6h) = 280). For all graphs mean ± SEM is plotted, minimum 5 cells/experiment, 3 independent experiments, tested for significant differences with Mann-Whitney test, ****p≤0.0001).

**Figure S6:**
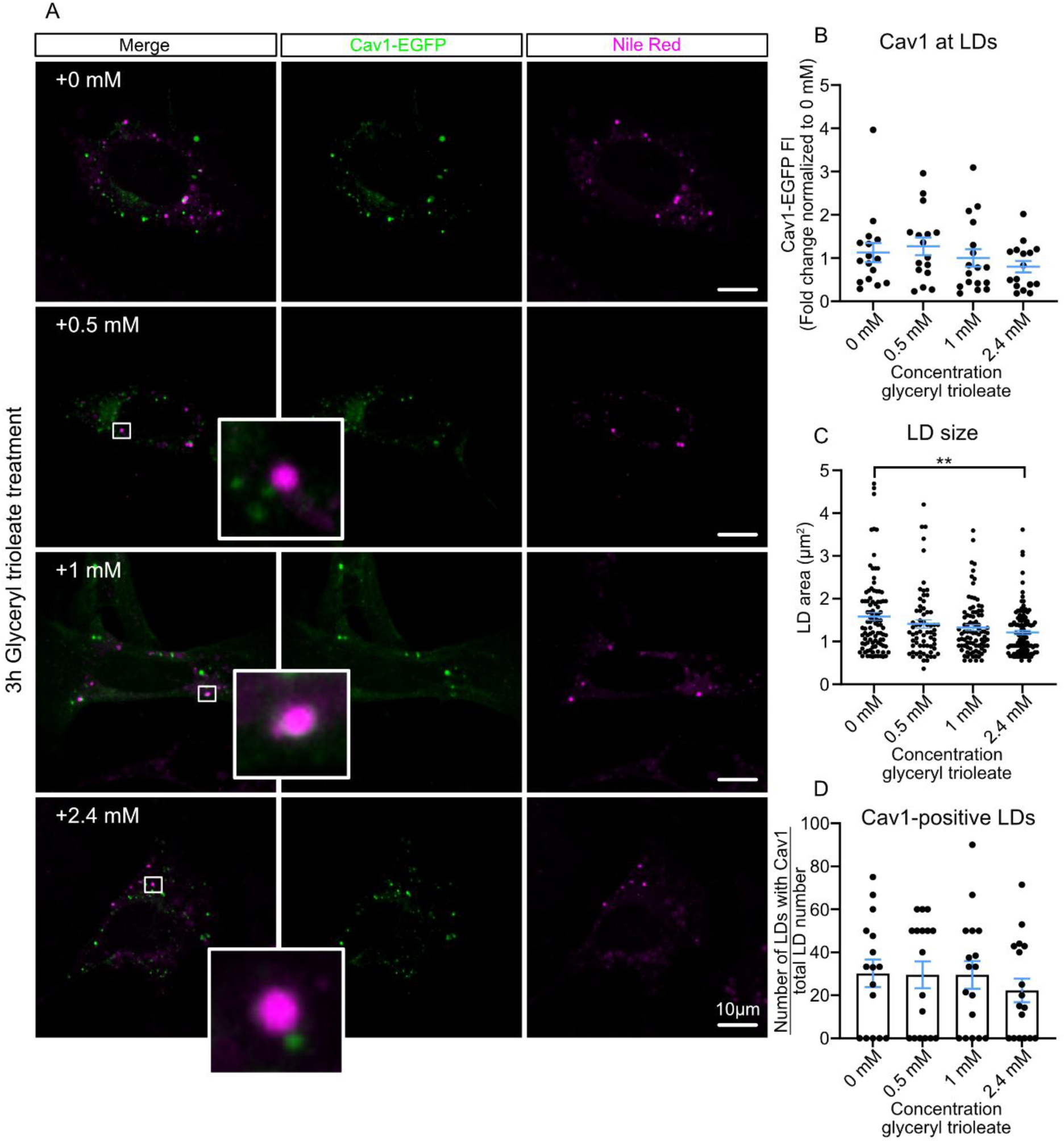
Glyceryl trioleate treatment does not trigger caveolin1 accumulation at lipid droplets. (A) Representative confocal microscopy images of MEFs transfected with caveolin1-EGFP (green), treated with 0.5, 1 or 2.4 mM glyceryl trioleate for 3h. Lipid droplets were stained with Nile Red (magenta). Scale bar is 10 µm. (B) Plot depicts normalized caveolin1-EGFP fluorescence intensity (FI) measured at lipid droplets, each spot represents averaged fluorescence intensities per cell (n(0mM) = 16, n(0.5 mM) = 16, n(1 mM) = 17, n(2.4 mM) = 16). (C) Lipid droplet (LD) area in µm^2^, each spot represents a single LD (n(0mM) = 96, n(0.5 mM) = 67, n(1 mM) = 88, n(2.4 mM) = 112). (D) Percentage of LDs positive for caveolin1 relative to the total number of LDs per cell (n(0mM) = 16, n(0.5 mM) = 17, n(1 mM) = 17, n(2.4 mM) = 17). For all graphs, mean +- SEM, 5 cells/experiment, 3 independent experiments, tested for significant differences with multiple comparison Kruskal-Wallis test, **p≤0.01.

**Figure S7:**
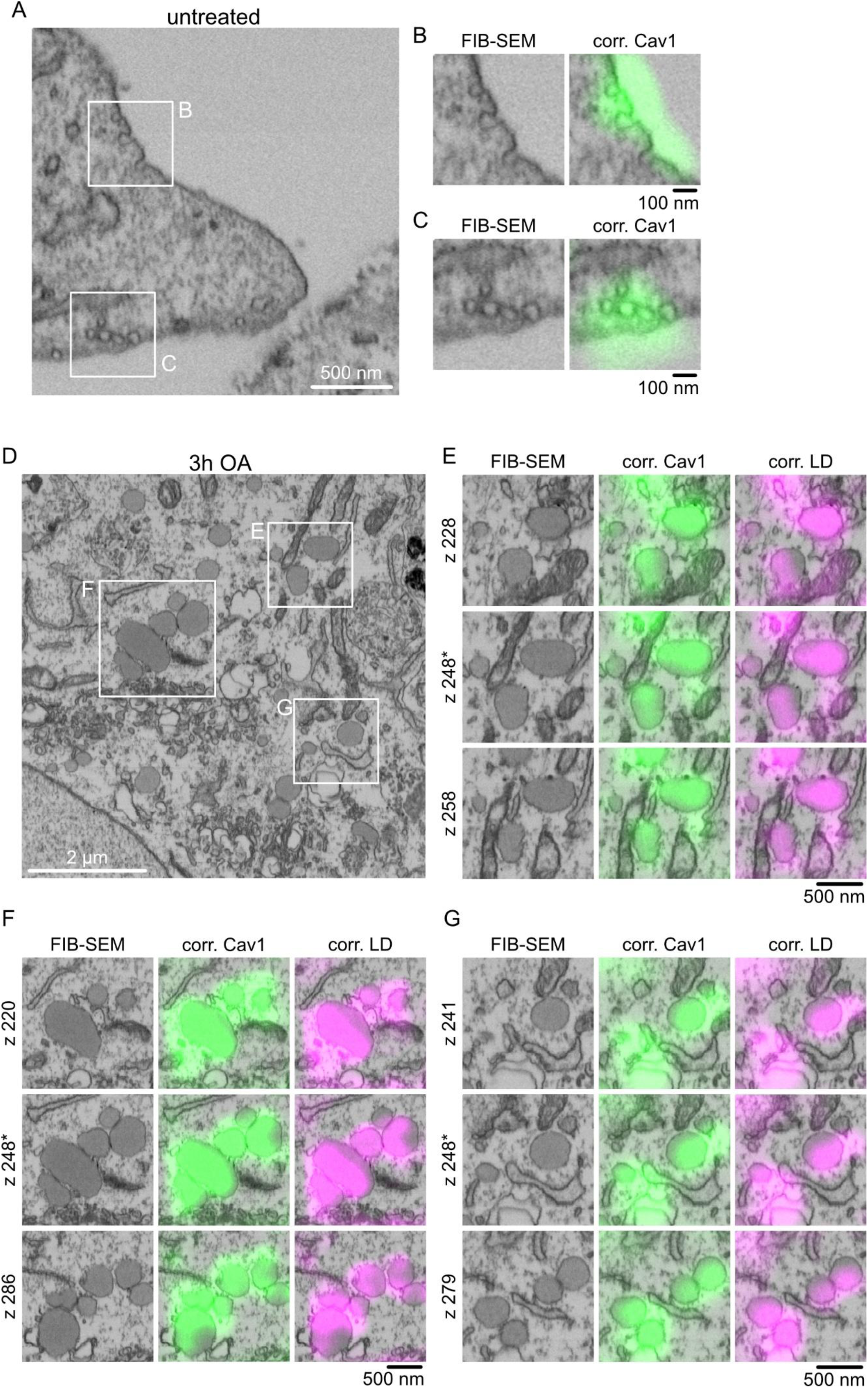
Detailed structural analysis of caveolin1 positive lipid droplets by correlative FIB-SEM. (A-C) FIB-SEM section of MEF expressing caveolin1-EGFP, inset B and C showing caveolae at the plasma membrane with correlated caveolin1-EGFP fluorescence (scale bar is 500 nm for A, scale bars are 100 nm for insets). See also supplemental video S1. (D) FIB-SEM section of MEF expressing caveolin1-EGFP treated with oleic acid (OA) for 3h. Lipid droplets are indicated in inlays B-D, scale bar is 2 µm. (E-G) Enlarged areas from (A) depict lipid droplets in FIB-SEM image correlated to caveolin1-EGFP fluorescence (green) and lipid droplet stain (magenta, Nile Red). For each tile section 3 slices of FIB-SEM volume are shown (z220 – z286, * indicates central FIB-SEM slice, depicted in A) to indicate lipid droplet volume. See also supplemental videos 2-4. Scale bars are 500 nm.

**Figure S8:**
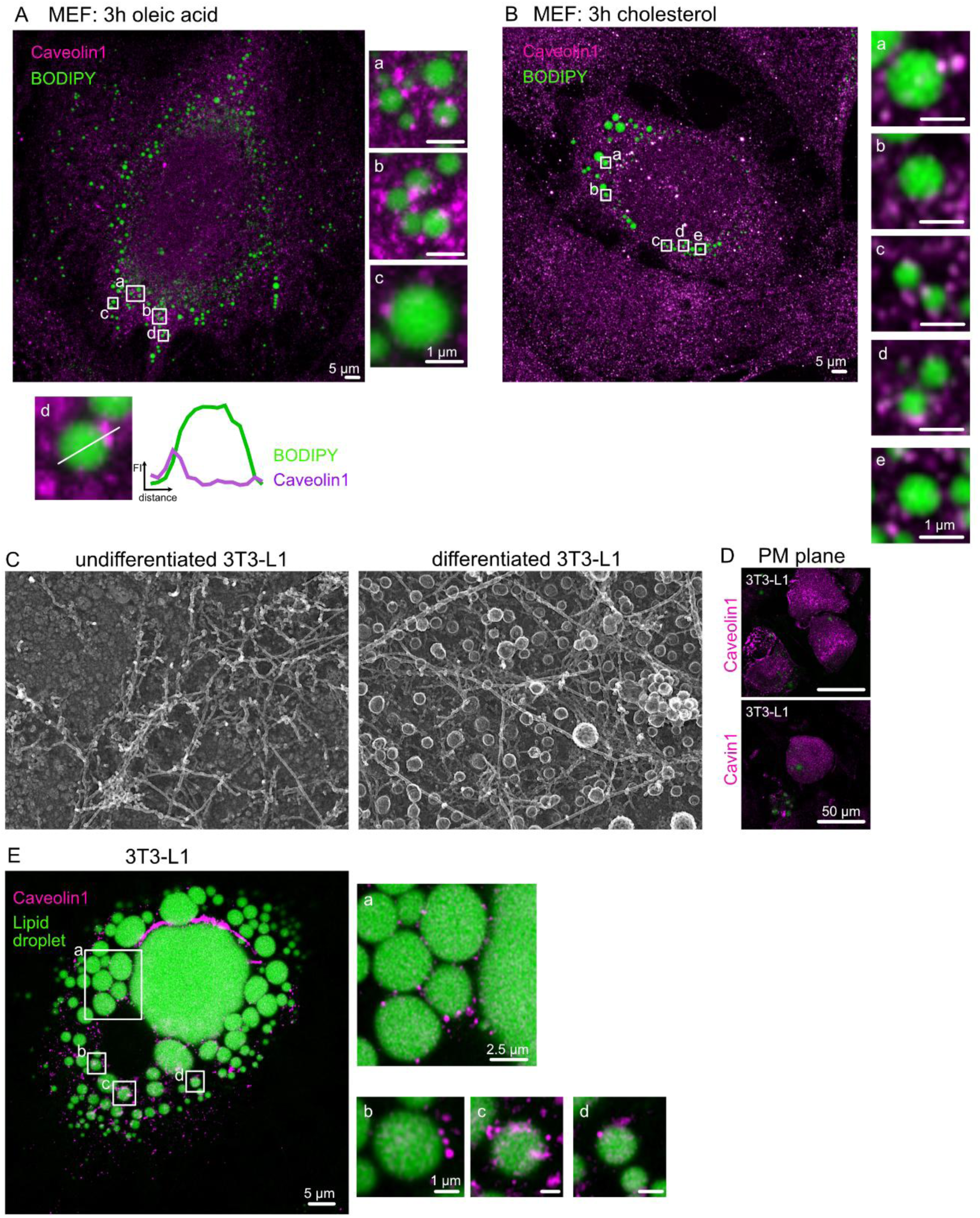
Endogenous caveolin1 accumulation at lipid droplets in MEFs and 3T3-L1 adipocytes. (A-B) Representative confocal images of MEFs treated with oleic acid (A) or cholesterol (B) for 3h, followed by immunostaining with specific antibody against caveolin1 (labelled with secondary antibody tagged with AlexaFluor594, magenta) and BODIPY (lipid droplets, green), scale bars are 5 µm. Numbered white rectangles in (A) or (B) are depicted enlarged in insets, scale bars are 1 µm. Line scan analysis (white line) of inset 4 from (A) depict fluorescence plot profile for BODIPY (green) and caveolin1 (magenta). (C) Representative platinum replica TEM images of plasma membrane sheets of undifferentiated 3T3-L1 fibroblasts and 3T3-L1 adipocytes after differentiation showing many caveolae invaginations. (D) Immunofluorescence-staining against caveolin1 and cavin1 in 3T3-L1 adipocyte plasma membrane (PM) sheets (magenta) illustrate endogenous level of both proteins. (E) Representative confocal images of 3T3-L1 adipocyte immuno-fluorescence stained against caveolin1 (magenta) and lipid droplets (green).

**Figure S9:**
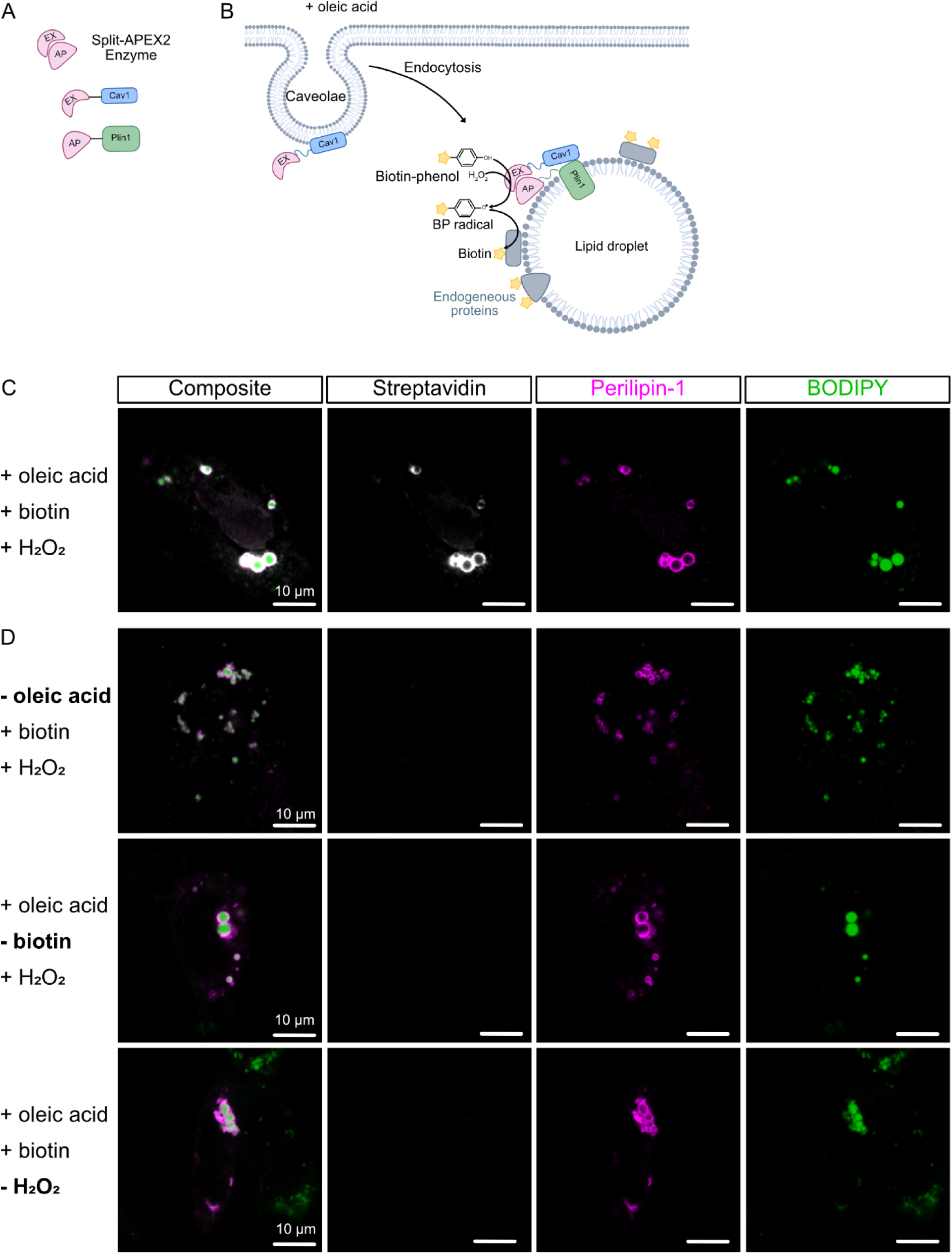
SplitAPEX2 Caveolin1- and Perilipin1- constructs showed oleic acid specific biotinylation at lipid droplets. (A-B) Scheme illustrates splitAPEX2 strategy for experimental design accordingly to Han et al. (2019)^63^. (C) Representative confocal image of MEF cells transfected with both Caveolin1-EX and Perilipin1-AP and treated with oleic acid for 3h. Cells were fixed and stained with Streptavidin-AF647 (grey), perilipin1 primary and Rb IgG (H+L) Alexa fluor 594 conjugated secondary antibody (magenta) and lipid droplet stain (BODIPY, green). Scale bars are 10 µm. (D) Control conditions omitting one of the essential components needed for APEX2 reaction (oleic acid, biotin or H2O2). Scale bars are 10 µm.

**Figure S10:**
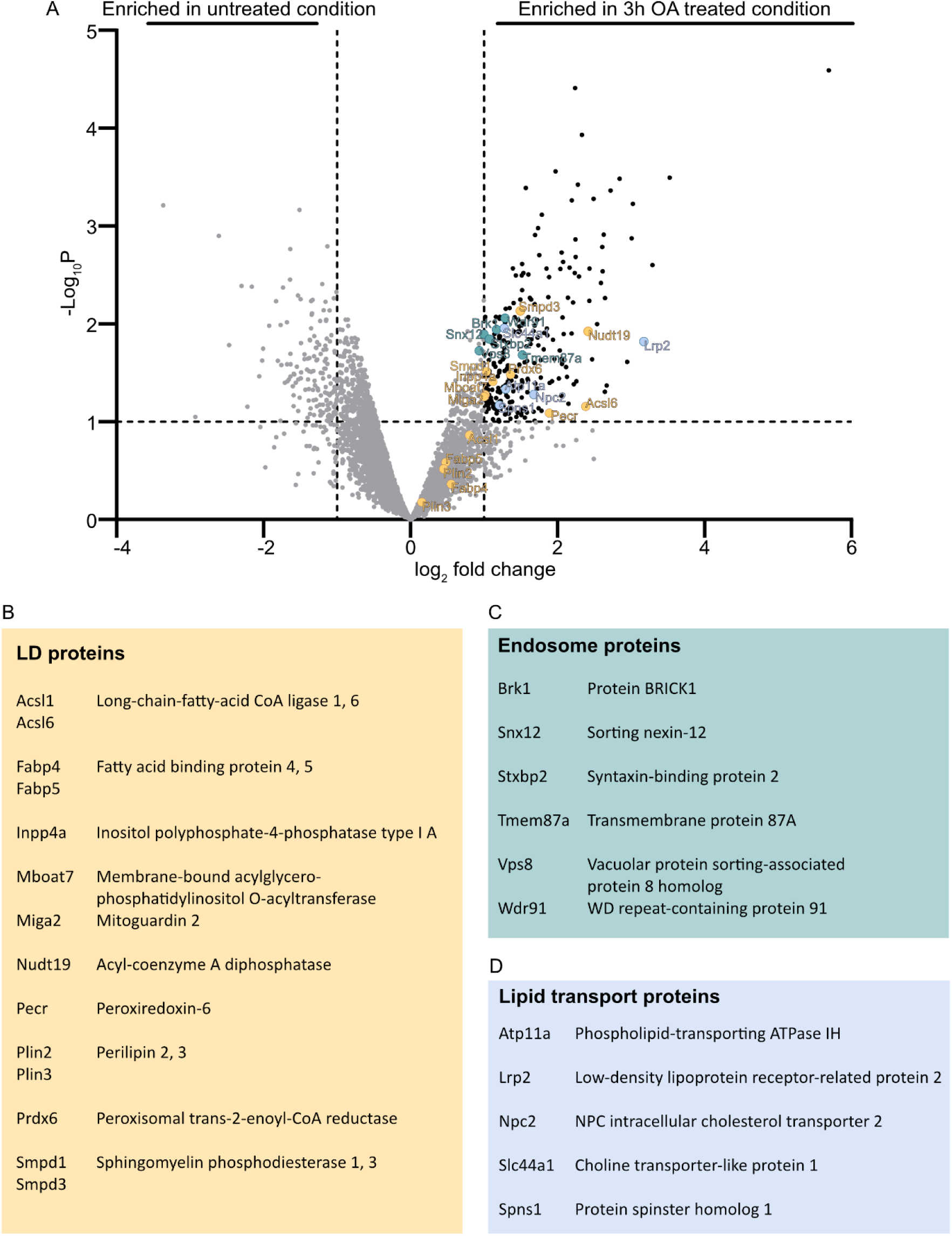
Proteomics of caveolin1 positive lipid droplets. A) MEFs were transfected with caveolin1-EX and perilipin1-AP, after which APEX2 reaction was carried out. Protein lysate samples were collected and biotinylated proteins were enriched via streptavidin beads. Eluates were subjected to mass spectrometry analysis (see detailed description in results and methods part). Volcano plot depicting log2 Fold changes and -log10 p-values. Proteins with a log2FC > 1 and -log10 p-value > 1 were considered significantly enriched when treated for 3 h with oleic acid followed by APEX reaction. B-D) Proteins of interest categorized by their function (retrieved from UniProt).

**Figure S11:**
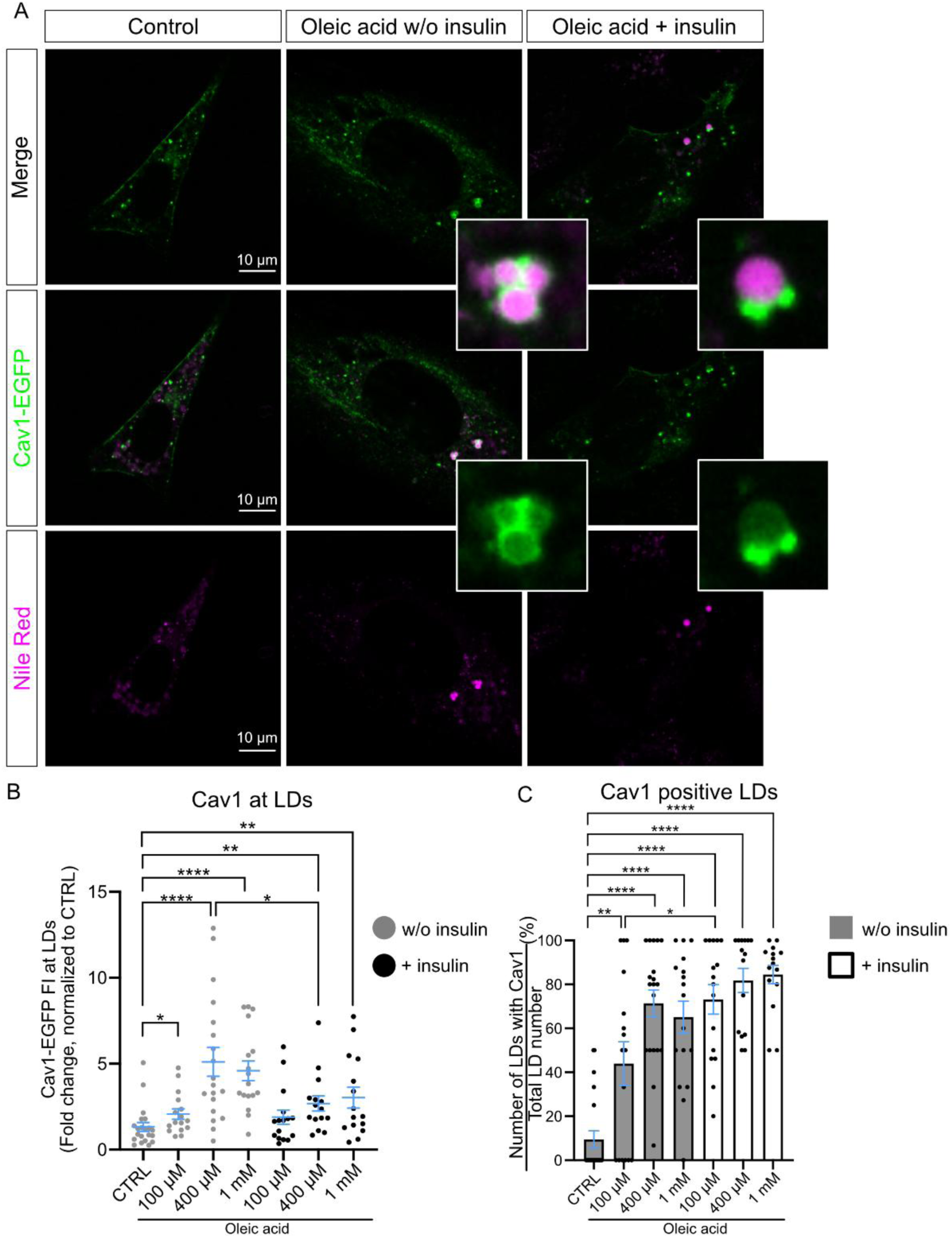
Caveolin1- lipid droplet trafficking is independent of insulin treatment. (A) Representative confocal images of MEFs transfected with caveolin1-EGFP (Cav1-EGFP, green), treated with 400 µM oleic acid for 3h in absence (-) or presence (+) of 1µg/ml insulin. Lipid droplets were stained with Nile Red (magenta). Scale bars are 10 µm. (B) Plot depicts normalized caveolin1-EGFP fluorescence intensity (FI) measured at lipid droplets, each spot represents averaged Cav1 FI per cell (n(CTRL) = 73 LDs/22 cells, n(100µM OA) = 54/16, n(400 µM OA) = 150/19, n(1 mM OA) = 113/17; + insulin: n(100µM OA) = 135/16, n(400 µM OA) = 137/15, n(1 mM OA) = 246/15). (C) Percentage of Cav1 positive lipid droplets per cell, each spot represents a cell (n(CTRL) = 22, n(100µM OA) =16, n(400 µM OA) = 19, n(1 mM OA) = 17; + insulin: n(100µM OA) = 16, n(400 µM OA) = 15, n(1 mM OA) = 15). For all graphs: mean ± SEM, minimum 5 cells per experiment from 3 independent experiments, tested for significant differences with multiple comparison Kruskal-Wallis test, *p≤0.05; **p≤0.01; ***p≤0.001 and ****p≤0.0001).

**Figure S12:**
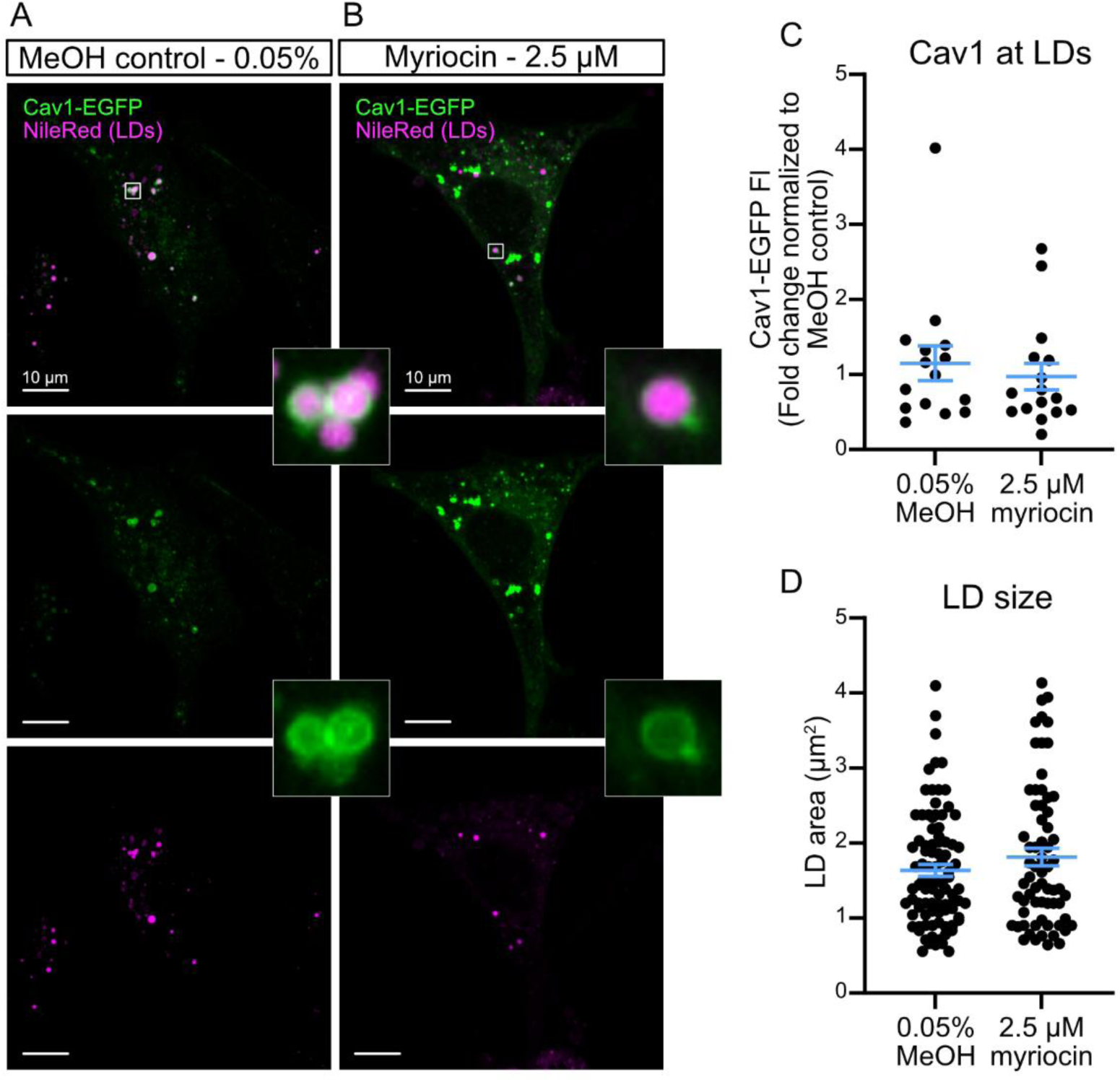
Inhibition of sphingosine biosynthesis does not inhibit caveolin1-lipid droplet trafficking. (A-B) MEF cells were transfected with caveolin1-egfp (green) and treated for 3h with oleic acid and methanol control 0.05% (A) or myriocin 2.5 µM (B) and stained with Nile Red (magenta, lipid droplets). Representative confocal images are depicted, scale bars are 10 µm. (C) Plot shows normalized caveolin1-EGFP fluorescence intensity (FI) measured at lipid droplets, each spot represents averaged fluorescence intensities per cell (n(MeOH) = 15, n(Myriocin) = 16). (D) Lipid droplet (LD) area in µm^2^, each spot represents a single LD (n(MeOH) = 91/15 cells, n(Myriocin) = 66/ 16 cells). For all graphs, mean +/- SEM, 5 cells/experiment, 3 independent experiments, tested for significant differences with Mann-Whitney test to compare methanol control and myriocin treated conditions.

**Figure S13:**
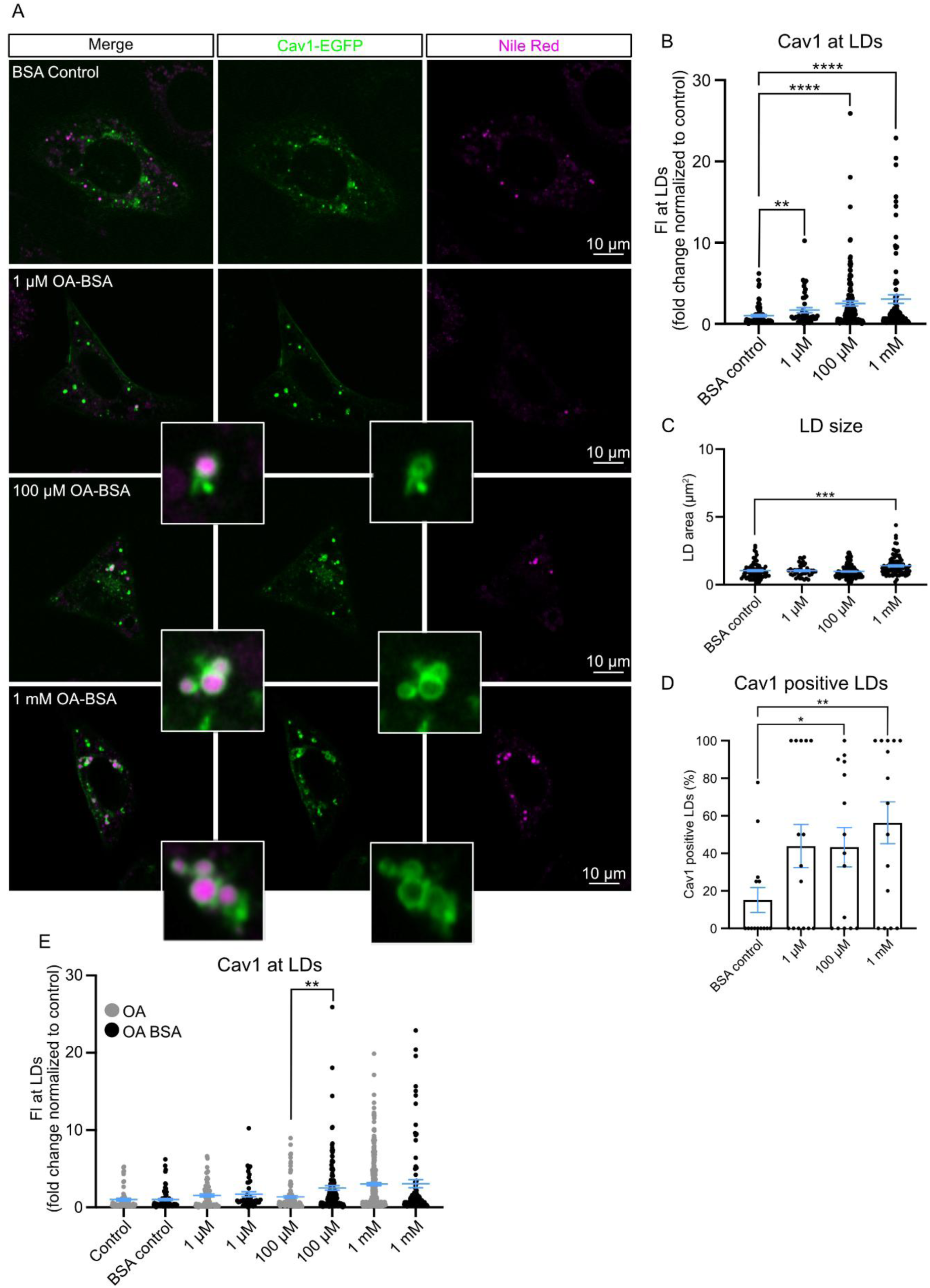
Caveolin1- lipid droplet trafficking in oleic acid vs. oleic acid-BSA treated MEFs revealed no differences in trafficking efficiency. (A) Representative confocal images of MEFs transfected with Caveolin1-EGFP (Cav1-EGFP, green), treated with BSA or oleic acid-BSA complexes (OA-BSA) for 3h. Lipid droplets are stained with Nile Red (magenta). Scale bars are 10 µm (B) Plot depicts normalized caveolin1-EGFP fluorescence intensity (FI) measured at lipid droplets, each spot represents a lipid droplet (n(BSA CTRL) = 72 / 14 cells, n(1 µM OA-BSA) = 43 / 15 cells, n(100 µM OA-BSA) = 133 / 15 cells, n(1 mM OA-BSA) = 87 / 15 cells). (C) Lipid droplet (LD) area in µm^2^, each spot represents a single lipid droplet (n(BSA CTRL) = 72 / 14 cells, n(1 µM OA-BSA) = 43 / 15 cells, n(100 µM OA-BSA) = 133 / 15 cells, n(1 mM OA-BSA) = 87 / 15 cells). (D) Percentage of lipid droplets positive for caveolin1 relative to the total amount of lipid droplets/cell (n(BSA CTRL) = 14, n(1 µM OA-BSA) =15, n(100 µM OA-BSA) = 15, n(1 mM OA-BSA) = 15). (E) Plot depicts normalized caveolin1-EGFP fluorescence intensity (FI) measured at lipid droplets in oleic acid vs. oleic acid-BSA treated MEFs, each spot represents a lipid droplet n(CTRL) = 56 / 14 cells, (n(BSA CTRL) = 72 / 14 cells, n(1 µM OA) = 70, / 13 cells, n(1 µM OA-BSA) = 43 / 15 cells, n(100 µM OA) = 114 / 16 cells, n(100 µM OA-BSA) = 133 / 15 cells, n(1 mM OA) = 285 / 15 cells, n(1 mM OA-BSA) = 87 / 15 cells For all graphs: mean ± SEM, 4-5 cells/experiment, 3 independent experiments, tested for significant differences with multiple comparison Kruskal-Wallis test, *p≤0.05; **p≤0.01; ***p≤0.001 and ****p≤0.0001).

**Figure S14:**
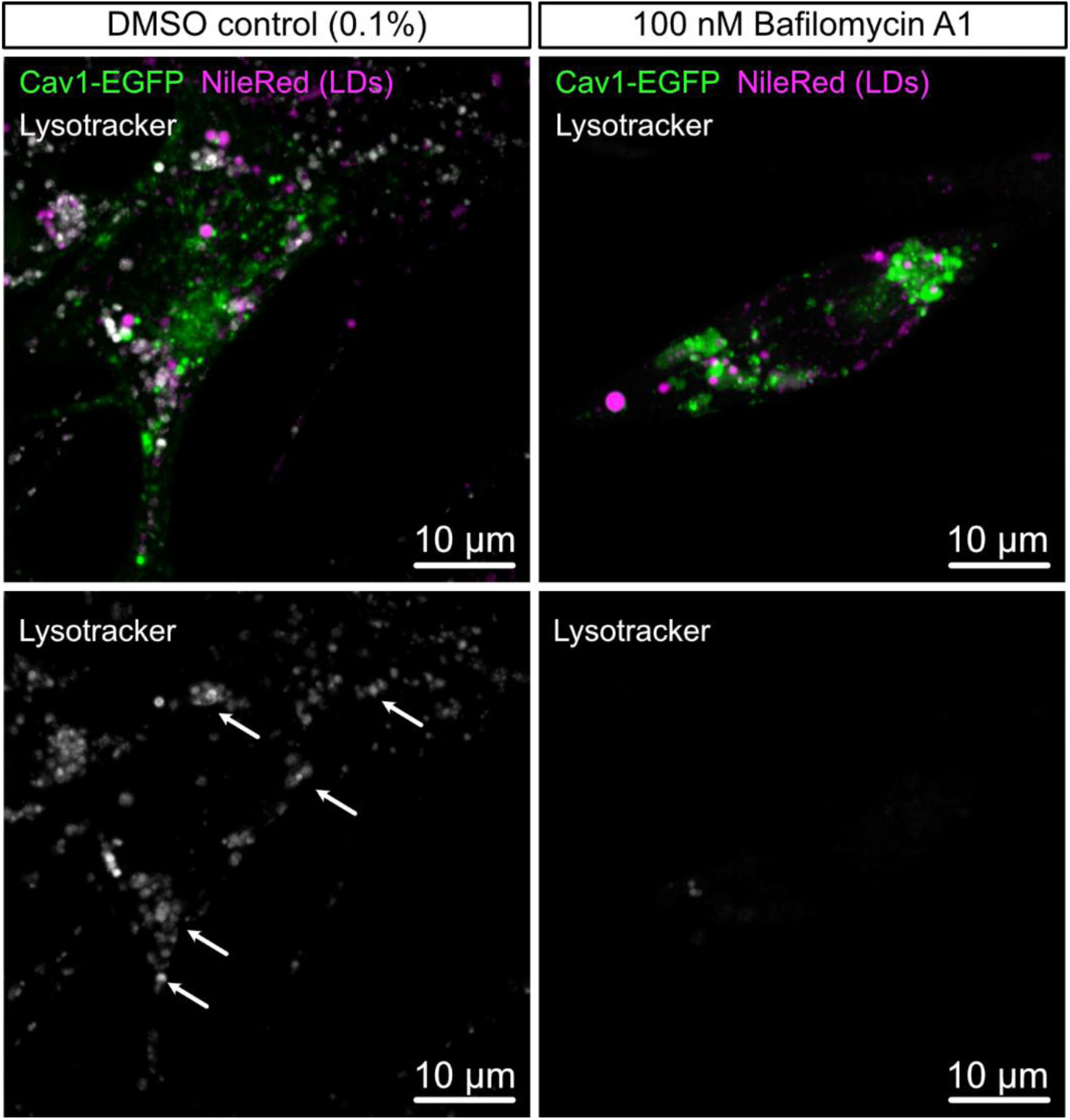
Lysotracker evaluation in MEF expressing caveolin1-EGFP. MEFs expressing caveolin1-EGFP (green) were treated with oleic acid and DMSO (0.1%) or 100 nM Bafilomycin for 3h, followed by lysotracker and Nile Red (magenta) staining for 15 min. Afterwards MEFs were imaged in live-cell imaging buffer. Lysotracker staining (white, indicated by arrows) indicate active lysosomes.

**Figure S15:**
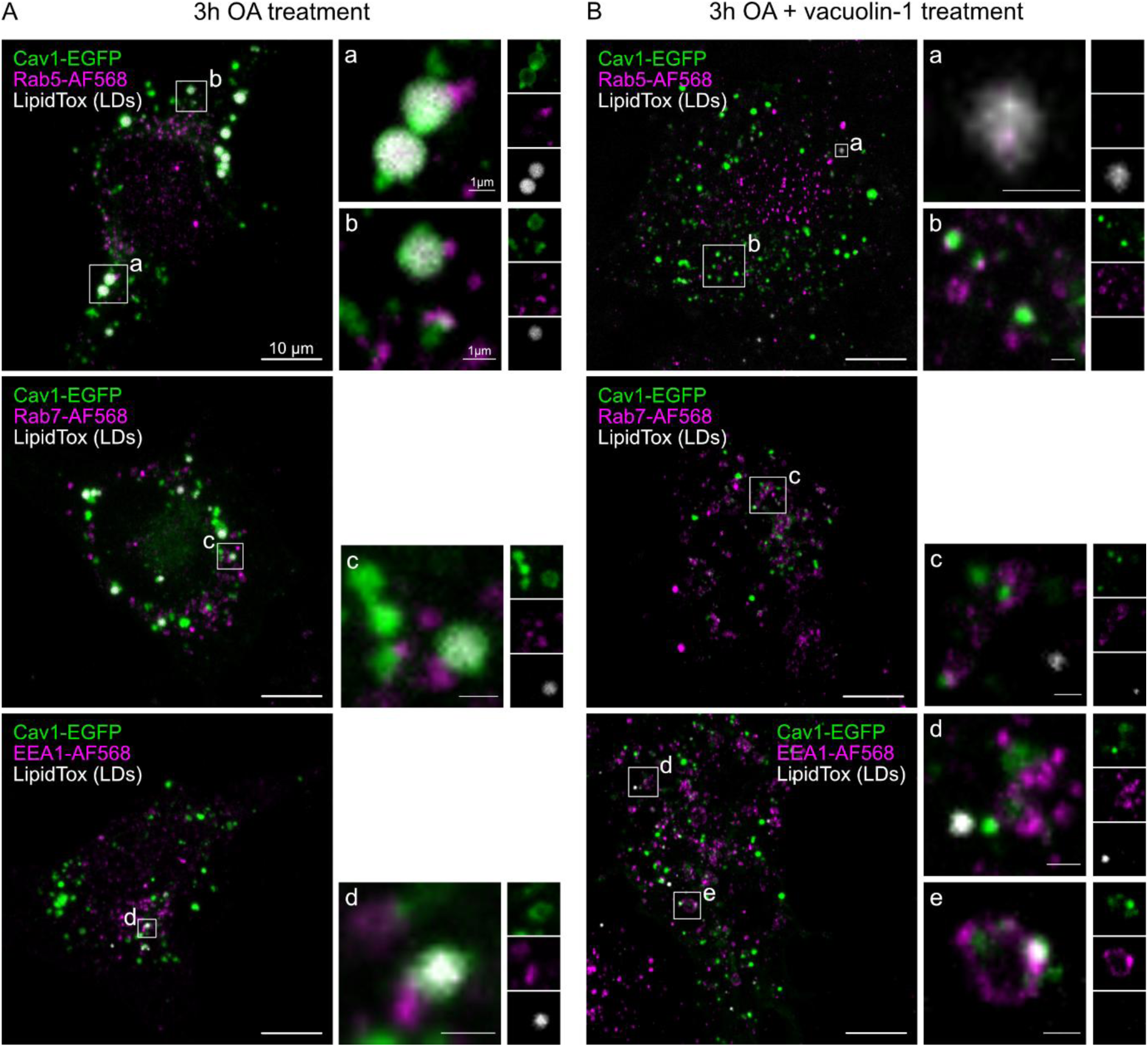
Inhibition of early-to-late endosome maturation results in caveolin1 accumulation in early endosomes. (A-B) Representative confocal images of MEFs transfected with caveolin1-EGFP (green), treated with oleic acid for up to 3h (A) and 1 µM vacuolin-1 (B), followed by immuno-fluorescence staining against EEA1, Rab5 or Rab 7 (magenta) and LipidTox staining (white, lipid droplets, scale bars are 10 µm). Insets represent single confocal planes of acquired z-stack (scale bars are 1 µm).

**Figure S16:**
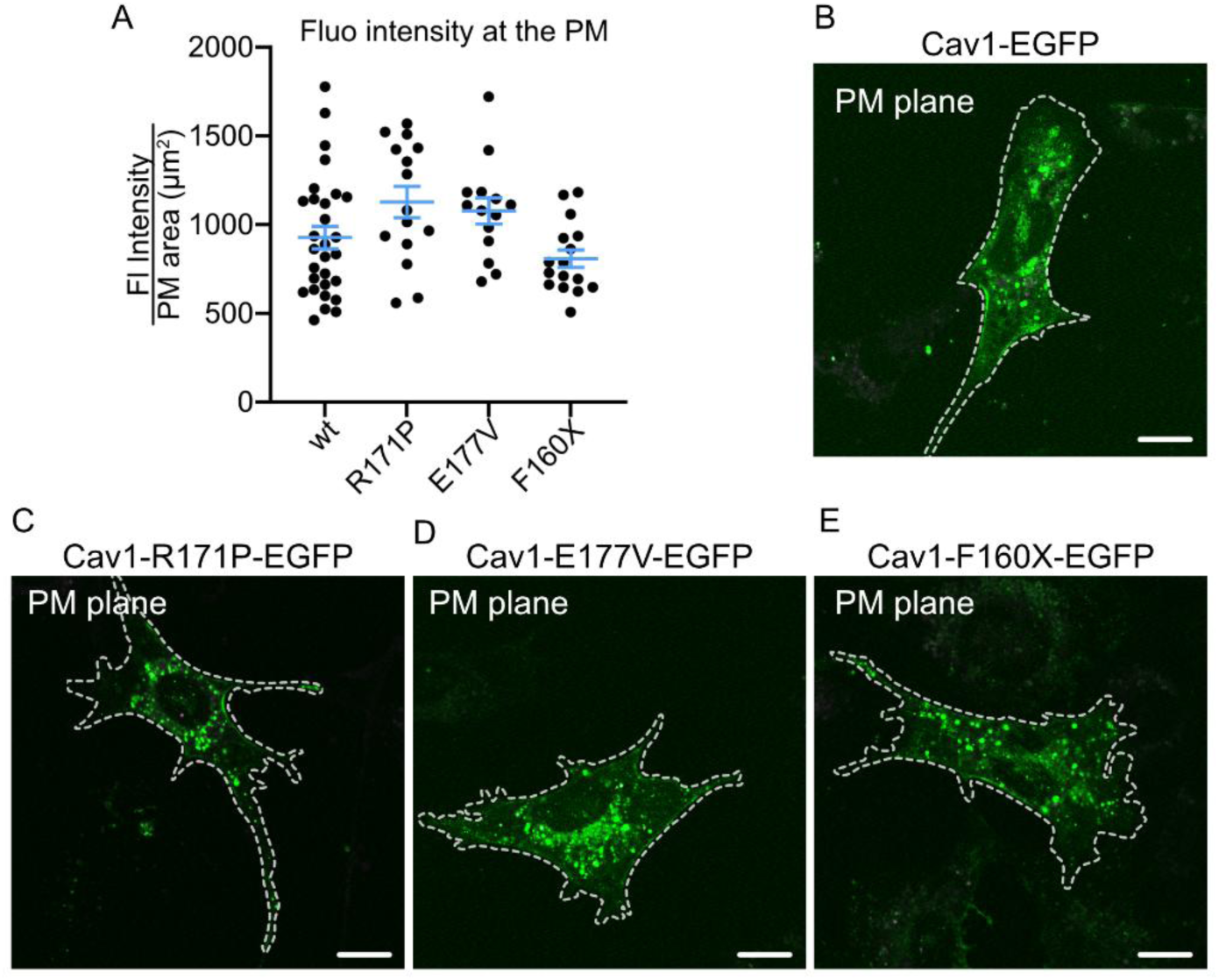
Plasma membrane localization of caveolin1 mutants expressed in MEFs. A) Plot shows normalized caveolin1-EGFP fluorescence intensity (FI, normalized to total cellular PM area) at the plasma membrane (corresponding confocal image stack), each spot represents averaged fluorescence intensities per cell. Graph shows mean ± SEM, minimum 4 cells/experiment, 3 independent experiments, tested for significant differences with one-way ANOVA (n(wt) = 29, n(R171P) =15, n(E177V) = 14, n(F160X) = 16). B-E) Representative confocal images of the plasma membrane plane of cells expressing caveolin1-EGFP wt or mutant plasmids (scale bars are 10 µm). Image acquisition settings were held constant across conditions.

## REFERENCES

1. Matthaeus C et al. The molecular organization of differentially curved caveolae indicates bendable structural units at the plasma membrane. Nat Commun 13, (2022).

2. Parton, R. G. & Del Pozo, M. A. Caveolae as plasma membrane sensors, protectors and organizers. Nat Rev Mol Cell Biol 14, 98–112 (2013).

3. Cheng, J. P. X. & Nichols, B. J. Caveolae : One Function or Many ? Trends Cell Biol 26, 177–189 (2016).

4. Del Pozo, M. A., Lolo, F.-N. & Echarri, A. Caveolae: Mechanosensing and mechanotransduction devices linking membrane trafficking to mechanoadaptation. Curr Opin Cell Biol 68, 113–123 (2021).

5. Cohen, A. W. et al. Caveolin-1-deficient mice show insulin resistance and defective insulin receptor protein expression in adipose tissue. Am J Physiol Cell Physiol 285, 222–235 (2003).

6. Razani, B. et al. Caveolin-1-deficient mice are lean, resistant to diet-induced obesity, and show hypertriglyceridemia with adipocyte abnormalities. Journal of Biological Chemistry 277, 8635–8647 (2002).

7. Liu, L. et al. Deletion of Cavin/PTRF Causes Global Loss of Caveolae, Dyslipidemia, and Glucose Intolerance. Cell Metab 8, 310–317 (2008).

8. Ocket, E. & Matthaeus, C. Insights in caveolae protein structure arrangements and their local lipid environment. Biological Chemistry Preprint at 10.1515/hsz-2024-0046 (2024).

9. Parton, R. G., Tillu, V., McMahon, K. A. & Collins, B. M. Key phases in the formation of caveolae. Curr Opin Cell Biol 71, 7–14 (2021).

10. Parton, R. G., McMahon, K. A. & Wu, Y. Caveolae: Formation, dynamics, and function. Curr Opin Cell Biol 65, 8–16 (2020).

11. Kessels, M. M. & Qualmann, B. The role of membrane-shaping BAR domain proteins in caveolar invagination: from mechanistic insights to pathophysiological consequences. Biochem Soc Trans 48, 137–146 (2020).

12. Parton, R. G., Kozlov, M. M. & Ariotti, N. Caveolae and lipid sorting: Shaping the cellular response to stress. Journal of Cell Biology 219, 1–13 (2020).

13. Örtegren, U. et al. Lipids and glycosphingolipids in caveolae and surrounding plasma membrane of primary rat adipocytes. Eur J Biochem 271, 2028–2036 (2004).

14. Zhou, Y. et al. Caveolin-1 and cavin1 act synergistically to generate a unique lipid environment in caveolae. Journal of Cell Biology 220, (2021).

15. Hubert, M. et al. Lipid accumulation controls the balance between surface connection and scission of caveolae. Elife 9:e55038 (2020) doi:10.7554/eLife.55038.

16. Kenworthy, A. K., Han, B., Ariotti, N. & Parton, R. G. The Role of Membrane Lipids in the Formation and Function of Caveolae. Cold Spring Harb Perspect Biol a041413 (2023) doi:10.1101/cshperspect.a041413.

17. Hubert, M., Larsson, E. & Lundmark, R. Keeping in touch with the membrane; protein-and lipid-mediated confinement of caveolae to the cell surface. Biochem Soc Trans 48, 155–163 (2020).

18. Larsson, E. et al. Lipid packing contributes to the confinement of caveolae to the plasma membrane. (2025) doi:10.7554/eLife.108369.1.

19. Foti, M., Porcheron, G., Fournier, M., Maeder, C. & Carpentier, J. L. The neck of caveolae is a distinct plasma membrane subdomain that concentrates insulin receptors in 3T3-L1 adipocytes. Proc Natl Acad Sci U S A 104, 1242–1247 (2007).

20. Fagerholm, S., Örtegren, U., Karlsson, M., Ruishalme, I. & Strålfors, P. Rapid insulin-dependent endocytosis of the insulin receptor by caveolae in primary adipocytes. PLoS One 4, (2009).

21. Karlsson, M. et al. Colocalization of insulin receptor and insulin receptor substrate-1 to caveolae in primary human adipocytes: Cholesterol depletion blocks insulin signalling for metabolic and mitogenic control. Eur J Biochem 271, 2471–2479 (2004).

22. Kabayama, K. et al. Dissociation of the insulin receptor and caveolin-1 complex by ganglioside GM3 in the state of insulin resistance. PNAS 104, 13678–13683 (2007).

23. Sessa, W. C. eNOS at a glance. J Cell Sci 117, 2427–2429 (2004).

24. Goligorsky, M., Li, H., Brodsky, S. & Chen, J. Relationships between caveolae and eNOS: everything in proximity and the proximity of everything. Am J Physiol Renal Physiol 283, F1–F10 (2002).

25. Parton, R. G. et al. Caveolae: The FAQs. Traffic 21, 181–185 (2020).

26. Matthaeus, C. & Taraska, J. W. Energy and Dynamics of Caveolae Trafficking. Front Cell Dev Biol 8, (2021).

27. Pilch, P. F. & Liu, L. Fat caves: Caveolae, lipid trafficking and lipid metabolism in adipocytes. Trends in Endocrinology and Metabolism 22, 318–324 (2011).

28. Pilch, P. F., Meshulam, T., Ding, S. & Liu, L. Caveolae and lipid trafficking in adipocytes. Clin Lipidol 6, 49–58 (2011).

29. le Lay, S., Briand, N. & Dugail, I. Adipocyte size fluctuation, mechano-active lipid droplets and caveolae. Adipocyte 4, 158–160 (2015).

30. Matthaeus, C. et al. EHD2-mediated restriction of caveolar dynamics regulates cellular fatty acid uptake. Proceedings of the National Academy of Sciences 117, 7471–7481 (2020).

31. Ring, A., Le Lay, S., Pohl, J., Verkade, P. & Stremmel, W. Caveolin-1 is required for fatty acid translocase (FAT/CD36) localization and function at the plasma membrane of mouse embryonic fibroblasts. Biochim Biophys Acta Mol Cell Biol Lipids 1761, 416–423 (2006).

32. Pohl, J. et al. Long-Chain Fatty Acid Uptake into Adipocytes Depends on Lipid Raft Function. Biochemistry 43, 4179–4187 (2004).

33. Hao, J. W. et al. CD36 facilitates fatty acid uptake by dynamic palmitoylation-regulated endocytosis. Nat Commun 11, 1–16 (2020).

34. Stremmel, W., Pohl, J., Ring, A. & Herrmann, T. A new concept of cellular uptake and intracellular trafficking of long-chain fatty acids. Lipids 36, 981–989 (2001).

35. Pohl, J. Uptake of long-chain fatty acids in HepG2 cells involves caveolae: analysis of a novel pathway. The Journal of Lipid Research 43, 1390–1399 (2002).

36. Fernandez, M. A. et al. Caveolin-1 Is Essential for Liver Regeneration. Science(1979) 96, 1628–1632 (2006).

37. Briand, N. et al. Caveolin-1 expression and cavin stability regulate caveolae dynamics in adipocyte lipid store fluctuation. Diabetes 63, 4032–4044 (2014).

38. Craveiro Sarmento, A. S., et al. The worldwide mutational landscape of Berardinelli-Seip congenital lipodystrophy. *Mutation Research - Reviews in Mutation Research* vol. 781 30–52 Preprint at 10.1016/j.mrrev.2019.03.005 (2019).

39. Han, B. et al. Characterization of a caveolin-1 mutation associated with both pulmonary arterial hypertension and congenital generalized lipodystrophy. Traffic 17, 1297–1312 (2016).

40. Garg, A., Kircher, M., del Campo, M., Amato, R. S. & Agarwal, A. K. Whole exome sequencing identifies de novo heterozygous CAV1 mutations associated with a novel neonatal onset lipodystrophy syndrome. Am J Med Genet A 167, 1796–1806 (2015).

41. Schrauwen, I. et al. A frame-shift mutation in CAV1 is associated with a severe neonatal progeroid and lipodystrophy syndrome. PLoS One 10, (2015).

42. Bersuker, K. et al. A Proximity Labeling Strategy Provides Insights into the Composition and Dynamics of Lipid Droplet Proteomes. Dev Cell 44, 97–112.e7 (2018).

43. Klingelhuber, F. et al. A spatiotemporal proteomic map of human adipogenesis. Nat Metab 6, 861–879 (2024).

44. Carpentier, M. et al. Seipin Governs caveolin-1 trafficking through modulating sphingolipid-glycerolipid balance. Cell Rep 44, 116320 (2025).

45. Pol, A. et al. A caveolin dominant negative mutant associates with lipid bodies and induces intracellular cholesterol imbalance. Journal of Cell Biology 152, 1057–1070 (2001).

46. Le Lay, S. et al. Cholesterol-induced caveolin targeting to lipid droplets in adipocytes: A role for caveolar endocytosis. Traffic 7, 549–561 (2006).

47. Blouin, C. M. et al. Lipid droplet analysis in caveolin-deficient adipocytes: alterations in surface phospholipid composition and maturation defects. J Lipid Res 51, 945–956 (2010).

48. Fujimoto, T., Kogo, H., Ishiguro, K., Tauchi, K. & Nomura, R. Caveolin-2 is targeted to lipid droplets, a new ‘membrane domain’ in the cell. Journal of Cell Biology 152, 1079–1085 (2001).

49. Blouin, C. M., Le Lay, S., Lasnier, F., Dugail, I. & Hajduch, E. Regulated association of caveolins to lipid droplets during differentiation of 3T3-L1 adipocytes. Biochem Biophys Res Commun 376, 331–335 (2008).

50. Storey, S. M., McIntosh, A. L., Senthivinayagam, S., Moon, K. C. & Atshaves, B. P. The phospholipid monolayer associated with perilipin-enriched lipid droplets is a highly organized rigid membrane structure. Am J Physiol Endocrinol Metab 301, 991–1003 (2011).

51. Cohen, A. W. et al. Role of Caveolin-1 in the Modulation of Lipolysis and Lipid Droplet Formation. Diabetes 53, 1261–1270 (2004).

52. Pol, A. et al. Cholesterol and Fatty Acids Regulate Dynamic Caveolin Trafficking through the Golgi Complex and between the Cell Surface and Lipid Bodies □ V. Mol Biol Cell 16, 2091–2105 (2005).

53. Stoeber, M. et al. Oligomers of the ATPase EHD2 confine caveolae to the plasma membrane through association with actin. EMBO J 31, 2350–2364 (2012).

54. Morén, B. et al. EHD2 regulates caveolar dynamics via ATP-driven targeting and oligomerization. Mol Biol Cell 23, 1316–29 (2012).

55. Matthaeus, C. et al. The molecular organization of differentially curved caveolae indicates bendable structural units at the plasma membrane. Nat Commun 13, 7234 (2022).

56. Shvets, E., Bitsikas, V., Howard, G., Hansen, C. G. & Nichols, B. J. Dynamic caveolae exclude bulk membrane proteins and are required for sorting of excess glycosphingolipids. Nat Commun 6, (2015).

57. Nichols, B. J. A distinct class of endosome mediates clathrin-independent endocytosis to the Golgi complex. Nat Cell Biol 4, 374–378 (2002).

58. Pelkmans, L., Bürli, T., Zerial, M. & Helenius, A. Caveolin-stabilized membrane domains as multifunctional transport and sorting devices in endocytic membrane traffic. Cell 118, 767–780 (2004).

59. Puchkov, D., Müller, P. M., Lehmann, M. & Matthaeus, C. Analyzing the cellular plasma membrane by fast and efficient correlative STED and platinum replica EM. Front Cell Dev Biol 11, (2023).

60. Anderson, R. H. et al. Sterols lower energetic barriers of membrane bending and fission necessary for efficient clathrin-mediated endocytosis. Cell Rep 37, (2021).

61. Sochacki, K. A. et al. The structure and spontaneous curvature of clathrin lattices at the plasma membrane. Dev Cell 56, 1131–1146.e3 (2021).

62. Brasaemle, D. L., Dolios, G., Shapiro, L. & Wang, R. Proteomic analysis of proteins associated with lipid droplets of basal and lipolytically stimulated 3T3-L1 adipocytes. Journal of Biological Chemistry 279, 46835–46842 (2004).

63. Han, Y. et al. Directed Evolution of Split APEX2 Peroxidase. ACS Chem Biol 14, 619–635 (2019).

64. Hung, V. et al. Spatially resolved proteomic mapping in living cells with the engineered peroxidase APEX2. Nat Protoc 11, 456–475 (2016).

65. Zimnicka, A. M. et al. Src-dependent phosphorylation of caveolin-1 Tyr-14 promotes swelling and release of caveolae. Mol Biol Cell 27, 2090–2106 (2016).

66. Mastick, C. C. & Saltiel, A. R. Insulin-stimulated tyrosine phosphorylation of caveolin is specific for the differentiated adipocyte phenotype in 3T3-L1 cells. Journal of Biological Chemistry 272, 20706–20714 (1997).

67. Kimura, A., Mora, S., Shigematsu, S., Pessin, J. E. & Saltiel, A. R. The insulin receptor catalyzes the tyrosine phosphorylation of caveolin-1. J Biol Chem 277, 30153–30158 (2002).

68. Gupta, A., Lu, D., Balasubramanian, H., Chi, Z. & Wohland, T. Heptanol-mediated phase separation determines phase preference of molecules in live cell membranes. J Lipid Res 63, (2022).

69. Ingólfsson, H. I. & Andersen, O. S. Alcohol’s effects on lipid bilayer properties. Biophys J 101, 847–855 (2011).

70. Schwenk, R. W., Holloway, G. P., Luiken, J. J. F. P., Bonen, A. & Glatz, J. F. C. Fatty acid transport across the cell membrane: Regulation by fatty acid transporters. Prostaglandins Leukot Essent Fatty Acids 82, 149–154 (2010).

71. Hoernke, M. et al. EHD2 restrains dynamics of caveolae by an ATP-dependent, membrane-bound, open conformation. Proc Natl Acad Sci U S A 114, E4360–E4369 (2017).

72. Lafourcade, C., Sobo, K., Kieffer-Jaquinod, S., Garin, J. & van der Goot, F. G. Regulation of the V-ATPase along the endocytic pathway occurs through reversible subunit association and membrane localization. PLoS One 3, (2008).

73. Lu, F. et al. Identification of NPC1 as the target of U18666A, an inhibitor of lysosomal cholesterol export and Ebola infection. Elife 1–16 (2015) doi:10.7554/eLife.12177.001.

74. Maharjan, Y. et al. Intracellular cholesterol transport inhibition Impairs autophagy flux by decreasing autophagosome–lysosome fusion. Cell Communication and Signaling 20, (2022).

75. Ye, Z. et al. Vacuolin-1 inhibits endosomal trafficking and metastasis via CapZβ. Oncogene 40, 1775–1791 (2021).

76. Lu, Y. et al. Vacuolin-1 potently and reversibly inhibits autophagosome-lysosome fusion by activating RAB5A. Autophagy 10, 1895–1905 (2014).

77. Jaber, N. et al. Vps34 regulates Rab7 and late endocytic trafficking through recruitment of the GTPase-activating protein Armus. J Cell Sci 129, 4424–4435 (2016).

78. Ronan, B. et al. A highly potent and selective Vps34 inhibitor alters vesicle trafficking and autophagy. Nat Chem Biol 10, 1013–1019 (2014).

79. Huotari, J. & Helenius, A. Endosome maturation. EMBO Journal 30, 3481–3500 (2011).

80. Barbieri, M. A., Li, G., Mayorga, L. S. & Stahl, P. D. Characterization of Rab5:Q79L-Stimulated Endosome Fusion 1 The Receptor-Mediated Internalization and Transport of Macromolecules from the Cell Surface into Intracellu. ARCHIVES OF BIOCHEMISTRY AND BIOPHYSICS vol. 326 (1996).

81. Hoffenberg, S. et al. Biochemical and functional characterization of a recombinant GTPase, Rab5, and two of its mutants. Journal of Biological Chemistry 270, 5048–5056 (1995).

82. Porta, J. C. et al. Molecular architecture of the human caveolin-1 complex. Sci Adv 8, eabn7232 (2022).

83. Han, B. et al. Characterization of a caveolin-1 mutation associated with both pulmonary arterial hypertension and congenital generalized lipodystrophy. Traffic 17, 1297–1312 (2016).

84. Le Lay, S., Magré, J. & Prieur, X. Not Enough Fat: Mouse Models of Inherited Lipodystrophy. *Frontiers in Endocrinology* vol. 13 Preprint at 10.3389/fendo.2022.785819 (2022).

85. Blouin, C. M. et al. Lipid droplet analysis in caveolin-deficient adipocytes: Alterations in surface phospholipid composition and maturation defects. J Lipid Res 51, 945–956 (2010).

86. le Lay, S. & Kurzchalia, T. v. Getting rid of caveolins: Phenotypes of caveolin-deficient animals. Biochim Biophys Acta Mol Cell Res 1746, 322–333 (2005).

87. Parton, R. G., Taraska, J. W. & Lundmark, R. Is endocytosis by caveolae dependent on dynamin? Nature Reviews Molecular Cell Biology vol. 25 511–512 Preprint at 10.1038/s41580-024-00735-x (2024).

88. Torrino, S. et al. EHD2 is a mechanotransducer connecting caveolae dynamics with gene transcription. Journal of Cell Biology 1–14 (2018).

89. Sinha, B. et al. Cells respond to mechanical stress by rapid disassembly of caveolae. Cell 144, 402–413 (2011).

90. Golani, G., Ariotti, N., Parton, R. G. & Kozlov, M. M. Membrane Curvature and Tension Control the Formation and Collapse of Caveolar Superstructures. Dev Cell 48, 523–538.e4 (2019).

91. Wu, Y. et al. Caveolae sense oxidative stress through membrane lipid peroxidation and cytosolic release of CAVIN1 to regulate NRF2. Dev Cell 58, 376–397.e4 (2023).

92. Wu, Y. et al. Pro-ferroptotic lipids as key control points for caveola formation and disassembly. Cell Rep 44, (2025).

93. Pope, L. E. & Dixon, S. J. Regulation of ferroptosis by lipid metabolism. Trends Cell Biol 33, 1077–1087 (2023).

94. Morén, B. et al. EHD2 regulates adipocyte function and is enriched at cell surface–associated lipid droplets in primary human adipocytes. Mol Biol Cell 30, 1147–1159 (2019).

95. Mundy, D. I., Li, W. P., Luby-Phelps, K. & Anderson, R. G. W. Caveolin targeting to late endosome/lysosomal membranes is induced by perturbations of lysosomal pH and cholesterol content. Mol Biol Cell 23, 864–880 (2012).

96. Hayer, A. et al. Caveolin-1 is ubiquitinated and targeted to intralumenal vesicles in endolysosomes for degradation. Journal of Cell Biology 191, 615–629 (2010).

97. He, K. et al. Internalization of the TGF-β type i receptor into caveolin-1 and EEA1 double-positive early endosomes. Cell Res 25, 738–752 (2015).

98. Kirchner, P., Bug, M. & Meyer, H. Ubiquitination of the n-terminal region of caveolin-1 regulates endosomal sorting by the VCP/p97 AAA-ATPase. Journal of Biological Chemistry 288, 7363–7372 (2013).

99. Liu, P. et al. Rab-regulated interaction of early endosomes with lipid droplets. Biochim Biophys Acta Mol Cell Res 1773, 784–793 (2007).

100. Alonso-Bivou, M., Pol, A. & Lo, H. P. Moving the fat: Emerging roles of rab GTPases in the regulation of lipid droplet contact sites. *Current Opinion in Cell Biology* vol. 93 Preprint at 10.1016/j.ceb.2025.102466 (2025).

101. Parton, R. G., Bosch, M., Steiner, B. & Pol, A. Novel contact sites between lipid droplets, early endosomes, and the endoplasmic reticulum. Journal of Lipid Research vol. 61 1364 Preprint at 10.1194/jlr.ILR120000876 (2020).

102. Peng, W. et al. Endosomal trafficking participates in lipid droplet catabolism to maintain lipid homeostasis. Nature Communications 16, (2025).

103. Martin, S., Driessen, K., Nixon, S. J., Zerial, M. & Parton, R. G. Regulated localization of Rab18 to lipid droplets: Effects of lipolytic stimulation and inhibition of lipid droplet catabolism. Journal of Biological Chemistry 280, 42325–42335 (2005).

104. Boschi, F. et al. Relationship between lipid droplets size and integrated optical density. European Journal of Histochemistry 63, 53–57 (2019).

105. Luckner, M. & Wanner, G. Precise and economic FIB/SEM for CLEM: with 2 nm voxels through mitosis. Histochem Cell Biol 150, 149–170 (2018).

106. Schott, M. B. et al. Ethanol disrupts hepatocellular lipophagy by altering Rab5-centric LD-lysosome trafficking. Hepatol Commun 8, (2024).

107. Brasaemle, D. L. & Wolins, N. E. Isolation of lipid droplets from cells by density gradient centrifugation. Curr Protoc Cell Biol 2016, 3.15.1-3.15.13. (2016).

108. Aik, D. Y. K. & Wohland, T. Microscope alignment using real-time Imaging FCS. Biophys J 121, 2663–2670 (2022).

109. Krieger, J. W. et al. Imaging fluorescence (cross-) correlation spectroscopy in live cells and organisms. Nat Protoc 10, 1948–1974 (2015).

110. Hughes, C. S. et al. Single-pot, solid-phase-enhanced sample preparation for proteomics experiments. Nat Protoc 14, 68–85 (2019).

111. Demichev, V., Messner, C. B., Vernardis, S. I., Lilley, K. S. & Ralser, M. DIA-NN: neural networks and interference correction enable deep proteome coverage in high throughput. Nat Methods 17, 41–44 (2020).

112. Schindelin, J. et al. Fiji: An open-source platform for biological-image analysis. Nature Methods vol. 9 676–682 Preprint at 10.1038/nmeth.2019 (2012).

113. Abramson, J. et al. Accurate structure prediction of biomolecular interactions with AlphaFold 3. Nature 630, 493–500 (2024).

114. Meng, E. C. et al. UCSF ChimeraX: Tools for structure building and analysis. Protein Science 32, (2023).

115. Perez-Riverol, Y. et al. The PRIDE database at 20 years: 2025 update. Nucleic Acids Res 53, D543–D553 (2025).

